# Optimization of Anticodon Edited Transfer RNAs (ACE-tRNAs) Function in Translation for Suppression of Nonsense Mutations

**DOI:** 10.1101/2024.01.11.575297

**Authors:** Joseph J. Porter, Wooree Ko, Emily G. Sorensen, John D. Lueck

## Abstract

Nonsense suppressor tRNAs or AntiCodon-Edited tRNAs (ACE-tRNAs) have long been envisioned as a therapeutic approach to overcome genetic diseases resulting from the introduction of premature stop codons (PTCs). The ACE-tRNA approach for the rescue of PTCs has been hampered by ineffective delivery through available modalities for gene therapy. Here we have screened a series of ACE-tRNA expression cassette sequence libraries containing >1,800 members to optimize ACE-tRNA function and provide a roadmap for optimization in the future. By optimizing PTC suppression efficiency of ACE-tRNAs, we have decreased the amount of ACE-tRNA required by ∼16-fold for the most common cystic fibrosis-causing PTCs.

## INTRODUCTION

Nonsense mutations are single-nucleotide mutations that convert a canonical amino acid codon to one of the three stop codons (UAA, UAG, and UGA). Nonsense mutations introduce a premature termination codon (PTC) generally with almost complete loss of function of the affected protein both due to translation of a truncated protein product and loss of mRNA transcript triggered by nonsense mediated decay (NMD)^1^. With growing access to human genomic sequence data, more than 7,500 nonsense mutations in nearly 1,000 different human genes have been discovered^2^. PTCs account for close to 11% of all described protein variants leading to inherited human disease. Known PTC-associated disease phenotypes include Duchenne muscular dystrophy^3^, inherited retinal disorders^4^, Hurler syndrome^5^, β-thalessemia^6^, Rett syndrome^7^, and cystic fibrosis (CF)^8^.

A number of approaches have been explored to correct PTCs or complement the affected gene^9^ including CRISPR-Cas approaches (prime editing, homologous recombination following double-strand breaks, and in-frame deletions)^10,11^, small molecules (aminoglycosides, synthetic aminoglycosides, oxodiazoles)^12,13^, pseudouridylation^14^, and exon skipping via antisense oligos^15^. The use of nonsense-suppressor tRNAs or anti-codon-edited tRNAs (ACE-tRNAs) for the suppression of PTCs has seen a resurgence in recent years following the development of a library of tRNAs with the anticodons engineered via mutagenesis to suppress UAA, UAG, or UGA PTCs^16^. Further, work by our group and others has demonstrated the safe and efficacious ability of ACE-tRNAs to suppress PTCs in the context of cDNAs and from genuine genomically-encoded PTCs in cell culture and in mice without appreciable readthrough of natural termination codons (NTCs) that terminate translation on every mRNA transcript^17–23^.

We recently demonstrated that ACE-tRNA delivery alone is sufficient to rescue near-WT cystic fibrosis transmembrane conductance regulator (CFTR) transcript abundance and channel function from the endogenous gene harboring common CF PTCs^22^. An important caveat however, is that these conditions were selected such that the cells had on the order of hundreds of ACE-tRNA copies delivered per cell by cDNA transfection. For ACE-tRNAs to realize their potential as a PTC therapeutic, efficient delivery to the affected tissue(s) will need to be resolved. Likely the success of ACE-tRNAs will be dependent on a matrix of conditions including the target tissue and delivery modalities, as well as the PTC suppression efficiency of the ACE-tRNA cargo. To that end we set out to optimize the ACE-tRNA expression cassette sequence to clear the therapeutic threshold for a wider range of nonsense-associated diseases with fewer ACE-tRNAs delivered and provide opportunity for use of a wider range of viral and non-viral delivery technologies. As a specific example, the therapeutic threshold for clinically meaningful rescue of CFTR is 10-30%, use of an ACE-tRNA expression cassette with a 5-fold lower concentration required to reach 30% rescue of CFTR can tolerate a 5-fold less efficient delivery modality, while still maintaining a clinically meaningful level of CF protein rescue.

Natural tRNA biogenesis and function in translation are a complex, multifaceted process^24–53^ (Supplemental Figure 1). Much of the work done to improve nonsense suppressor tRNA (sup-tRNA) function has been examined in the context of genetic code expansion for the incorporation of non-canonical amino acids in response to a nonsense codon^54–56^. These efforts can be broadly classified into categories of approaches to improve sup-tRNA transcription^57–61^ or sup-tRNA function in translation^62–66^. With these efforts in mind, we turned to natural tRNA biogenesis in human cells to pinpoint areas of potential optimization. We aimed to optimize the transcription, processing, stability, and translational elements of several of our best-performing ACE-tRNAs to increase PTC suppression efficiency. Human tRNA sequences targeted in this study and their proposed function are as follows: 1) the tRNA 5’-upstream control element (5’-UCE) is involved in transcription of tRNAs, 2) the 5’- and 3’-flanking sequences of the tRNA are involved in endo- and exonuclease processing of the pre-tRNA to a mature tRNA, 3) the anticodon loop and stems of the tRNA body can be optimized for interactions with the translational apparatus including the aminoacyl-tRNA synthetase (aaRS) for charging with the cognate amino acid, elongation factor 1A (EF-1A) for transport to the ribosome, and the ribosome function in polypeptide elongation.

We screened a series of libraries composed of >1,800 unique ACE-tRNA expression cassette sequences employing our previously reported high-throughput screening (HTS) platform based on a PTC-containing nanoluciferase (NLuc) reporter. For our current best-performing ACE-tRNA^Arg^_UGA_ and ACE-tRNA^Leu^_UGA_ sequences, we have significantly improved their PTC suppression efficiency for the most prevalent CF-causing nonsense mutations.

## RESULTS

### The influence of extragenic ACE-tRNA sequences on nonsense suppressor function

There are a number of aspects of tRNA biogenesis that should be considered when optimizing genetic sequence elements for nonsense suppression in human cells^17^. These genetic sequence elements can be generally grouped into extragenic sequence elements (5’ or 3’ to the mature tRNA sequence) and intragenic sequence elements (contained within the mature tRNA sequence). Extragenic sequence elements include the 5’ upstream control element (5’ UCE), 55 base pairs (bp) immediately upstream of the tRNA sequence, which includes both a degenerate TATA box and the transcription start site (Figure 1A). After transcription the 5’ UCE also determines the sequence of the transcribed 5’ leader sequence, which is removed from the pre-tRNA by nuclease processing. Downstream of the tRNA sequence is the 3’ trailer region, of which the main defined feature is a short poly-thymidine tract (poly-T), sufficient for RNA polymerase III (RNA Pol III) transcription termination. This region is also thought to play a role in transcription re-initiation and contains the sequence of the transcribed 3’ trailer sequence which is removed from the pre-tRNA by nuclease processing. As such, we sought to optimize these sequences for our best-performing ACE-tRNA^Arg^_UGA_, which represents the most common PTC leading to human disease^67^. To test these sequence elements, we employed a similar all-in-one high throughput cloning and screening (HTCS) platform as described previously^16^. As before, this iteration of our HTCS platform contains a PTC-NanoLuc luciferase (PTC-Nluc) to provide a readout of nonsense suppression efficiency resulting from the ACE-tRNA cassette sequence, a high-throughput cloning Golden Gate (GG) cassette with ccdB negative selection marker for ∼100% efficiency cloning of ACE-tRNA sequence elements, with the addition of a firefly luciferase (Fluc) expression cassette for transfection efficiency normalization. These features comprise the HTCS platform we have termed pNanoRePorter 2.0 (Figure 1B).

**Figure 1.**
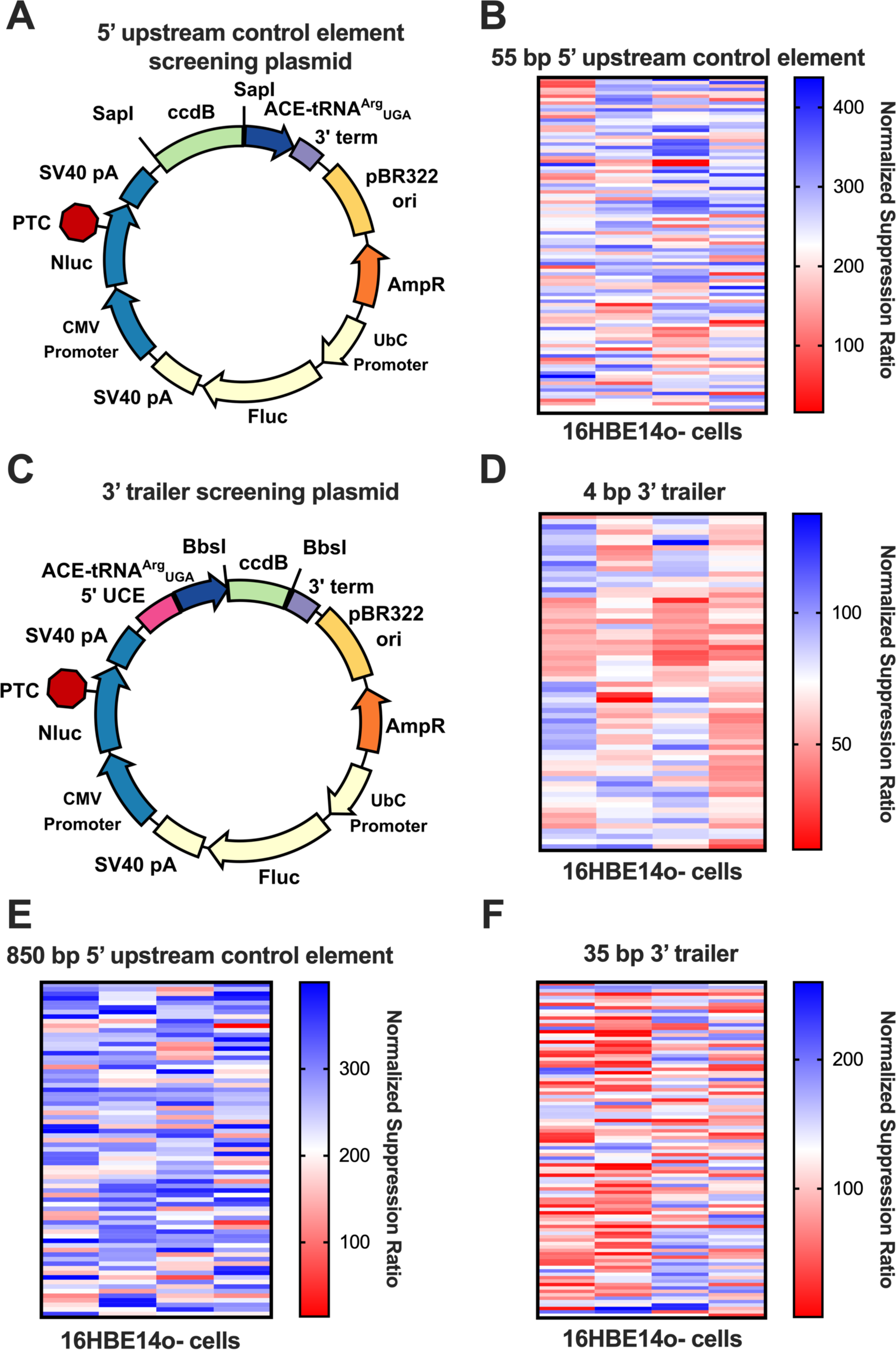
Optimization of ACE-tRNA extragenic sequences. (A) Plasmid map of the high-throughput cloning and screening (HTCS) vector for determining optimal 5’ upstream control element (UCE) sequences for ACE-tRNA nonsense suppressor function. SapI Type IIS restriction enzyme sites flank a negative selection marker (ccdB) for HTCS of 5’ UCE sequences cloned via Golden Gate assembly. As previously reported, the PTC-containing NanoLuciferase expression cassette reports on PTC suppression efficiency of the ACE-tRNA^16^, while the firefly luciferase expression cassette allows for normalization of the signal to transfection efficiency. This dual luciferase reporter is denoted pNanoRePorter 2.0 (B) Heat map representing the results of screening a 386-member library of 55-bp 5’ UCE sequences derived from each unique tRNA gene 5’ UCE in the human genome in 16HBE14o-cells. (C) Plasmid map of the HTCS vector for determining optimal 3’ trailer sequences for ACE-tRNA nonsense suppressor function. BbsI Type IIS restriction enzyme sites flank a negative selection marker (ccdB) for HTCS of 3’ trailer sequences cloned via Golden Gate assembly. (D) Heat map representing the results of screening a 256-member library of 4-bp 3’ trailer sequences representing every 4-bp combination of nucleotides following the ACE-tRNA in 16HBE14o-cells. (E) Heat map representing the results of screening a 292-member library of 850-bp 5’ UCE sequences derived from every synthetically accessible, unique, 5’ UCE in the human genome in 16HBE14o-cells. (F) Heat map representing the results of screening a 389-member library of 35-bp 3’ trailer sequences derived from every unique 35-bp 3’ trailer in the human genome in 16HBE14o-cells. All values displayed in these heat maps represent the average of 6 independent transfections of HTCS library members. The normalized suppression ratio shown here is calculated from the equation (PTC-NanoLuciferase luminescence [+ACE-tRNA]/Firefly luminescence)/(PTC-Nanoluciferase luminescence [no ACE-tRNA]/Firefly luminescence).

We set out to optimize the extragenic ACE-tRNA sequences in a similar manner to the process previously used to determine the ACE-tRNA sequences with the highest PTC-suppression efficiency^16^. We first determined all unique 55-bp 5’ UCE sequences from the human genomic DNA sequences represented in the GtRNAdb (Supplemental Figure 2B)^68^. These 386 sequences were ordered as duplexed oligonucleotides with 4 nucleotide overhangs for GG cloning into the 5’ UCE pNanoReporter 2.0 UGA (Supplemental Figure 2C). Assembled plasmids were transformed into chemically competent NEB 5-alpha *E. coli*, grown in suspension non-clonally, miniprepped, and Sanger sequence verified. The 5’ UCE library members were transfected into the 16HBE14o-human bronchial epithelial (Figure 1C) or HEK293T cell line (Supplemental Figure 2D) in a 96-well format. The normalized nonsense suppression ratio was determined by measuring the Fluc and Nluc luminescence for each well and normalizing the Nluc/Fluc ratio for each member of the 5’ UCE library to the Nluc/Fluc ratio returned when no ACE-tRNA is present. Remarkably, the 5’ UCE screen displayed an ∼4-fold range of nonsense suppression efficiency when paired with ACE-tRNA^Arg^_UGA_, likely due both to differences in transcription and processing of ACE-tRNA^Arg^_UGA_. While the core transcription machinery only requires ∼55-bp of 5’ UCE sequence, we also wanted to test whether longer 5’ UCE sequences may have an altered impact on transcription. To that end, we ordered 292 unique 850-bp synthetically accessible 5’ UCE sequences as double stranded DNA eBlocks and cloned them into the 5’ UCE pNanoRePorter 2.0 ACE-tRNA^Arg^_UGA_ construct as described above. The results from screening the 850-bp 5’ UCE library (Figure 1F) are generally positively correlated with those from the 55-bp 5’ UCE screen (Supplemental Figure 6).

We next sought to optimize the 3’ trailer of the ACE-tRNA^Arg^_UGA_ using a saturation mutagenesis approach by cloning the 256-variant library of the 4-bp sequence immediately 3’ to the tRNA using the 3’ trailer pNanoRePorter 2.0 UGA (Figure 1D). The results of screening the 4-bp 3’ trailer library in 16HBE14o-cells (Figure 1E) and HEK293T cells (Supplemental Figure 2E) revealed sequences with both a positive and negative influence on the nonsense suppression activity, likely through differences in 3’ processing of the pre-tRNA. In designing the 3’ trailer pNanoRePorter 2.0 UGA plasmid, we were surprised to find a negative impact on nonsense suppression efficiency (∼50%) when we altered the sequence context of the poly-U tract from the 5’ UCE HTCS plasmid. With this in mind, we chose to screen a library of all 389 unique 35-bp 3’ trailer sequences from human tRNA genes. When screening this 35-bp 3’ trailer library in 16HBE14o-cells (Figure 1G) we again noted a range of positive and negative influences on nonsense suppression with the most efficient sequences nearly 2-fold higher than the top member of the 4-bp 3’ trailer library. The 35-bp 3’ trailer sequence likely leads to higher levels of ACE-tRNA production through both re-initiation of transcription and nuclease processing of the pre-tRNA. As with the 5’ UCE, we expect that the optimal 35-bp 3’ trailer sequence will be necessary to ensure consistent expression of ACE-tRNA from different therapeutic viral and non-viral vectors.

Mutations resulting in PTCs cause >7,000 distinct genetic diseases, here we chose to focus on PTC-associated CF. Three of the most common CF-causing PTCs are G542X_UGA_ [2.5% of all CF-causing mutations], W1282X_UGA_ [1.2%], and R1162X_UGA_ [0.4%]. We have previously shown we can rescue CFTR containing endogenous genomic PTCs at both the mRNA transcript and protein translational levels^22^. While the W1282X_UGA_ is a UGG (Trp) to UGA mutation, it has been previously shown that W1282L-CFTR mutant retains near-WT CFTR expression and function^69^. As our best-performing ACE-tRNA^Leu^_UGA_ displays higher nonsense suppression efficiency than ACE-tRNA^Trp^_UGA_ we have chosen to also optimize ACE-tRNA^Leu^_UGA_ in this study^16,22^. With rescue of these PTCs in mind, we set out to optimize the intragenic sequences of our best performing ACE-tRNA^Arg^_UGA_, ACE-tRNA^Leu^_UGA_, ACE-tRNA^Gly^_UGA_, and ACE-tRNA^Trp^_UGA_.

### The influence of intragenic ACE-tRNA sequences on nonsense suppressor function

Optimization of nonsense suppressor tRNA sequences has been explored previously, mostly in the context of the incorporation of non-canonical amino acids for genetic code expansion^17^. These optimized nonsense suppressor tRNA sequence elements include increasing stem C-G content^70,71^, optimizing the t-stem^62,63,65,72^, and optimizing the anticodon loop^66,73–75^, among others. Using these previous studies as a roadmap, we designed libraries to increase the C-G content of stems, which we have termed ‘sticky stems’ (Figure 2A, 2C), optimize the t-stem sequence (Figure 2E, Supplemental Figure 5), and optimize the anticodon loop sequence (Figure 2I). The sticky stem libraries for ACE-tRNA^Arg^_UGA_ and ACE-tRNA^Leu^ were designed based on the results of previous screens of ACE-tRNA family members^16^. The sequence of all functional ACE-tRNAs within each of the two tRNA families were aligned to each other and non-conserved sites in stems were chosen as sites to introduce a C-G or G-C pair. The choice of whether the library pair was to be C-G or G-C was made based on which of those two pairs was present at that site in one of the functional ACE-tRNA sequences. The original ACE-tRNA sequences are shown as cloverleaf diagrams, while sticky stem library member pairs are shown next to the library sites (Figure 2A, 2C). Each of these 128-member libraries with every combination of original or library pair at each site was cloned using duplexed oligos for GG assembly into the pNanoRePorter 2.0 UGA ACE-tRNA HTCS vector (Figure 2K) and screened in 16HBE14o-cells as described above. The ACE-tRNA^Arg^_UGA_ sticky stem library did reveal several sequences with ∼2-fold higher nonsense suppression efficiency but only weak trends in the position of library members improving function were observed, specifically those in the acceptor stem were slightly more favored, while those in the anticodon stem were slightly more disfavored (Supplemental Figure 4). In contrast, several members of the ACE-tRNA^Leu^ sticky stem library displayed strong interdependence. For instance, the ACE-tRNA^Leu^_UGA_ variable loop library members in conjunction are entirely disallowed, although each alone are not as deleterious. The G1-C72 was generally a favorable substitution but less so in conjunction with the G4-C69 pair (Supplemental Figure 4). The most active member of the ACE-tRNA^Leu^_UGA_ sticky stem library displayed ∼7-fold higher nonsense suppression efficiency than the original sequence.

**Figure 2.**
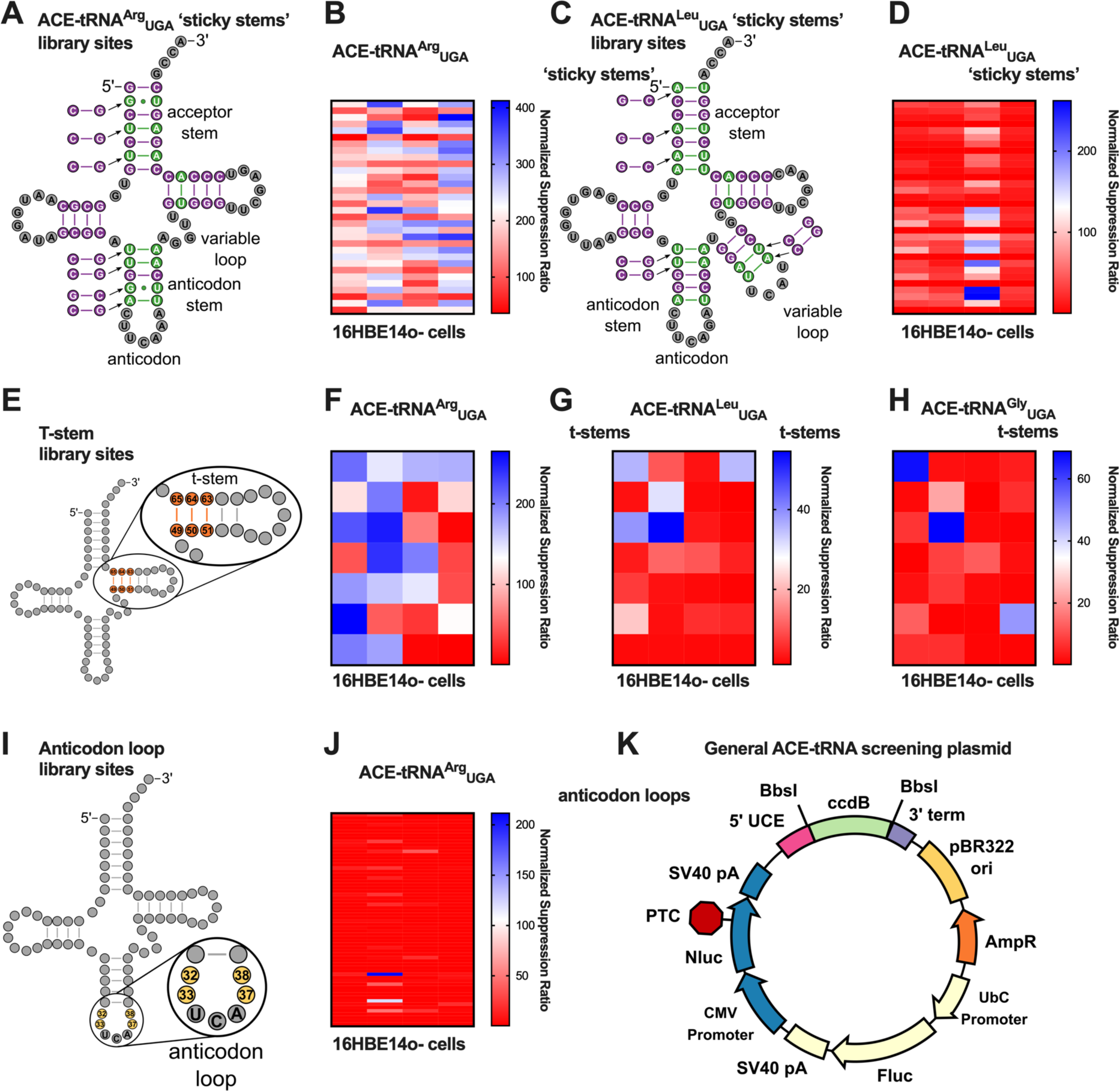
Optimization of ACE-tRNA intragenic sequences. (A) Diagram indicating sites of ‘sticky stem’ mutations for ACE-tRNA^Arg^_UGA_. The ‘sticky stem’ library is composed of every combination of each of the original base pairs indicated in the tRNA cloverleaf or the C-G or G-C pair indicated to the side. (B) Heat map representing the results of screening the 128-member ACE-tRNA^Arg^_UGA_ ‘sticky stem’ library in 16HBE14o-cells. (C) Diagram indicating sites of ‘sticky stem’ mutations for ACE-tRNA^Leu^_UGA_. (D) Heat map representing the results of screening the 128-member ACE-tRNA^Arg^_UGA_ ‘sticky stem’ library in 16HBE14o-cells. (E) Diagram indicating the sites of t-stem mutations for the ACE-tRNA t-stem libraries. The t-stem mutations in these libraries were derived from^45^ as they represent a range of affinities for EF1a. Heat maps representing the results of screening the 28-member ACE-tRNA^Arg^_UGA_ (F), ACE-tRNA^Leu^_UGA_ (G), and ACE-tRNA^Gly^_UGA_ (H) t-stem libraries in 16HBE14o-cells. (I) Diagram indicating the sites of mutations for the ACE-tRNA^Arg^_UGA_ anticodon loop library. Each of these sites was mutated to every combination of the 4 nucleotides. (J) Heat map representing the results of screening the 256-member ACE-tRNA^Arg^_UGA_ anticodon loop library in 16HBE14o-cells. (K) Plasmid map of the high-throughput cloning and screening (HTCS) vector for determining optimal intragenic sequences for ACE-tRNA nonsense suppressor function. BbsI Type IIS restriction enzyme sites flank a negative selection marker (ccdB) for HTCS of ACE-tRNA sequences cloned via Golden Gate assembly. All values displayed in these heat maps represent the average of 6 independent transfections of HTCS library members. The normalized suppression ratio shown here is calculated from the equation (PTC-NanoLuciferase luminescence [+ACE-tRNA]/Firefly luminescence)/(PTC-Nanoluciferase luminescence [no ACE-tRNA]/Firefly luminescence).

We next turned to optimization of the t-stem sequence for each of ACE-tRNA^Arg^_UGA_, ACE-tRNA^Leu^_UGA_, and ACE-tRNA^Gly^_UGA_. Elongation factors in bacteria (EF-Tu) and in eukaryotes (EF1A) have a primary and critical function to shuttle aminoacylated tRNAs to the ribosome for participation in protein translation^43^. It has been previously shown that both the esterified amino acid and tRNA body both contribute to the free energy of formation of the EF1A-tRNA complex, with thermodynamic compensation of both interaction facets resulting in uniform release of aminoacylated tRNAs at the ribosomal A site^45^. To optimize this interaction, we screened a 28-member library of t-stem sequences which cover a wide range of binding affinities (Supplemental Figure 5)^45^. For all ACE-tRNAs tested, C49-G65, C50-G64, G51-C63 representing a medium/high strength t-stem (TS-9, Supplemental Figure 5) performed the best (Figure 2F-H). Several t-stems showed a slightly improved function for ACE-tRNA^Arg^_UGA_, while one showed improved function for ACE-tRNA^Leu^_UGA_. The only t-stem mutant for ACE-tRNA^Gly^_UGA_, which retained any nonsense suppression activity had the same sequence as the original ACE-tRNA^Gly^_UGA_ t-stem sequence.

The last library we assessed for optimizing intragenic ACE-tRNA sequences was the anticodon loop of ACE-tRNA^Arg^_UGA_. The rationale for this library was that altering the anticodon likely introduced structural changes to the anticodon loop that may be reverted to a more WT-like state via another compensatory mutation in the anticodon loop. To that end we cloned a library of all 4 nucleotides for all combinations of residues 32, 33, 37, and 38 (Figure 2I). Screening this 256-member library in 16HBE14o-cells demonstrated that the original sequence provided the highest nonsense suppression efficiency with almost all other sequences resulting in almost fully ablated activity. The few other sequences exhibiting function were identified in our previous ACE-tRNA^Arg^_UGA_ screen^16^. In general, ACE-tRNA^Arg^_UGA_ displayed a remarkable level of sequence plasticity, although not in the anticodon loop. In retrospect this was perhaps not unexpected as 3 of the 4 sites of the anticodon loop library are invariable in all human tRNA^Arg^ sequences^76^.

### Generality of optimized sequences for all ACE-tRNAs

With optimized sequences in hand, we wanted to determine the general applicability of optimized 5’ UCE, 3’ trailer, and t-stem sequences to ACE-tRNAs obtained from our previous screens^16^. To that end, we generated ACE-tRNA expression cassettes as either the original sequence, with the optimized 5’ UCE and 3’ trailer sequences, or swapping the original t-stem for TS-9 into the ACE-tRNA HTCS vector and screened for PTC suppression function in 16HBE14o-cells (Figure 3A-C). The optimized 5’ UCE and 3’ trailer sequences displayed variable responses with 11/19 ACE-tRNAs displaying up to a 3-fold increase in nonsense suppression efficiency, with the other ACE-tRNAs displaying similar suppression efficiency as to the original 5’ UCE and 3’ trailer sequences. This variability in response suggests that there exists an interplay between each ACE-tRNA body sequence and the efficiency of transcription and processing of the 5’ UCE and 3’ trailer sequences. The optimized t-stem screen displayed similar variability of impact on nonsense suppression efficiency with 8/19 ACE-tRNAs displaying up to a 2-fold increase in nonsense suppression efficiency, and 4/19 ACE-tRNAs displaying markedly curtailed nonsense suppression efficiency. The variability of response for optimized sequences for different ACE-tRNAs indicates that while there is some generality of optimized sequences an earnest effort to improve ACE-tRNA suppression efficiency for a panel of ACE-tRNAs should include several optimal sequences identified here to account for the proposed interplay between ACE-tRNA sequence elements.

**Figure 3.**
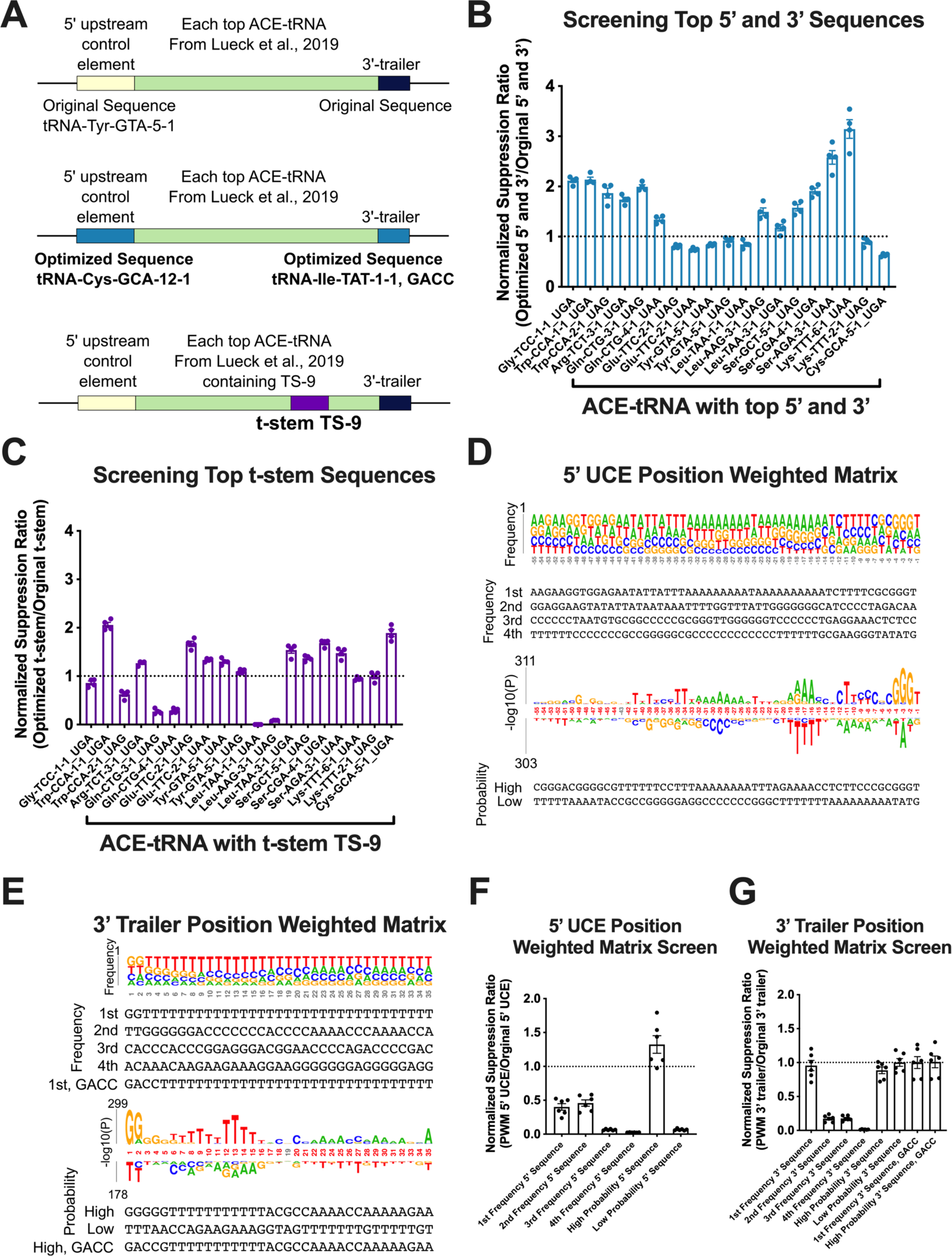
Applicability of optimized sequences for all ACE-tRNAs and determination of general sequences for improved ACE-tRNA function. (A) Diagram of ACE-tRNA sequences to determine the general applicability of the optimized 5’ UCE, 3’ trailer, and t-stem sequences determined from screens shown in Figs. 1 and 2. Each top ACE-tRNA for each isoacceptor/PTC determined previously (Lueck et al., 2019) was tested in concert with either the top 5’ UCE/3’ trailer combination (light blue boxes) or the top t-stem (purple box). (B) Each of the optimized 5’ UCE/3’ trailer sequences was cloned flanking each ACE-tRNA and transfected into HEK293 cells. The data shown here is normalized to the original 5’ UCE/3’ trailer sequences flanking the ACE-tRNA with the dashed line representing the normalized suppression efficiency of the original 5’ UCE/3’ trailer. (C) Each ACE-tRNA containing the top t-stem sequence was cloned and transfected into HEK293 cells. The data shown here is normalized to the original ACE-tRNA t-stem sequence with the dashed line representing the normalized suppression efficiency of the original t-stem. (D and E) Show position weighted matrices (PWM) representing the nonsense suppression efficiency of ACE-tRNA^Arg^_UGA_ paired with each 55-bp 5’ UCE (D) or 35-bp 3’ trailer (E) as either the frequency or probability for each nucleotide. The sequences shown below represent those tested. Each sequence determined from the PWMs was cloned into either the 5’ UCE (F) or 3’ trailer (G) HTCS vector and transfected into HEK293 cells. The data shown here is normalized to the original 5’ UCE or 3’ trailer sequence with the dashed line representing the normalized suppression efficiency of the originals.

### ACE-tRNA 5’ UCE and 3’ trailer screens reveal degenerate consensus motifs

In an effort to find sequence motifs for the 55-bp 5’ UCE and 35-bp 3’trailer libraries that influence nonsense suppression efficiency, we determined the position weight matrix (PWM) for each library. The sequence abundance of each library member was weighted based on the normalized nonsense suppression ratio returned from screening that sequence in 16HBE14o-cells. The frequency and probability logos for the 5’ UCE library sequences (Figure 3D) and 3’ trailer library sequences (Figure 3E) were then determined using *k*pLogo^77^. To experimentally validate these findings, we tested the high and low frequency and probability sequences of the 5’ UCE and 3’ trailer in 16HBE14o-cells (Figure 3F-G). In general, the higher frequency or probability sequences did fare better than the low frequency or probability sequences, however none of them performed significantly better than the original 5’ or 3’ sequences identified through screens of naturally occurring human tRNA elements performed here.

### Optimized ACE-tRNA sequences influence tRNA expression

While reverse transcriptase quantitative PCR (RT-qPCR) provides a facile method to quantify RNA transcript abundance, this basic method is not amenable to quantifying tRNAs due to the presence of on average 8 modified nucleotides per mature tRNA (Figure 4C)^39^, as reverse transcriptase cannot accommodate many of these modifications. To that end, we sought a straightforward method to quantify ACE-tRNA abundance to assay the influence of the optimized sequences on transcription. Here we developed a facile approach to overcome this challenge by appending a self-cleaving ribozyme to the 3’ end of the ACE-tRNA to act as tRNA transcript tabulator (TTT), which can be quantified by RT-qPCR (Figure 4A-C). Because the ribozyme sequence remains consistent throughout tRNA processing and is cleaved from the ACE-tRNA, it is not sensitive to altered ACE-tRNA stability imparted by nucleotide substitution and modifications. We employed the self-cleaving ribozyme drz-Bflo-2 due to its small size (70 bp) and superior self-cleavage kinetics^78^. We first appended the TTT to a series of ACE-tRNA^Arg^_UGA_ cassettes which displayed either high (green), medium (purple) or low (orange) PTC suppression efficiency. While the TTT expression level generally fit the trend of PTC suppression efficiency for the high and low efficiency sequences, the medium PTC suppression sequences displayed a more mixed response (Figure 4D, Supplemental Figure 7). From this, we hypothesize that the mechanisms by which 5’ UCE sequences can influence PTC suppression efficiency are related to transcription and/or 5’ processing of pre-tRNAs (e.g. RNase P). Generally, TTT abundance would be expected to directly vary with pre-tRNA transcription levels, while it would be unlinked to pre-tRNA 5’-processing. With this in mind, we propose the varied TTT response for the medium PTC suppressors is due to this decoupling between the influence of transcription and 5’-processing. When comparing the original 5’ UCE (tRNA-Tyr-GTA-5-1) to the optimized 5’ UCE (tRNA-Cys-GCA-12-1) for each of ACE-tRNAs we again see a mixed response with increases in PTC suppression efficiency for ACE-tRNA^Arg^_UGA_ and ACE-tRNA^Gly^_UGA_ likely due to improvements in 5’-processing, and ACE-tRNA^Leu^_UGA_ and ACE-tRNA^Trp^_UGA_ likely due to increases in transcription. For the intragenic optimized ACE-tRNA^Arg^_UGA_ there was little influence on transcription, while surprisingly intragenic sequences for both ACE-tRNA^Leu^_UGA_, and ACE-tRNA^Trp^_UGA_ led to markedly increased transcription. While we found these results unexpected, these optimized sequences may influence TFIIIC interactions with internal A- and B-box sequences within the tRNA cassettes, leading to increased transcription.

**Figure 4.**
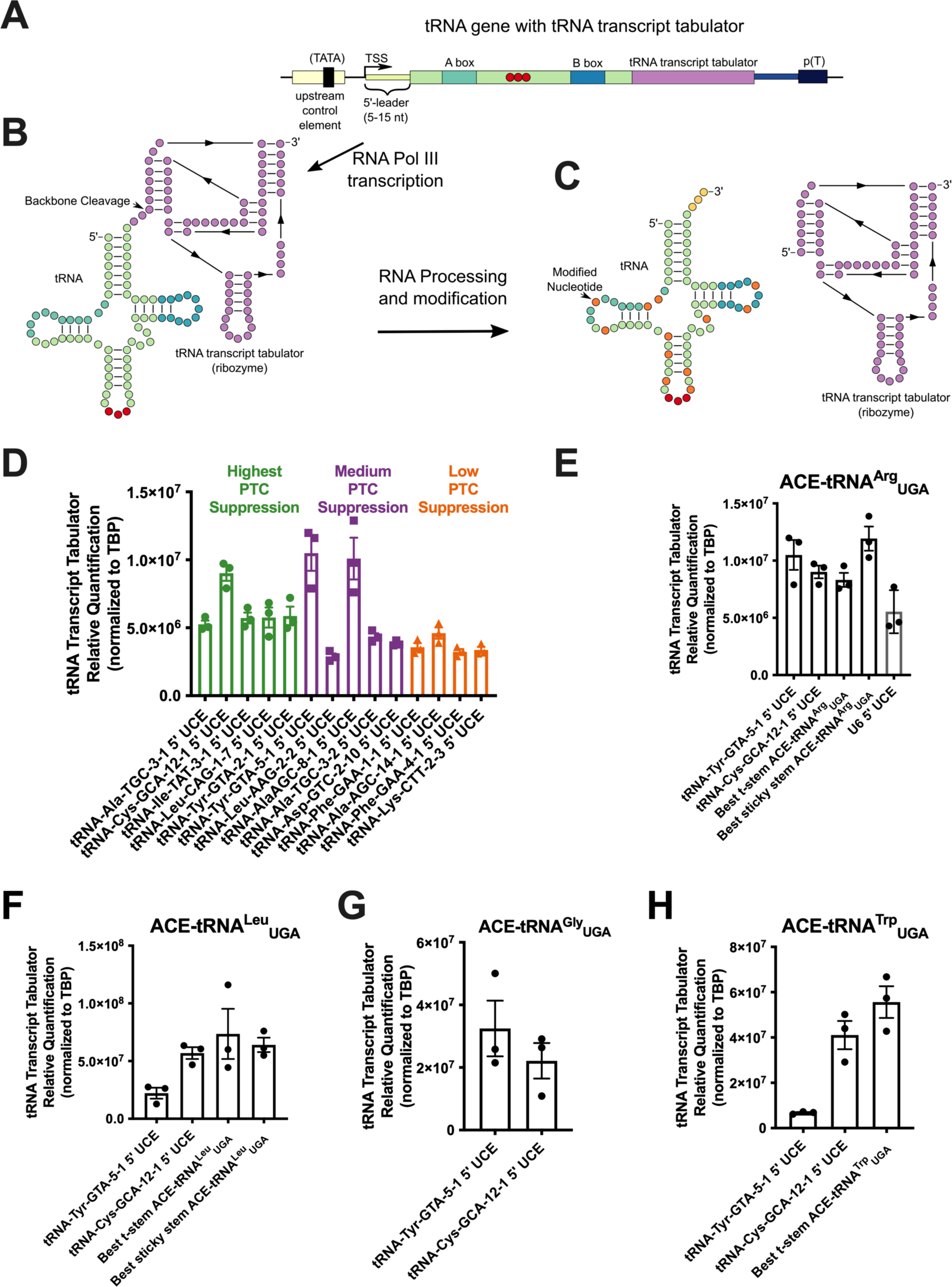
Determination of the steady-state ACE-tRNA expression using the tRNA Transcript Tabulator (TTT) (A) Diagram of a tRNA gene with ribozyme sequence appended (tRNA transcript tabulator, TTT, purple box). (B) RNA Pol III-dependent transcription of the tRNA gene results in production of the pre-tRNA with appended TTT ribozyme (purple circles display a general ribozyme secondary structure). Following transcription, the ribozyme self-cleaves the RNA backbone (indicated by arrow). (C) After tRNA transcription a number of nucleotides are modified (orange circles) which inhibit reverse transcriptase, preventing direct quantification of ACE-tRNAs via RT-qPCR. Instead, the TTT serves as a proxy for ACE-tRNA expression allowing for direct measurement of relative steady-state expression via RT-qPCR. (D) The indicated 5’ UCE sequences representing those which promote high nonsense suppression function when paired with the ACE-tRNA^Arg^_UGA_ (green bars), medium function (purple bars), or low function (orange bars) were paired with ACE-tRNA^Arg^_UGA_ with appended ribozyme TTT. The constructs indicated were transfected into 16HBE14o-cells and the steady-state concentration of TTT was assayed via RT-qPCR. The TTT was appended to several different optimized ACE-tRNA sequences as indicated in the figure for each of ACE-tRNA^Arg^_UGA_ (E), ACE-tRNA^Leu^_UGA_ (F), ACE-tRNA^Gly^_UGA_ (G), and ACE-tRNA^Trp^_UGA_ (H). Each data point represents the average of 3 technical replicates for total RNA isolated from a single transfection with the error bars representing the standard error of the mean.

### Optimized ACE-tRNA expression cassettes require less DNA delivered to rescue equal amounts of PTC-containing protein

While we had separately optimized extragenic and intragenic ACE-tRNA sequence elements we wanted to test them in concert to see if they imparted an additive effect on ACE-tRNA PTC suppression efficiency. To that end we cloned optimized expression cassettes for each of ACE-tRNA^Arg^_UGA_, ACE-tRNA^Leu^_UGA_, ACE-tRNA^Gly^_UGA_, and ACE-tRNA^Trp^_UGA_ (Figure 5A) into a pUC57 mini vector without a reporter for uncoupled delivery of pNanoRePorter 2.0 UGA and ACE-tRNA constructs. Each ACE-tRNA was serially diluted with an empty pUC57 vector and mixed with the pNanoRePorter 2.0 UGA construct, which was held at a constant concentration of 25 ng/μL. The DNA mixes were transfected into HEK293 cells in a 96-well format with the normalized nonsense suppression ratio calculated as outlined above (Figure 5 B-E). To express the DNA concentration dependence of the PTC suppression, we fit the data to the model outlined (Supplemental Figure 8; Supplemental Table 1), with the variables Sup_max_ corresponding to the maximal level of nonsense suppression and DD_50_ ([Delivered DNA]_50_) corresponding to the concentration of DNA in ng/μL required for a half-maximal nonsense suppression response. Improvements were seen for the optimized ACE-tRNAs both in higher Sup_max_ values (∼1.3-fold ACE-tRNA^Arg^_UGA_ and ∼3.5-fold ACE-tRNA^Trp^_UGA_) and in lower DD_50_ values (∼2.3-fold ACE-tRNA^Arg^_UGA_, ∼3.5-fold ACE-tRNA^Leu^_UGA_, and ∼1.4-fold ACE-tRNA^Gly^_UGA_). As a point of reference, this means that ∼10 ng of optimized ACE-tRNA^Arg^_UGA_ will rescue an equivalent amount of PTC-containing protein as 75 ng of the original ACE-tRNA^Arg^_UGA_ expression cassette, with a similar response seen for ACE-tRNA^Leu^_UGA_. We predict the lowered DNA delivery dependence will significantly benefit the function of ACE-tRNAs as therapeutic cargo.

**Figure 5.**
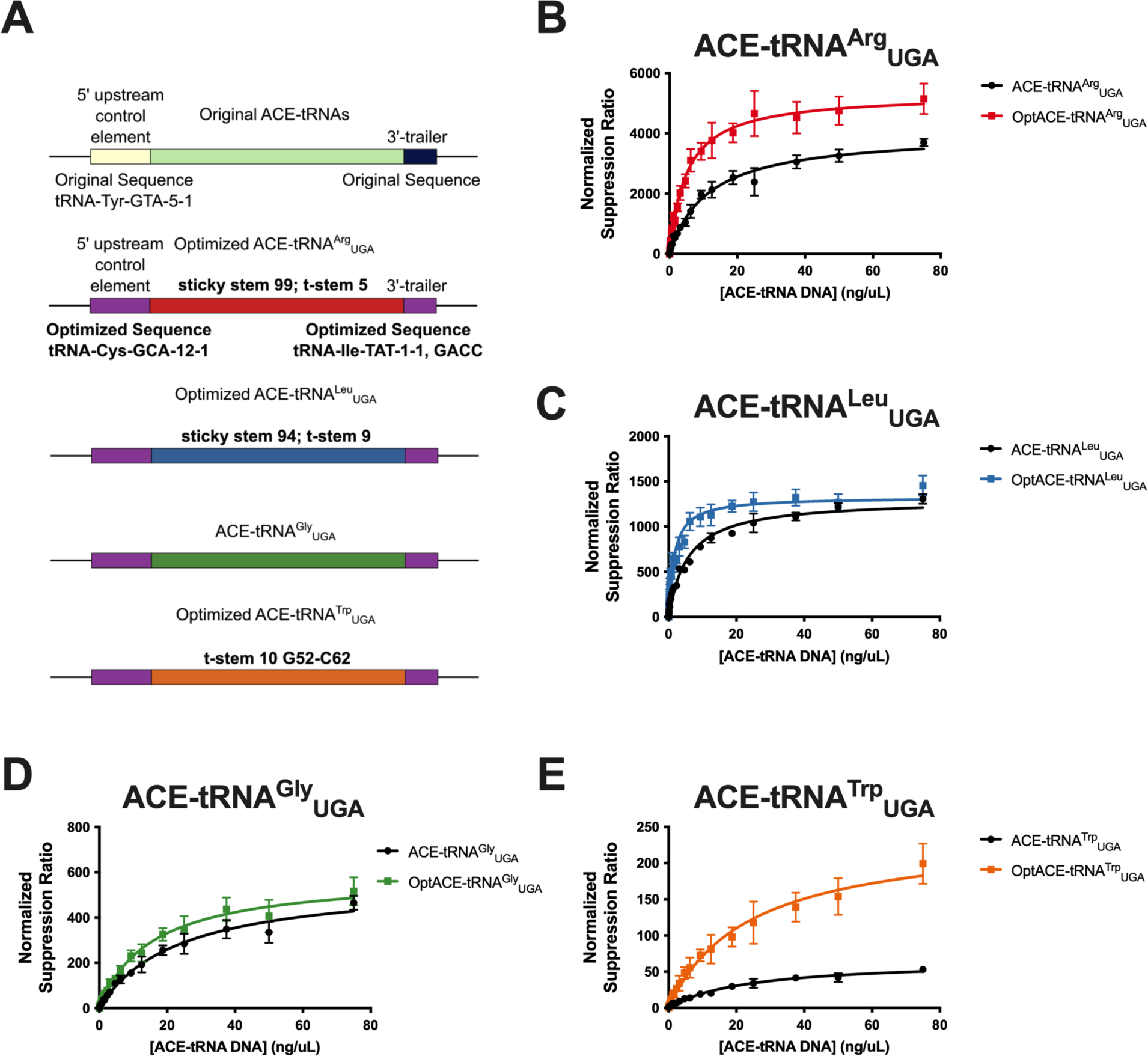
DNA concentration dependence of PTC-protein rescue for optimized ACE-tRNA expression cassettes. (A) Diagram displaying the sequence elements included for each of original or optimized ACE-tRNA^Arg^_UGA_ (red), ACE-tRNA^Leu^_UGA_ (blue), ACE-tRNA^Gly^_UGA_ (green) and ACE-tRNA^Trp^_UGA_ (orange) expression cassettes. In all cases the purple boxes respresent the tRNA-Cys-GCA-12-1 optimized 5’ UCE sequence and the tRNA-Ile-TAT-1-1, GACC optimized 3’ trailer sequence. Other optimized sequences are denoted for each ACE-tRNA. A range of ACE-tRNA concentrations for both original (black data points) and optimized (colored data points) expression cassettes along with a constant amount of pNanoRePorter 2.0 were transfected into HEK293 cells. The concentration dependence of PTC-Nanoluciferase rescue was determined for each of ACE-tRNA^Arg^_UGA_ (red), ACE-tRNA^Leu^_UGA_ (blue), ACE-tRNA^Gly^_UGA_ (green) and ACE-tRNA^Trp^_UGA_ (orange). All values here represent the average of 6 independent transfections. The normalized suppression ratio shown here is calculated from the equation (PTC-NanoLuciferase luminescence [+ACE-tRNA]/Firefly luminescence)/(PTC-Nanoluciferase luminescence [no ACE-tRNA]/Firefly luminescence). The error bars represent the standard error of the mean.

### Optimized ACE-tRNA sequence elements generally do not compromise fidelity in translation

While the optimized ACE-tRNA sequence elements improved nonsense suppression efficiency, we wanted to determine if these improvements come at the expense of tRNA charging fidelity by its cognate tRNA synthetase and subsequent incorporation during translation. While the optimized ACE-tRNAs were predicted by tRNAscan-SE to remain in the same isoacceptor families as their parental tRNA sequences, we sought to demonstrate their fidelity experimentally. We employed the well-characterized superfolder GFP (sfGFP) as a model soluble protein with a UGA codon at amino acid position 150 (p.150). To aid with full-length protein purification we appended a C-terminal Strep-Tag II-8xHistidine-Strep-Tag II series of tags to the sfGFP construct. The sfGFP-pcDNA3.1/Hygro (+) construct and a plasmid expressing 4 copies of ACE-tRNA were co-transfected into HEK293T cells (Figure 6A). Soluble proteins were resolved via SDS-PAGE and sfGFP fluorescence was imaged in the gel, revealing full-length sfGFP expression, with subsequent silver staining of the total protein serving as a loading control (Figure 6B). The full-length sfGFP protein was purified from cell lysates using Strep-Tactin XT resin, resolved by SDS-PAGE, and subsequently stained with Coomassie Blue (Figure 6C). Protein bands representing the full-length protein were excised, subjected to tryptic digest and analyzed by mass spectrometry. All amino acids incorporated during translation at sfGFP p.150 were identified, with >97% incorporation of the cognate amino acid revealed for each of optimized ACE-tRNA^Arg^_UGA_, optimized ACE-tRNA^Leu^_UGA_, and optimized ACE-tRNA^Gly^_UGA_ (Figure 6D). By contrast, we were surprised to find the amino acid incorporated for optimized ACE-tRNA^Trp^_UGA_ was >99.9% Arg, indicating that the mutations made to the anticodon and t-stem of ACE-tRNA^Trp^_UGA_ resulted in misacylation of this tRNA by the arginyl-tRNA synthetase (RARS).

**Figure 6.**
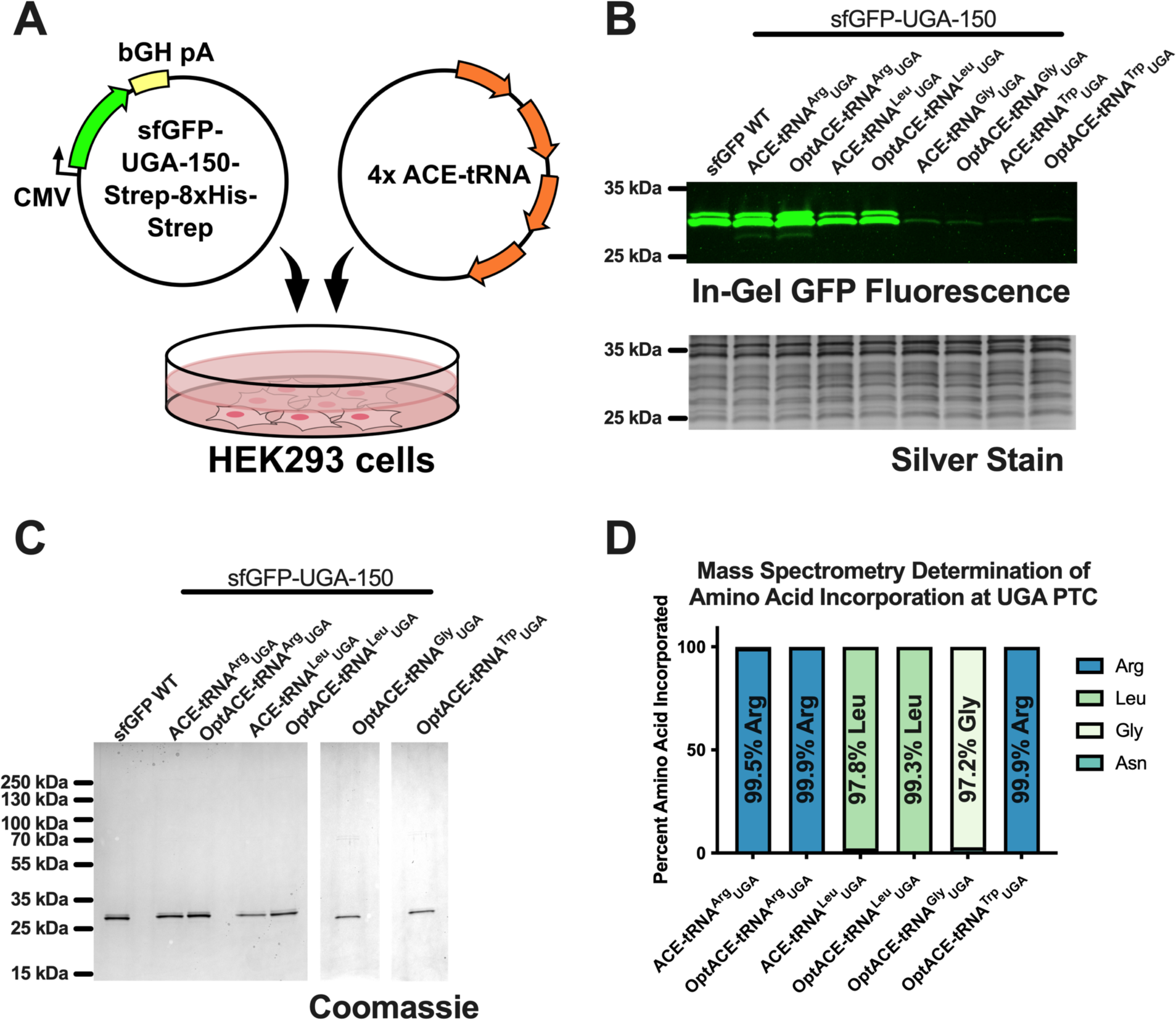
Optimized ACE-tRNA sequences maintain fidelity in translation. (A) A pcDNA3.1/Hygro(+) plasmid expressing superfolder GFP containing a UGA PTC at amino acid position 150 and a C-terminal Strep-8xHis-Strep tag was co-transfected with a plasmid containing 4 copies of either the original or optimized (Opt) ACE-tRNA expression cassette into HEK293 cells. (B) 48 hours following transfection the cells were harvested, lysed via dounce homogenization, a fraction of which was mixed with 6x Laemmli resolving buffer without heating, and resolved on a 10-20% gradient SDS-PAGE gel. The GFP fluorescence was imaged in the gel, with total protein being subsequently assayed via silver stain. (C) The remainder of the soluble cell lysate was subjected to purification using Strep-Tactin XT Superflow resin with the eluted sfGFP protein resolved via SDS-PAGE and the protein visualized with Coomassie stain. (D) The sfGFP protein band was excised from the gel and subjected to trypsin digest followed by mass spectrometric determination of the resulting peptide masses. The mass of each peptide which contained the amino acid at site 150 in sfGFP was determined and the amino acid with the most consistent mass for site 150 in that peptide was determined.

### Optimized ACE-tRNA expression cassettes require less DNA to rescue PTC-containing CFTR protein and mRNA

While the optimized ACE-tRNA sequence elements improved nonsense suppression efficiency for the Nluc reporter protein, we wanted to determine if these improvements translate to other proteins containing PTCs. To that end we introduced 4 of the most common CF-causing PTCs (G542X_UGA_, R553X_UGA_, R1162X_UGA_, and W1282X_UGA_), which represent ∼75% of all CF-causing PTCs, into a construct containing CFTR cDNA with appended C-terminal Nluc expressed from a short ubiquitin C (UbC) promoter (Figure 7A). Each CFTR-PTC-Nluc was co-transfected into HEK293T cells with varying amounts of original or optimized (Opt) ACE-tRNA along with a small amount of plasmid expressing WT sfGFP as a control for transfection efficiency. 48 hours after transfection the cells were lysed in RIPA buffer containing protease inhibitors, 10 μg of total protein was resolved via 10-20% SDS-PAGE gel, GFP fluorescence in the gel was imaged, the gel was subjected to treatment with Nluc substrate (furimazine), and in-gel luminescence was imaged (Figure 7B). Fully glycosylated (C-band) and core glycosylated (B-band) CFTR was evident on the luminescence gel images with C-band intensity normalized to sfGFP intensity used as the measure of full-length CFTR expression. For almost all cDNA amounts tested the OptACE-tRNA cassette was able to rescue significantly more PTC-CFTR than the corresponding ACE-tRNA cassette (Figure 7C). Further, the maximal rescues for each PTC-CFTR as compared to WT (24 ± 4% OptACE-tRNA^Gly^_UGA_/G542X_UGA_, 40 ± 2% OptACE-tRNA^Arg^_UGA_/G542X_UGA_, 58 ± 5% OptACE-tRNA^Arg^_UGA_/R553X_UGA_, 55 ± 3% OptACE-tRNA^Arg^_UGA_/R1162X_UGA_, and 66 ± 5% OptACE-tRNA^Leu^_UGA_/W1282X_UGA_) is well in excess of the 15-30% therapeutic threshold for rescue of CFTR^79^. OptACE-tRNA^Arg^_UGA_ was tested in addition to OptACE-tRNA^Gly^_UGA_ for rescue of G542X_UGA_-CFTR, as G542R-CFTR has previously been shown to retain CFTR channel function^80^, and OptACE-tRNA^Arg^_UGA_ displays higher PTC rescue (Figure 5). Perhaps most strikingly, 16-fold less cDNA is required for OptACE-tRNA^Arg^_UGA_ to rescue the same amount of R553X_UGA_- and R1162X_UGA_-CFTR and >16-fold less cDNA is required for OptACE-tRNA^Leu^_UGA_ to rescue the same amount of W1282X_UGA_-CFTR as compared to their respective original ACE-tRNA expression cassettes. Similar results were seen for rescue of W1282X_UGA_ CFTR PTC mRNA with varying amounts of ACE-tRNA^Leu^_UGA_ and OptACE-tRNA^Leu^_UGA_ cDNA transfected into 16HBEge cells containing the genomically encoded W1282X_UGA_ CFTR variant. When comparing CFTR mRNA rescue exhibited by ACE-tRNA^Leu^ and OptACE-tRNA^Leu^_UGA_, there was no significant difference (p = 0.641) between 2500 ng and 62.5 ng respectively, indicating that ∼40-fold less OptACE-tRNA^Leu^_UGA_ cDNA is required to rescue the same amount of W1282X_UGA_ CFTR mRNA (Figure 7D).

**Figure 7.**
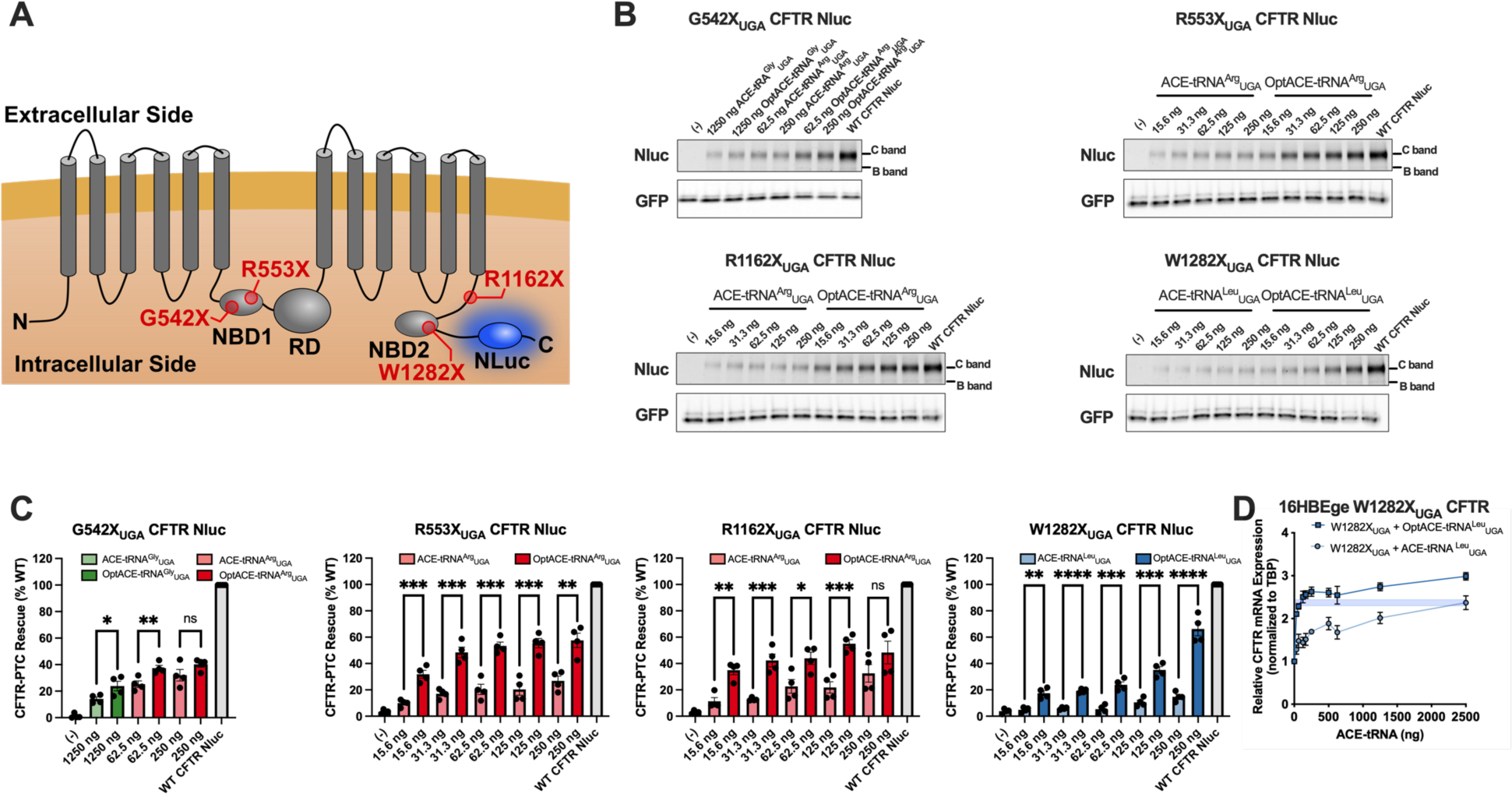
cDNA concentration dependence of PTC-CFTR rescue for optimized ACE-tRNA expression cassettes. (A) CFTR topology diagram displaying the positions of the 4 most common PTC variants found and the C-terminal nanoluciferase tag (NLuc) used for the in-gel luminescence assay. (B) HEK293T cells were co-transfected with original or optimized (Opt) ACE-tRNAs at the amounts shown, along with PTC-containing CFTR variants with a C-terminal translationally fused Nluc tag, and WT sfGFP as a transfection control. 48 hours after transfection the cells were lysed in RIPA buffer containing protease inhibitor cocktail and resolved on a 10-20% SDS-PAGE gel, a GFP fluorescence image was taken, and then the gel was exposed to Nluc luminescence reagent (furimazine) and a luminescence image acquired with representative images shown here. (C) C-band intensity (fully glycosylated CFTR) was determined by densitometry and then normalized to the GFP fluorescence intensity for that sample to normalize for any differences in transfection efficiency. The values plotted represent the normalized C-band luminescence as compared to CFTR-WT-Nluc expressed as a percent for four independent transfections. Solid colored bars represent the optimized ACE-tRNA cassette, while light colored bars represent the original ACE-tRNA cassette, with bars representing the mean and error bars representing the standard error of the mean. Significance was determined by unpaired two-tailed t test where *p < 0.05, **p < 0.01, ***p < 0.001, ****p <0.0001 comparing original and optimized ACE-tRNA cassettes at the same cDNA amount. (D) W1282X_UGA_ CFTR PTC mRNA rescue was assayed by RT-qPCR with varying amounts of ACE-tRNA^Leu^_UGA_ and OptACE^Leu^_UGA_ cDNA transfected.

## DISCUSSION

In transiting the multifaceted biological apparatus leading to translation, each tRNA makes a series of sequential molecular interactions between the tRNA and maturation and modification enzymes, aminoacyl-tRNA synthetases, the elongation factors, and the three ribosomal states of the translation process, among others^81^. The tRNA sequence and its chemical structure by extension should be amenable to each of these interactions, ideally without any sequence element for a particular interaction negatively impacting a separate interaction. With this in mind, when we seek to improve the ACE-tRNA function by altering its expression cassette, we should assay directly for PTC suppression. This allows us to determine the totality of the impact of any specific ACE-tRNA sequence variation on its global function in translation. Further, we performed the PTC suppression assay in human cells, as previous efforts to improve sup-tRNA function for use in human cell culture have demonstrated that improvements made in one domain of life (prokaryotes) were not applicable in another (eukaryotes)^63^.

The sequentiality of tRNA interactions could imply that one specific process in translation represents an interaction bottleneck (the rate-limiting step in enzyme kinetics), however currently there is little evidence to indicate which step this is for ACE-tRNAs. The necessary sequence alterations made to the anticodon of ACE-tRNAs, making them nonsense suppressor tRNAs^16^, suggest that this bottleneck may lie with the ACE-tRNA-aaRS interaction. While alterations to the tRNA can be made to improve this interaction, the application of ACE-tRNAs as a stand-alone PTC therapeutic preclude mutagenesis-based engineering of the aaRS. Another route to tackle a weak ACE-tRNA-aaRS interaction is to simply increase the intracellular pool of mature ACE-tRNA. This increase could arise from increased expression through enhanced transcription, improved post-transcriptional processing efficiency, or extended cellular half-life of the mature ACE-tRNA.

The initial step in transcription of a tRNA gene is modulated by interactions between transcription factor IIIC (TFIIIC) and the internal A- and B-boxes of the tRNA gene^27^, followed by recruitment of TFIIIB, and finally RNA polymerase III (Pol III) which initiates transcription upstream of the tRNA^26^. Following transcription, the pre-tRNA is processed to remove the 5’-RNA leader and 3’-RNA trailer and splicing of an intron if present^39^. As the A- and B-box DNA sequences are retained in the mature tRNA sequence, they cannot be altered without influencing the tRNA function. Therefore, we opted to increase both transcription and 5’-processing by modulating the 55 bp immediately upstream of the tRNA comprising the 5’-upstream control element (5’-UCE). In screening all unique 55 bp 5’-UCEs from the human genome (Fig. 1, Supplemental Fig. 2), we saw a range of responses for the PTC rescue exhibited by ACE-tRNA^Arg^_UGA_ (Fig. 1, Supplemental Fig. 2), likely resulting from impacts of the 5’-UCE sequences on both transcription and 5’ processing of ACE-tRNA^Arg^_UGA_. When considering the steady state abundance of the ACE-tRNA^Arg^_UGA_ assayed with the tRNA transcript tabulator (TTT), we observed a relatively poor correlation between the normalized nonsense suppression ratio and TTT abundance (Supplemental Fig. 7), indicating that some improvements to nonsense suppression were mediated by increased transcription of ACE-tRNA^Arg^_UGA_, while others were likely mediated by enhanced 5’-ACE-tRNA processing. Transplanting the optimal 55 bp 5’-UCE obtained from screening ACE-tRNA^Arg^_UGA_ (tRNA-Cys-GCA-12-1) onto the other extant ACE-tRNAs^16^ demonstrated a range of effects on nonsense suppression efficiency (Fig. 3B). It is known that the binding of pre-tRNA to RNase P can be tuned by the 5’ leader sequence in pre-tRNA in conjunction with the tRNA body^35^, possibly accounting for some of the inconsistencies seen between the 5’-UCE response for different ACE-tRNAs. Nearly half of human tRNAs are silent or poorly expressed^25^, so our results showing that nearly all 55 bp 5’-UCEs promote function of ACE-tRNA^Arg^_UGA_ (Fig. 1, Supplemental Fig. 2) appear inconsistent. However, the expression of genomically encoded human tRNAs is controlled by the regional chromatin state of each tRNA ^82–84^. In this study, the ACE-tRNAs are expressed episomally and therefore it is not expected that the chromatin state would influence expression. As the tRNA expression cassette is not subjected to large scale genomic regulation, it is not surprising that we see consistent function from almost all 55 bp 5’-UCEs in both 16HBE14o- and HEK293T cell lines. For the longer 850 bp 5’-UCE sequences, slightly higher function was seen on average as compared to the 55 bp sequences in the 16HBE14o-cell line, while slightly lower function was seen in the HEK293T cell line for ACE-tRNA^Arg^_UGA_ (Supplemental Fig. 6). This may be explained by differential expression of transcription factors in these cell lines that result in subtle modulation of ACE-tRNA^Arg^_UGA_ transcription from tRNA expression cassettes containing 850 bp 5’-UCEs^85,86^. Important for determining optimal therapeutic ACE-tRNA cargo, significant benefit is not gained by a >55bp 5’ UCE sequence, thus permitting the use of compact ACE-tRNA expression cassettes as therapeutic cargo.

Human RNA pol III transcription is terminated at a series of four or more thymidines (in vertebrates), leaving a poly-U tail on the tRNA^26^. Facilitated recycling of human RNA pol III has been demonstrated with stable DNA-bound RNA pol III complexes going through many rounds of reinitiation^30^. Following transcription, the pre-tRNA is processed by endo- and exo-nucleases to remove the 3’-RNA trailer^34^. When screening the 3’ libraries with ACE-tRNA^Arg^_UGA_ both for the 4 bp immediately following the tRNA and the 35 bp downstream library, a range of PTC suppression responses was demonstrated (Fig. 1, Supplemental Fig. 2). Strikingly, when comparing the same ACE-tRNA^Arg^_UGA_ expression cassette sequence from the 55 bp 5’-UCE and 4 bp 3’-trailer library, a two-fold difference in nonsense suppression efficiency was noted (Fig. 1B and 1D). The only difference between these sequences was the identity of the nucleotides immediately following the polyT transcription terminator. However, the best-performing members of the 35 bp 3’-trailer library recovered the lost nonsense suppressor function (Fig. 1F), potentially due to enhanced pol III recycling from the native transcription terminators. While it was generally thought that tRNA expression cassettes are ‘plug and play’, these results highlight the importance of ensuring optimal ACE-tRNA expression cassette sequence context for use in therapeutic vectors.

Sup-tRNAs for genetic code expansion have been improved in several ways, including conversion of a U-G wobble pair to a C-G pair in the anticodon stem^59,70,71^. Conversion of A-U pairs to C-G pairs in the ACE-tRNA stems should increase the tRNA stability and subsequent steady state levels, providing a higher concentration of ACE-tRNAs for charging with their cognate amino acid. To approach this in a manner congruent with the ACE-tRNAs function in translation, we determined all A-U (or U-A) pairs for which another human isoacceptor of that tRNA contained a C-G (or G-C) pair at that position for ACE-tRNA^Arg^_UGA_ and ACE-tRNA^Leu^ (Fig. 2A, 2C). While no general trends emerged, data suggested that certain C-G pairs are favorable in conjunction with other pairs while others are not. When introduced in conunction, the two C-G pairs in the ACE-tRNA^Leu^_UGA_ variable loop are strongly unfavorable for PTC suppression efficiency but can improve function when introduced separately (Supplemental Fig. 4). These results emphasize that while the determination of which library members to include should be guided by available sequence information, there is still a need to screen these sequences in conjunction with each other to determine their additive impact on function.

tRNAs require unique structural elements for some steps of translation (e.g. charging by aaRSs and codon-anticodon decoding on the ribosome), while other steps in translation are better served by more uniform characteristics of the tRNA (e.g. EF1a transport of aminoacylated-(aa-) tRNAs to the ribosome and transition through the three states of the translation process). To satisfy both requirements, multiple components of the tRNA are tuned to present uniform thermodynamic properties to other parts of the translational apparatus. Tuning of tRNA interactions is perhaps best understood for the thermodynamic compensation of aa-tRNAs interaction with EF1a^87–89^. As EF1a interacts with both the amino acid and the t-stem of the aa-tRNA, both interactions contribute differentially to maintain uniform overall thermodynamics associated with each EF1a-aa-tRNA complex. Therefore, each aa-tRNA participates in translation somewhat differently where the esterified amino acid, the anticodon, and the tRNA body each make unique kinetic and/or thermodynamic contributions to maintain an overall uniformity in translation^46^. While ACE-tRNAs are generally esterified with their cognate amino acid and as such should already be tuned for interaction with EF1a, the codon-anticodon interaction has been altered. As uniformity of the aa-tRNA-EF1a-ribosome ternary complex relies partly on the codon-anticodon interaction, the aa-ACE-tRNA-EF1a-ribosome interaction can perhaps be tuned to improve the rate of translation. While improvements were modest when screening a small library of t-stems (Fig. 2E-H, Fig. 3C), our results indicate that this is an area of potential improvement for ACE-tRNAs.

Improvements for several sup-tRNAs have come from selection of sequence variants of the anticodon loop^73,75^. Even if an aaRS does not directly interact with the anticodon, subtle structural shifts in the tRNA can influence the aaRS-sup-tRNA interaction and could potentially be compensated for by altering the other nucleotides of the anticodon loop. To that end, we screened all possible combinations of the four non-anticodon-nucleotides of the anticodon loop for ACE-tRNA^Arg^_UGA_ (Fig. 2I). Only four members of the library showed appreciable levels of nonsense suppression, with the original anticodon loop sequence showing the highest PTC rescue. Perhaps this is not unsurprising as the four library members retaining function account for the only nucleotide in the library known to vary in all human tRNA^Arg^ sequences. Further, interactions between the anticodon loop and the Arg-aaRS (RARS) have been seen in structural characterizations of the tRNA^Arg^-RARS complex^90,91^. While there are trends in structure and function for type I and type II aaRSs, the tRNA-aaRS complex will be somewhat different for all isoacceptor pairs, and as such, while this library was not successful for ACE-tRNA^Arg^_UGA_, we cannot rule it out for other ACE-tRNAs. Indeed, modulation of the anticodon stem-loop has been shown to increase function for other sup-tRNAs^92^.

To promote the seamless rescue of PTCs during translation, the fidelity of translation should be maintained for the optimized ACE-tRNA sequences as demonstrated previously for ACE-tRNAs^16^. When assaying the protein products resulting from PTC rescue, each of the optimized ACE-tRNA^Arg^_UGA_, ACE-tRNA^Leu^_UGA_, and ACE-tRNA^Gly^_UGA_ sequences demonstrated >97% incorporation of the expected cognate amino acid, while the optimized ACE-tRNA^Trp^_UGA_ sequence showed incorporation of >99% arginine (Fig. 6). It has been noted that the anticodon serves as an anti-determinant between tRNA^Arg^ and TrpRS^40^. By converting the anticodon of tRNA^Trp^ to UCA along with alterations to the t-stem sequence, it appears the cognate specificity between tRNA^Trp^ and RARS was lost. These results highlight the necessity of determining translational fidelity of altered ACE-tRNA sequences.

The success of a genetic therapy is ultimately reliant upon adequate delivery of the genetic cargo to the tissue and cell type of interest. A sufficient quantity of the ACE-tRNA encoding vector must be delivered to clear the therapeutic threshold and produce a clinically meaningful response, and adequate delivery of genetic cargo has posed a major hurdle in the implementation of genetic therapies^93^. Despite major advances in viral^94^ and non-viral^95^ vector-based approaches, delivery of gene therapy-based genetic cargo remains a hurdle. However, because of the compact size of ACE-tRNAs, they are particularly amendable to existing and emerging delivery strategies^17^.

## ACKNOWLEDGMENTS

We thank members of the Lueck Laboratory and Dr. Amy M. Martin for reading and editing the manuscript and constructive discussion throughout the study. We would like to thank the Cystic Fibrosis Foundation Therapeutics Lab and Dr. Hillary Valley for providing 16HBE14o- and 16HBE14ge cell lines used in this study, and Matthew K Tanner for help with bioinformatics. This research has been facilitated by the services provided by the University of Rochester Mass Spectrometry Resource Laboratory and NIH instrument grant (S10OD025242) and the University of Rochester Genomics Research Center (GRC). We thank Kevin Welle at the University of Rochester Mass Spectrometry Resource Laboratory for helpful discussion. This work was supported by Cystic Fibrosis Foundation Postdoctoral Fellowship (PORTER20F0) to J.J.P., and a Cystic Fibrosis Foundation Research Grant (LUECK20GO), and NIH grant (R01 HL153988) to J.D.L.

## AUTHOR CONTRIBUTIONS

J.J.P., W.K., and J.D.L. designed the study. J.J.P., E.G.S., and W.K. performed experiments. J.J.P., E.G.S., and W.K. analyzed the data and constructed the figures. J.J.P., W.K., and J.D.L. wrote the manuscript. All authors read and revised the manuscript.

## DECLARATION OF INTERESTS

J.D.L. is a co-inventor of a technology presented in this study and receives royalty payments related to the licensing of the technology from the University of Iowa. PCT/US2018/059065, filed November 2, 2018; METHODS OF RESCUING STOP CODONS VIA GENETIC REASSIGNMENT WITH ACE-tRNA; Inventors - University of Iowa – Inventors J.D.L. and Christopher A. Ahern pertains to the tRNA sequences used in this study. J.D.L. and J.J.P. are co-inventors of a technology presented in this study and receives royalty payments related to the licensing of the technology from the University of Rochester. PCT/US2023/62053, filed February 6th, 2023; OPTIMIZED SEQUENCES FOR ENHANCED tRNA EXPRESSION OR/AND NONSENSE MUTATION SUPPRESSION; Inventors - University of Rochester – Inventors J.D.L. and J.J.P pertains to the tRNA sequences used in this study.

## METHODS

### Nonsense reporter high throughput cloning and screening (HTCS) plasmid

The HTCS plasmid (pNanoRePorter 2.0) used in this study was based on a previously used HTCS plasmid^16^ with the addition of a UbC-Fluc2**-**SV40 pA expression cassette for transfection normalization. The UbC-Fluc2-SV40 pA cassette was synthesized as DNA gBlocks (Integrated DNA Technologies, IDT) and inserted by Gibson assembly using NEBuilder HiFi assembly mix (New England Biolabs). Unless otherwise noted, all molecular biology reagents were obtained from New England Biolabs. A version of the pNanoRePorter 2.0 plasmid was cloned with the negative selection cassette placed upstream (5’), spanning, or downstream (3’) of the ACE-tRNA for cloning each of the ACE-tRNA expression cassette libraries via golden gate assembly (see supplemental information for sequences). All intragenic and extragenic ACE-tRNA library members (except the 850 bp 5’ UCE) were obtained as DNA oligonucleotides (IDT) and were annealed and cloned as previously described with some modifications^16^. The 850 5’ UCE clones were order as double stranded DNA eBlocks (IDT) and cloned via Golden Gate assembly. For this study, the *E. coli* cultures were grown in 96-well plates (Enzyscreen) and the NucleoSpin 96 Plasmid Transfection, 96-well kit was used to miniprep the plasmid DNA (Macherey-Nagel). All plasmid DNA was normalized to 50 ng/μL and the sequence verified by Sanger sequencing (Eurofins Scientific). The transcription tabulator sequences and fully optimized ACE-tRNA expression cassettes were obtained as gBlocks (IDT) and assembled into pNanoRePorter 2.0 via golden gate assembly. The pUC57 mini plasmid used as carrier cDNA for transfections was gigaprepped as per kit instructions (Macherey-Nagel). The normalized ACE-tRNA library cDNA at 50 ng/μL was mixed at equal volumes with pUC57 mini carrier plasmid at 150 ng/μL to give final concentrations of 25 ng/μL and 75 ng/μL of the library and carrier plasmids respectively.

### High throughput screening of the ACE-tRNA libraries

HEK293T cells (ATCC) were cultured in Dulbecco’s Modified Eagle Medium (DMEM) supplemented with 10% FBS, 1% Penicillin-Streptomycin, and 2 mM L-glutamine (Thermo Fisher). 16HBE14o-cells were cultured in minimal essential medium (MEM) supplemented with 10% FBS, and 1% Penicillin-Streptomycin as previously described^22^. Cells were plated at 3 x 10^4^ cells per well immediately before transfection. The HTCS/carrier plasmid mix was transfected in duplicate 96-well plates using lipofectamine 2000 (Thermo Fisher). Lipofectamine 2000 reagent was prepared for transfection by mixing at 0.3 μL Lipofectamine 2000 per 7.2 μL of OptiMEM (Thermo Fisher). The cDNA (3 μL at 100 ng/μL per well) was mixed with 12 μL OptiMEM, followed by 15 μL of the Lipofectamine 2000/OptiMEM mix. After 5 minutes of incubation at RT, 10 μL of the cDNA/Lipofectamine mix was delivered to each well of cells. Twenty-four hours post-transfection, the media was aspirated and 15 μL of PBS was added to each well. Fifteen microliters of reconstituted One-Glo Ex Reagent (Promega) was added to each well, the plate was incubated shaking at 600 rpm for 3 minutes, and the Fluc luminescence was measured by a Synergy2 multi-mode microplate reader (Biotek Instruments). After measuring the Fluc luminescence, 15 μL of Nano DLR Stop & Glo reagent was added to each well, and the plate was incubated shaking at 600 rpm for 10 minutes. After quenching the Fluc luminescence, 15 μL of NanoDLR Stop & Glo substrate (furimazine) diluted 1:100 in PBS was added to each well, the plate was incubated shaking at 600 rpm for 3 minutes, and the Nluc luminescence was measured with the microplate reader. The normalized suppression ratio is calculated from the equation (PTC-NanoLuciferase luminescence [+ACE-tRNA]/Firefly luminescence)/(PTC-Nanoluciferase luminescence [no ACE-tRNA]/Firefly luminescence).

### Determination of position weighted matrices (PWM)

The position weighted matrices for each of the 55 bp 5’ UCE and 35 bp 3’ trailer sequences were determined using kpLogo with unweighted user input^77^. The nonsense suppression efficiency of the ACE-tRNA sequence library members were directly converted to sequence abundance (i.e. a normalized suppression ratio of 100 was treated as equivalent to 100 copies of that ACE-tRNA sequence) as kpLogo is computed based on sequence abundance. For the frequency logos, the residues are scaled relative to their frequencies at each position, while in the probability logo, residues are scaled relative to the statistical significance (-log10(*P* value)) of each residue at each position.

### Measurement of tRNA transcript tabulator expression via RT-qPCR

The self-cleaving ribozyme drz-Bflo-2^78^ was appended immediately downstream of the ACE-tRNA sequence for measurement of steady-state ACE-tRNA levels using RT-qPCR. Plasmid DNA containing the ACE-tRNA TTT sequences was midiprepped with the NucleoBond PC 20 miniprep kit (Macherey-Nagel). 16HBE14o-cells were diluted to 6 x 10^5^ cells per mL with 1 mL added per well of a coated 6-well plate as previously described^22^. For each well, 75 ng of TTT containing DNA was mixed with 2425 ng of pUC57 carrier DNA, 150 μL of OptiMEM containing 3 μL of PLUS reagent (Thermo Fisher) was mixed with the cDNA, followed by 150 μL of OptiMEM containing 9 μL of LTX reagent (Thermo Fisher). The cDNA/Lipofectamine mixture was incubated for 7 minutes at room temperature, before 250 μL of the mixture was delivered dropwise to each well of cells. All transfections were performed in triplicate. Twenty-four hours post-transfection the media was exchanged for fresh media. Forty eight hours post-transfection the media was aspirated and the cells washed with DPBS (Thermo Fisher), trypsinized with 0.5 mL TrypLE (Thermo Fisher), dislodged from the culture plate in 1.3 mL MEM complete media, and transferred to a microcentrifuge tube. The cell suspension was centrifuged at 700 rcf at 4 °C for 5 minutes, the media aspirated, the cell pellet loosened by flicking the tube, and 350 μL of RLT Buffer was used to lyse the cells. Lysed cell material was stored overnight at −80 °C before total RNA was isolated using the RNeasy Plus Mini Kit (Qiagen). RNA isolation and RT-qPCR was performed by the University of Rochester Genomics Research Center. High RNA quality was demonstrated for all samples using an Agilent Bioanalyzer. FAM labeled TTT probe and TET labeled TATA Binding Protein (TBP) probe were obtained from IDT and used with TaqMan Fast Master Mix (Thermo Fisher) on an Applied Biosystems QuantStudio 12k Flex Real-Time PCR System. Each sample was quantified in triplicate, and a no-template control containing water instead of RNA, was included as a negative control. The quantification of the TTT for each ACE-tRNA are represent by the calculated QuantStudio relative target quantity (RQ) value of gene expression for the technical replicate group associated with each RNA sample.

### sfGFP protein expression, purification, and mass spectrometry sample preparation

HEK293T cells (ATCC, USA) were grown in standard conditions as outlined above. A pcDNA 3.1/Hygro(+) plasmid expressing sfGFP-TGA-Strep-6xHis-Strep was co-transfected with a plasmid expressing 4 copies of the original or optimized (Opt) ACE-tRNA expression cassettes into HEK293T cells at 75% confluency using Calfectin (SignaGen Laboratories) according to standard protocols. At 36 hours post-transfection the cells were trypsinized, pelleted at 700 rcf for 10 minutes at 4 °C, and washed once in DPBS. The cell pellets were resuspended in Strep-Tactin XT wash buffer (100 mM Tris-HCl, pH 8.0, 150 mM NaCl, 1 mM EDTA, and protease inhibitors (Medchem Express). The cells were lysed by thorough Dounce homogenization, cell debris was removed by centrifugation at 20 k rcf at 4 °C for 30 minutes, and the soluble lysate was filtered through a 0.22 μm filter. Strep-tactin XT resin (IBA Lifesciences) was washed with 50 bed volumes of wash buffer, the soluble cell lysate was mixed with the resin, the resin was washed with 500 bed volumes of wash buffer, and the sfGFP was eluted in 5 bed volumes of elution buffer (wash buffer without protease inhibitors, containing 50 mM D-biotin). The eluted protein was resolved on 10-20% Novex WedgeWell SDS-PAGE gels (Invitrogen) and stained with SimplyBlue SafeStain Coomassie (Thermo Fisher) or mass spectrometry compatible silver stain (Pierce). The bands of interest were excised from the gel, cut into 1 mm cubes, de-stained, then reduced and alkylated with DTT and IAA, respectively (Sigma), and dehydrated with acetonitrile. Mass Spectrometry grade trypsin (Promega), reconstituted at 10 ng/μL 50 mM ammonium bicarbonate. Gel pieces were incubated for 30 minutes at room temperature in just enough trypsin solution to entirely cover the pieces. After the intial incubation, additional ammonium bicarbonate was added until the gel pieces were completely submerged and incubated at 37 °C overnight. Tryptic digest peptides were extracted by the addition of 50% acetonitrile, 0.1% TFA, dried in a CentriVap concentrator (Labconco), desalted with homemade C18 spin columns, dried again, and reconstituted in 0.1% TFA.

### Mass spectrometry conditions

Peptides were injected onto a homemade 30 cm C18 column with 1.8 μm beads (Sepax) using an Easy nLC-1200 HPLC (Thermo Fisher). Solvent A was 0.1% formic acid in water, while solvent B was 0.1% formic acid in 80% acetonitrile. The elution gradient began at 3% solvent B held for 2 minutes, increased to 10% B over 5 minutes, increased to 38% B in 38 minutes, then increased to 90% B in 3 minutes and was held for 3 minutes before returning to the starting buffer conditions in 2 minutes, before finally re-equilibrating for 7 minutes, for a total run time of 60 minutes. A Nanospray Flex ion source operating at 2 kV was used to introduce the ions to a Fusion Lumos Tribrid mass spectrometer (Thermo Fisher). The Fusion Lumos was operated in targeted data-dependent mode, with both MS1 and MS2 scans acquired in the Orbitrap. The cycle time was set to 1.3 seconds, the monoisotopic precursor selection (MIPS) was set to peptide, the full scan was done over a range of 375-1400 m/z, with a resolution of 120,000 at m/z of 200, and AGC target of 4×10^5^, and a maximum injection time of 50 ms. Two branches were created to determine which precursor ions to fragment. An inclusion list comprised of the 20 theoretical tryptic peptides from the single amino acid substitution of sfGFP at p.150 was created, and ions that matched the m/z of these peptides were preferentially selected for fragmentation if the mass was within 7 ppm, regardless of intensity. Otherwise, peptides with a charge state between 2-5 were picked for fragmentation based on their intensity. Precursor ions were fragmented by high energy collisional dissociation (HCD) using a collision energy of 30% with an isolation width of 1.5 m/z. When fragmenting ions from the targeted inclusion list, the maximum injection time was set to 100 ms; otherwise it was set to 22 ms. For all MS2 scans, the AGC target was set to 5×10^4^, and the resolution was set to 15,000. The dynamic exclusion was set to 20 seconds, but was only used for peptides not on the targeted inclusion list.

### Mass spectrometry data analysis

A FASTA database was created that contained each of the 20 theoretical protein sequences that could arise from the single amino acid substitution at sfGFP p.150. The raw data was searched using the SEQUEST search engine within the Proteome Discoverer software platform, version 2.4 (Thermo Fisher), using the sfGFP p.150 FASTA database appended to the SwissProt *Homo sapiens* database. Trypsin was selected as the enzyme allowing up to 2 missed cleavages, with an MS1 tolerance of 10 ppm, and an MS2 tolerance of 0.025 Da. Carbamidomethyl was set as a fixed modification, while oxidation of methionine (15.995 Da), and N-terminal methionine-loss were set as variable modifications. Percolator was used as the FDR calculator, filtering out peptides which had a q-value greater than 0.01. The Minora Feature Detector Node was used to integrate peptides and determine relative abundance values.

### PTC-CFTR-Nluc expression and in-gel luminescence

CFTR cDNA with a C-terminal Nluc fusion was cloned via Gibson assembly downstream of a short ubiquitin C (UbC) promoter in pUC57 mini. PTCs were introduced into the CFTR cDNA construct using PCR based site-directed mutagenesis. The CFTR expression plasmids were co-transfected along with a plasmid expressing 4 copies of the original or optimized (Opt) ACE-tRNA expression cassettes into HEK293T cells using Lipofectamine LTX (Thermo Fisher) as outlined above. For all transfections 750 ng of CFTR-Nluc plasmid was used, the amount of 4x ACE-tRNA plasmid was varied with the remainder up to 700 ng being composed of pUC57 carrier DNA, and 50 ng WT-sfGFP plasmid was included in the mix as a transfection control. Transfected HEK293T cells were incubated for 24 hours before the media was aspirated, the cells washed with DPBS, and lysed in RIPA buffer (Invitrogen) containing protease inhibitors (Medchem Express). Insoluble cell debris was removed by centrifugation at 20k rcf for 30 minutes at 4 °C, total protein was quantified by BCA assay (Pierce), and 10 μg of protein was resolved on 4-12% Novex WedgeWell SDS-PAGE gels (Invitrogen). Following electrophoresis, the SDS-PAGE gel was rinsed once in water and the GFP fluorescence was imaged in-gel using a ChemiDoc Imaging System (Biorad). The SDS-PAGE gel was then washed twice for 15 minutes in 25% isopropanol in water, twice for 15 minutes in water, incubated for 5 minutes in NanoGlo buffer containing 1:500 NanoGlo reagent (furimazine), and the in-gel luminescence imaged on the ChemiDoc Imaging System. Quantification of GFP and CFTR-Nluc band intensity was determined using Image Lab version 6.1.0 (Biorad).

### Rescue of PTC-containing CFTR mRNA from 16HBEge cell line

Rescue of PTC-containing CFTR mRNA following ACE-tRNA treatment was conducted as previously described^22^. Briefly, total RNA was isolated using a Monarch Total RNA Miniprep Kit (NEB), RNA quantity and quality were determined with a NanoDrop One^C^ Spectrophotometer (Thermo Fisher), one-step RT-qPCR was performed using a QuantStudio 3 Real-Time PCR System (Applied Biosystems) using the Luna Universal One-Step RT-qPCR Kit (NEB), and then analyzed with QuantStudio Design & Analysis Software v.1.5.1. Each sample was quantified in triplicate, and a no-template control reaction, with just nuclease-free water, was included as a negative control. The fold difference in CFTR gene expression normalized to TATA binding protein (TBP) was calculated using the comparative Ct method, 2^-ΔΔCt^.

## Notes

### Competing Interest Statement

JDL is on the SAB for hC Biosciences, Inc.

