## Supplemental Data and Information for "Optimization of Anticodon Edited Transfer RNAs (ACE-tRNAs) Function in Translation for Suppression of Nonsense Mutations"

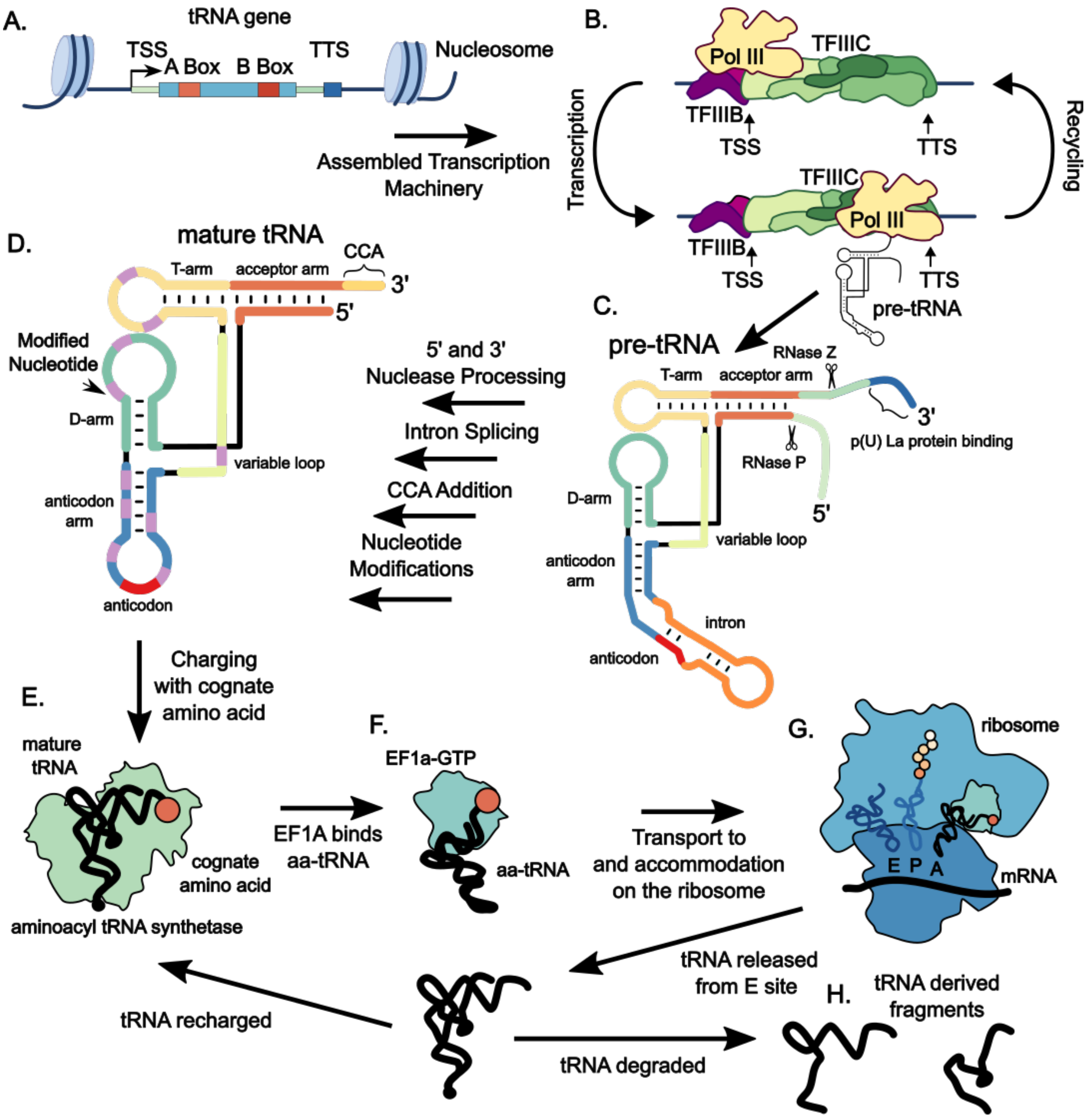


**Supplemental Figure 1. Natural tRNA biogenesis and function in translation** (**A**) The human genome contains ~500 tDNAs[1], which are subjected to epigenetic regulation of expression. While nearly half of human tDNAs are silent[2], actively expressed tDNAs display nucleosome-free gene cassettes, delimited by nucleosomes centered ~150 bp upstream of the transcription start site (TSS) and ~125 bp downstream of the transcription termination signal (TTS)[3]. (**B**) tDNA located in areas of active chromatin are transcribed by RNA polymerase III (Pol III). tRNAs are expressed from type 2 Pol III promoters with the multi-subunit transcription factor III C (TFIIIC) binding to A and B box sequences (orange and red boxes in panel A) which are internal to the mature tRNA sequence (light blue box in panel A)[4-6]. TFIIIC then recruits the multi-subunit transcription factor IIIB (TFIIIB) to the tRNA upstream control element sequence ~50 bp upstream of the tRNA which contains a weak TATA box and the TSS[7]. TFIIIB then recruits RNA Pol III which transcribes the pre-tRNA. Following formation of the transcription complex multiple rounds of transcription take place before the factors dissociate from the tRNA gene[8]. The TTS consists of a stretch of greater than four (in humans) thymidines[4]. (**C**) The pre-tRNA transcript is synthesized with 5'-leader and 3'-trailer sequences at either end. The short polyU tract remaining at the 3' end of the pre-tRNA serves as the binding site for the La protein, which protects the 3' end of the transcript from spurious exonuclease digestion and helps with pre-tRNA folding[9]. (**D**) The 5'-leader and 3'-trailer are removed through the endonuclease action of the ribonucleoprotein RNase P (5' end) and RNase Z (3' end), along with the action of other exonucleases[10-12]. Tuning has been demonstrated in pre-tRNA binding to RNase P between the 5' leader sequence in the pre-tRNA and the tRNA body[13]. Following 3' cleavage, the CCA adding enzyme catalyzes the addition of CCA nucleotides to the tRNA 3' terminus without the need for a template[14]. A subset of tRNAs contain introns which are primarily located immediately 3' to the anticodon. As these tRNA introns disrupt the anticodon arm, they must be removed via splicing to produce a functional mature tRNA[15]. Following processing, ~12% of tRNA nucleotides are modified with one of many modifications, both before and after trafficking from the nucleus to the cytoplasm[16,17]. tRNA chemical modifications play a role in all aspects of tRNA function including, structure, stability, aminoacylation, and decoding at the ribosome[17]. (**E**) Mature tRNAs interact with their cognate aminoacyl-tRNA synthetase (aaRS) and are charged with their cognate amino acid[18,19]. Recognition by the correct aaRS represents the first major step in maintaining translational fidelity, along with ensuring the correct codon-anticodon pairing on the ribosome. (**F**) The aminoacyl tRNA (aa-tRNA) is then passed to GTP-bound translation elongation factor 1a (EF1a), which shuttles the aa-tRNA to the ribosomal A site, forming a ribosome-EF1a-aa-tRNA ternary complex[20,21]. Sequences in the t-stem of tRNAs are primarily responsible for tuning affinities for EF1a[22,23]. (**G**) If the codon-anticodon interaction is correct, the EF1a-bound GTP is hydrolyzed, and EF1a releases the aa-tRNA, which transits the ribosome extending the polypeptide chain in protein synthesis[24]. Binding of the aa-tRNA with EF1a to the A site of the ribosome and interactions with the mRNA occurs in discrete steps that require flexibility in the tRNA structure[25-27]. (**H**) While tRNAs are generally stable, exhibiting a half-life of 2-3 days in eukaryotes[28], their steady state level is influenced by complex turnover mechanisms[29]. tRNA derived fragments, formed following tRNA breakdown have been increasingly shown to influence biology. Given the importance of RNA fragments in a number of aspect of gene regulation, the emerging role of tRNA fragments in development and disease is not surprising[30,31].

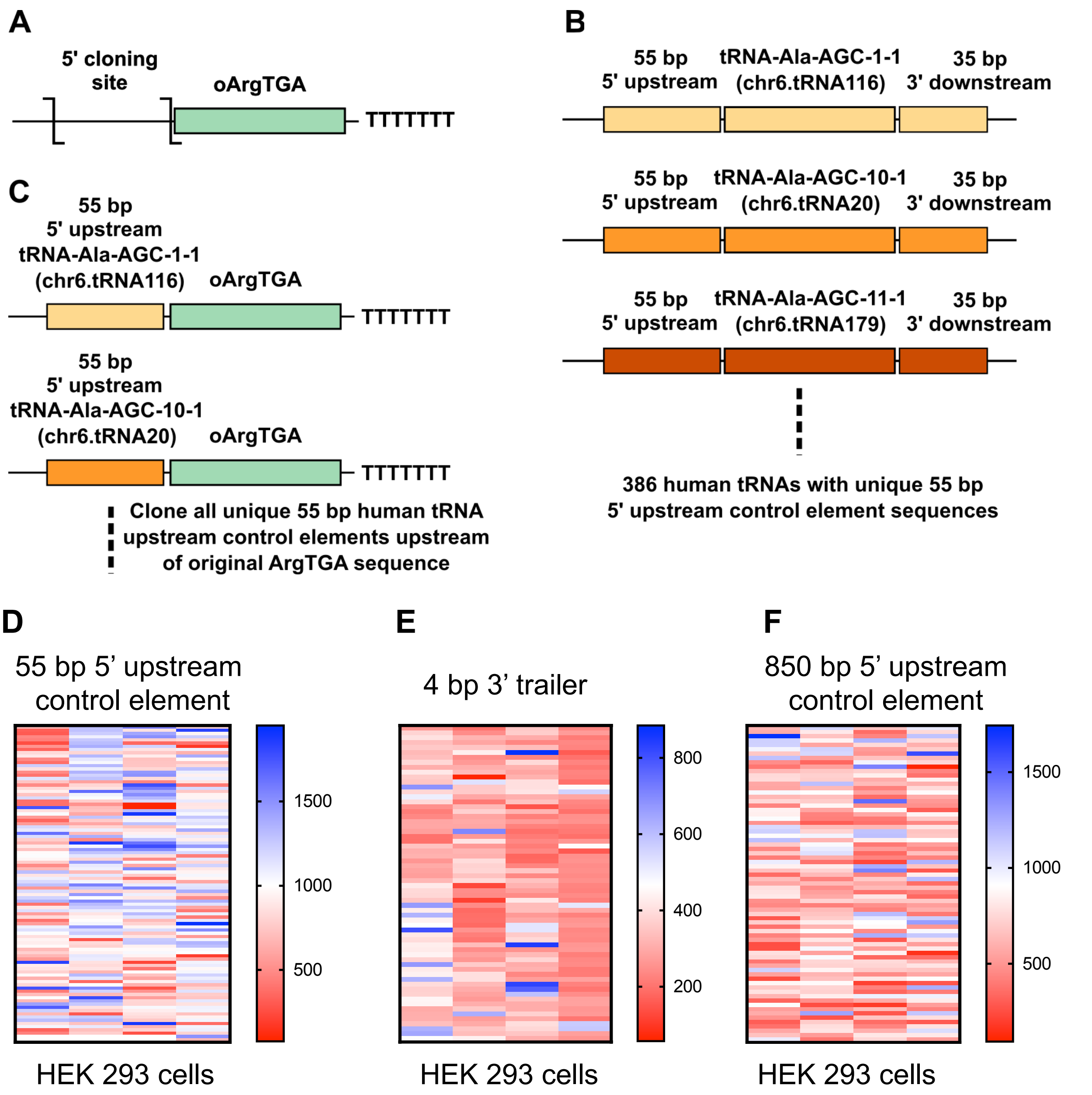


**Supplemental Figure 2. Derivation of 5' upstream control element sequences and results of extragenic screens for HEK293 cells** (A) The parent high-throughput-cloning and -screening vector for the tRNA 5' region, contains a cloning site immediately 5' to the original ArgTGA (oArgTGA) tRNA sequence/the original 3' trailer sequence. (B) All human tRNAs are denoted with a name like "tRNA-Ala-AGC-1-1" (where Ala is the three-letter amino acid code for the tRNA isotype, AGC is the anticodon, the first 1 corresponds to the numeric ID of a unique tRNA transcript or "isodecoder", and the second 1 corresponds to the gene locus ID – for tRNAs that have multiple identical copies, this gene locus ID represents the particular gene copy in the genome). Scanning the entire human genome returns 386 predicted tRNAs with a unique 55 bp sequence immediately upstream of the tRNA. (C**)** To determine the impact of these different sequences on the function of ACE-tRNAs we had each unique 55 bp 5' upstream control element (UCE, what we call this 5' upstream transcriptional element) synthesized as complementary oligonucleotides, which when annealed provide overhangs for golden gate cloning into our parent vector shown in (A). (D) Heat map representing the results of screening a 386-member library of 55-bp 5’ UCE sequences derived from each unique tRNA gene 5’ UCE in the human genome in HEK293 cells. (E) Heat map representing the results of screening a 256-member library of 4-bp 3’ trailer sequences representing every 4-bp combination of nucleotides following the ACE-tRNA in HEK293 cells. (F) Heat map representing the results of screening a 326-member library of 850-bp 5’ UCE sequences derived from every synthetically accessible, unique, 5’ UCE in the human genome in HEK293 cells. All values displayed in these heat maps represent the average of 6 independent transfections of HTCS library members. The normalized suppression ratio shown here is calculated from the equation (PTC-NanoLuciferase luminescence [+ACE-tRNA]/Firefly luminescence)/(PTC-Nanoluciferase luminescence [no ACE-tRNA]/Firefly luminescence).

A

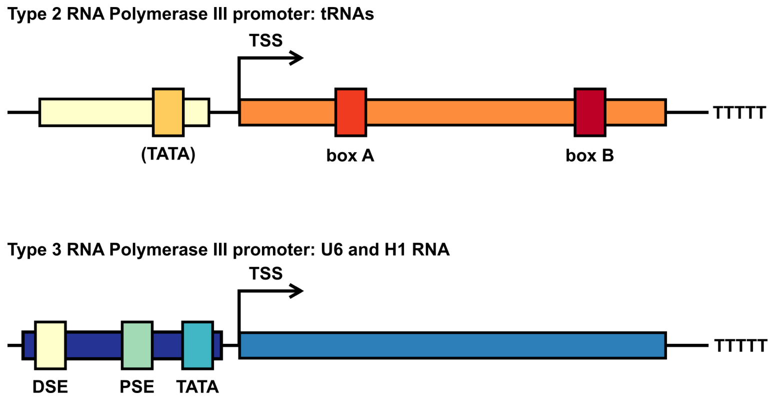


B

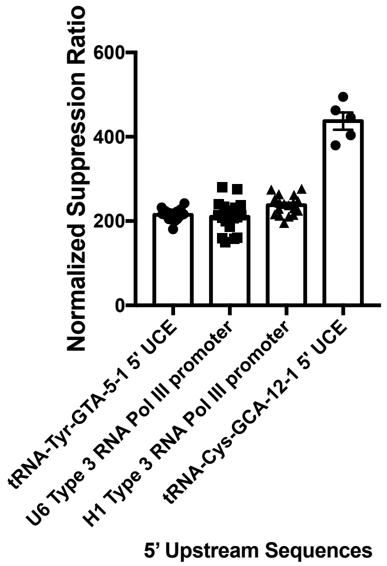


**Supplemental Figure 3. Types of RNA Pol III promoters and their impact on ACE-tRNA function** (A) Type 2 RNA Pol III promoters are employed to expressed tRNA genes in humans. Intragenic box A and box B sequences recruit TFIIIC, which then recruits TFIIIB, which then recruits RNA Pol III. Type 3 RNA Pol III promoters express other genes including U6 and H1 RNAs. Type 3 promoters do not require any intragenic sequences and as such are often used to express exogenous RNAs including CRISPR guide RNAs, siRNAs, and shRNAs which do not contain native A or B boxes. Type 3 promoters have been used to express human nonsense suppressor tRNAs although many of the endogenous human 55 bp 5'-leaders yield higher nonsense suppression activity *in vivo*. (B) Each of the 5' upstream sequences was cloned upstream of ACE-tRNA$^{Arg}_{UGA}$ and transfected into 16HBE14o- cells. tRNA-Tyr-GTA-5-1 5' UCE represents the original 5' UCE and tRNA-Cys-GCE-12-1 5' UCE represents the best-performing 5' UCE obtained from the 55-bp 5' UCE screen. The normalized suppression ratio shown here is calculated from the equation (PTC-NanoLuciferase luminescence [+ACE-tRNA]/Firefly luminescence)/(PTC-Nanoluciferase luminescence [no ACE-tRNA]/Firefly luminescence). The error bars represent the standard error of the mean.

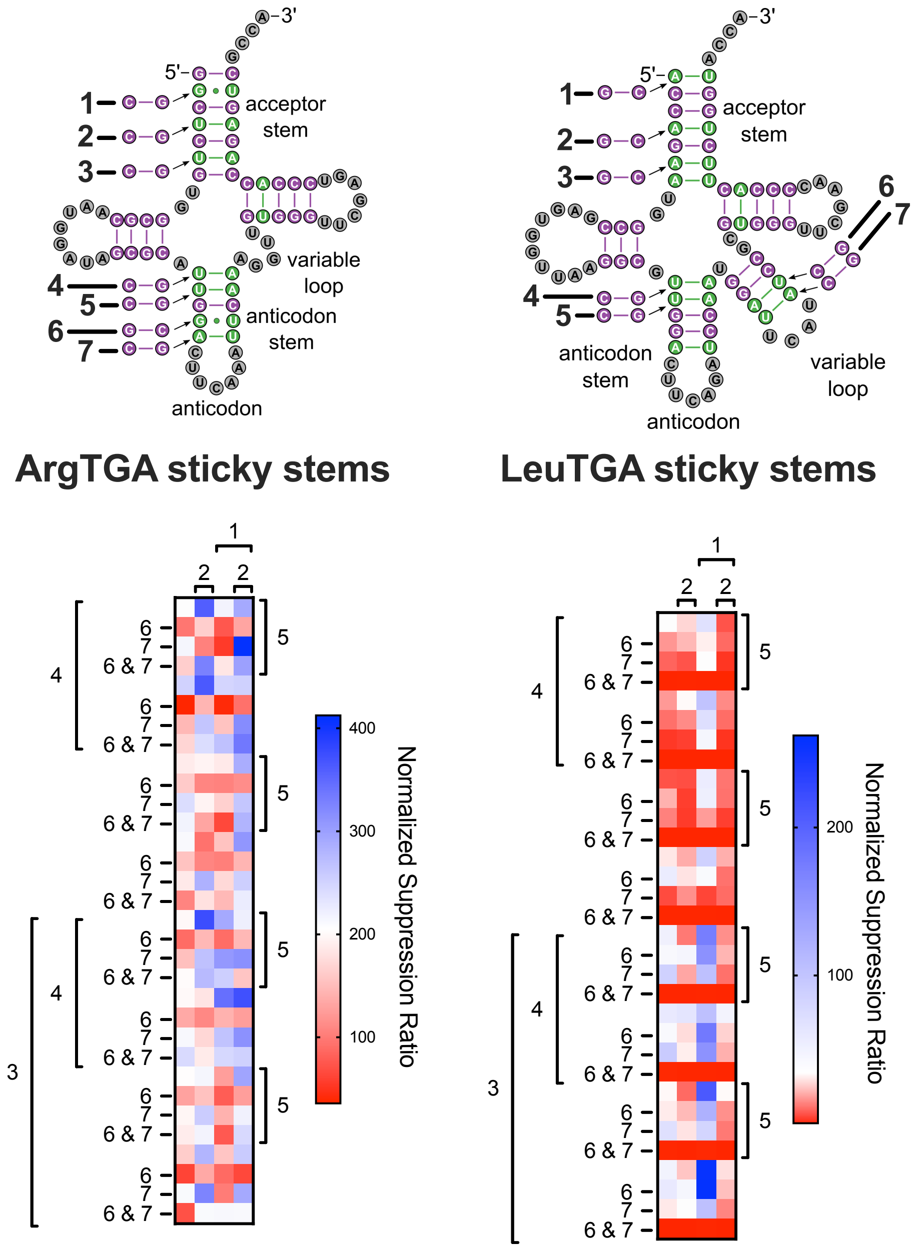


**Supplemental Figure 4. Influence of individual ACE-tRNA$^{Arg}_{UGA}$ and ACE-tRNA$^{Leu}_{UGA}$ sticky stem sites on nonsense suppressor efficiency.** Sites for each of the sticky stem library pairs are labeled 1-7 on the tRNA diagrams. The heatmap representations of each sticky stem screen are shown below. The presence of a number means that this library member is present in the sequence, while no number means the original pair is present in that sequence.

**A**

| Variant (Saks et al.) | Variant (as described here) | T-stem positions 49-65 | 50-64 | 51-63 |
|---|---|---|---|---|
| 1 | TS-1 | A-U | C-G | A-U |
| 2 | TS-2 | A-U | C-G | G-C |
| 4 | TS-3 | A-U | C-G | U-A |
| 5 | TS-4 | G-C | C-G | A-U |
| 6 | TS-5 | G-C | C-G | G-C |
| 7 | TS-6 | G-C | C-G | C-G |
| 8 | TS-7 | G-C | C-G | U-A |
| 9 | TS-8 | C-G | C-G | A-U |
| 10 | TS-9 | C-G | C-G | G-C |
| 11 | TS-10 | C-G | C-G | C-G |
| 12 | TS-11 | C-G | C-G | U-A |
| 13 | TS-12 | U-A | C-G | A-U |
| 14 | TS-13 | U-A | C-G | G-C |
| 15 | TS-14 | U-A | C-G | C-G |
| 16 | TS-15 | U-A | C-G | U-A |
| 17 | TS-16 | G•U | C-G | A-U |
| 18 | TS-17 | G•U | C-G | G-C |
| 19 | TS-18 | G•U | C-G | C-G |
| 20 | TS-19 | G•U | C-G | U-A |
| 21 | TS-20 | A-U | C-G | G•U |
| 22 | TS-21 | G-C | C-G | G•U |
| 23 | TS-22 | C-G | C-G | G•U |
| 25 | TS-23 | G•U | C-G | G•U |
| 27 | TS-24 | C-G | G-C | U-A |
| 31 | TS-25 | A-U | G•U | C-G |
| 33 | TS-26 | C-G | G-C | C-G |
| 35 | TS-27 | G•U | A-U | A-U |

**B**

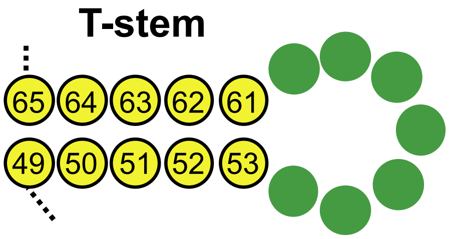


**Supplemental Figure 5. T-stem mutations tested** (A) Sequences for each of the t-stem nucleotide pairs 49-65, 50-64, 51-63. These sequences were derived from (Saks et al., 2011), with the variant numbering in the first column representing the number scheme used in Saks et al. and the numbering in the second column representing the numbering scheme used in this paper. (B) Depicts a generic t-stem with canonical tRNA nucleotide position numbering.

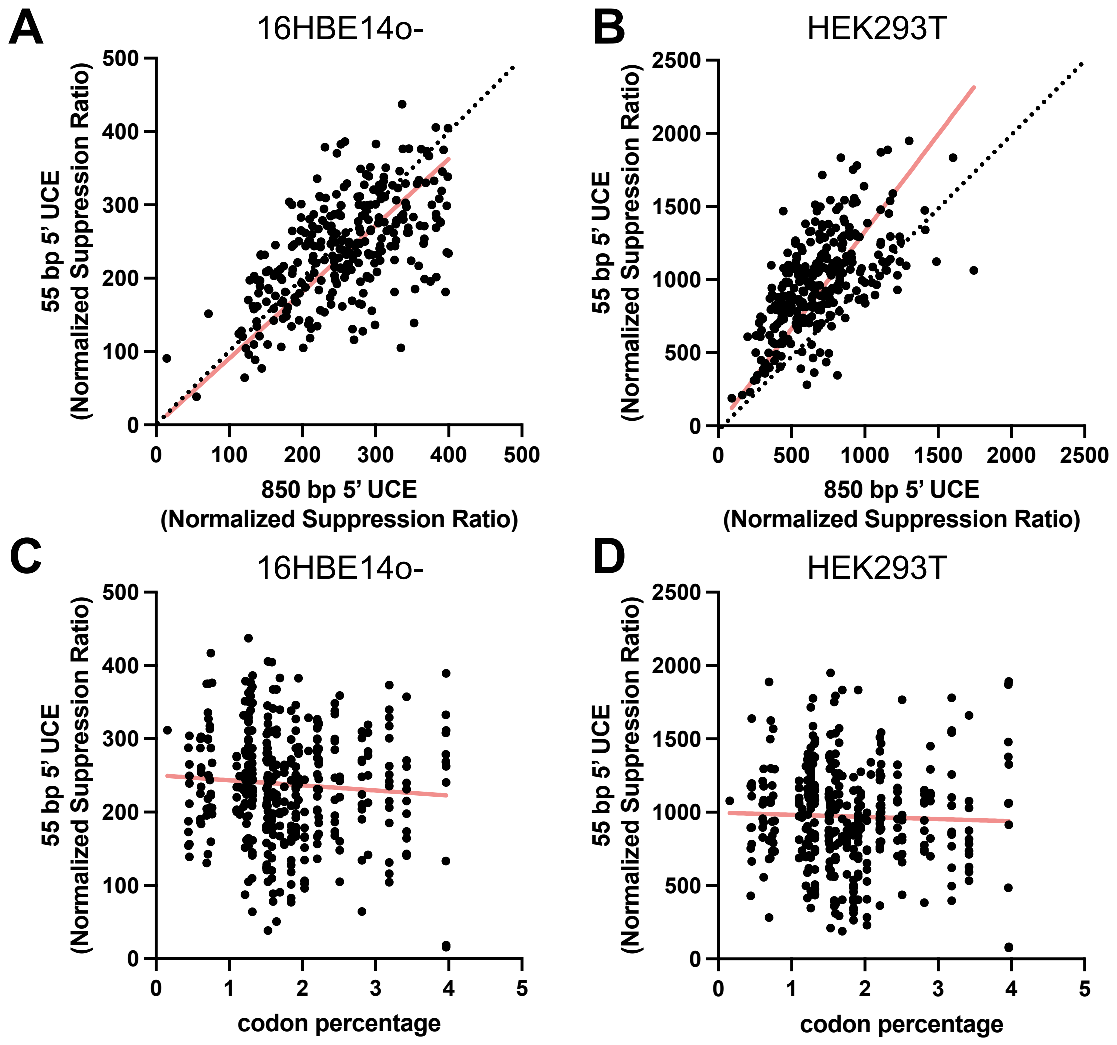


**Supplemental Figure 6. Comparison of 5' UCE sequences.** The influence of the 55 bp versus 850 bp 5' UCE sequences on the nonsense suppression efficiency of ACE-tRNA$^{Arg}_{UGA}$ was compared in both 16HBE14o- (A) and HEK293T (B) cell lines. The line of best fit (y-intercept set to y = 0) for each data set (red line) was determined using Prism 10 (slope = 0.9065 for A and slope = 1.328 for B), with the dotted black line representing a line with a slope of 1. The influence of the 55 bp 5' UCE sequences were also compared with codon abundance in the human transcriptome for both 16HBE14o- (C) and HEK293T (D) cell lines. For each 55 bp 5' UCE, the cognate codon corresponding to the tRNA from which it was derived (Supplemental Fig. 2) was determined, and the percent of all codons in the human transcriptome was determined for each codon. The line of best fit for each data set (red line) was determined using Prism 10 ($R^2$ = 0.005455 for C and $R^2$ = 0.001112 for D).

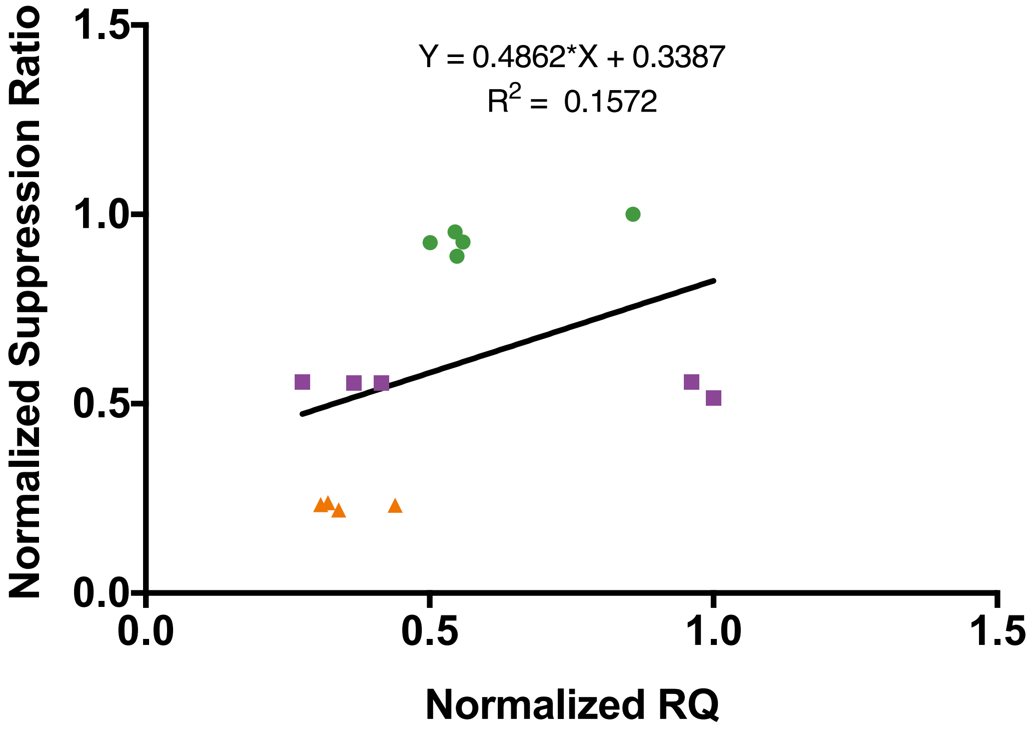


**Supplemental Figure 7. Effect of 5' UCE sequences on ACE-tRNA$^{Arg}_{UGA}$ nonsense suppression activity compared to impact of the same 5' UCE sequences on steady-state ACE-tRNA$^{Arg}_{UGA}$ expression levels** The equation for the line of best fit is displayed on the graph with the $R^2$ value

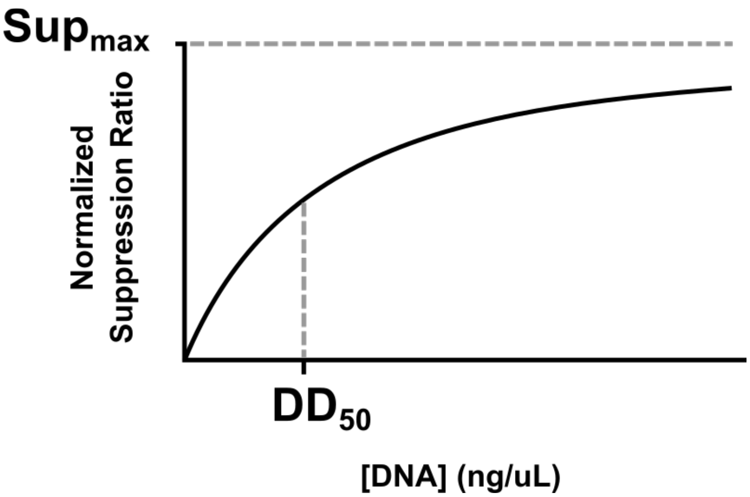


**Supplemental Figure 8. Model to fit DNA dependence of ACE-tRNA nonsense suppressor efficiency** This diagram outlines the parameters of the hyperbolic model to fit the DNA concentration dependence of ACE-tRNA nonsense suppression efficiency. The model is defined by the equation Normalized Suppression Ratio = $(Sup_{max} * [DNA])/(DD_{50} + [DNA])$ where $Sup_{max}$ corresponds to the maximal level of nonsense suppression displayed by the ACE-tRNA and $DD_{50}$ (Delivered DNA) corresponds to the concentration of DNA (ng/μL) required for a half-maximal nonsense suppression response.

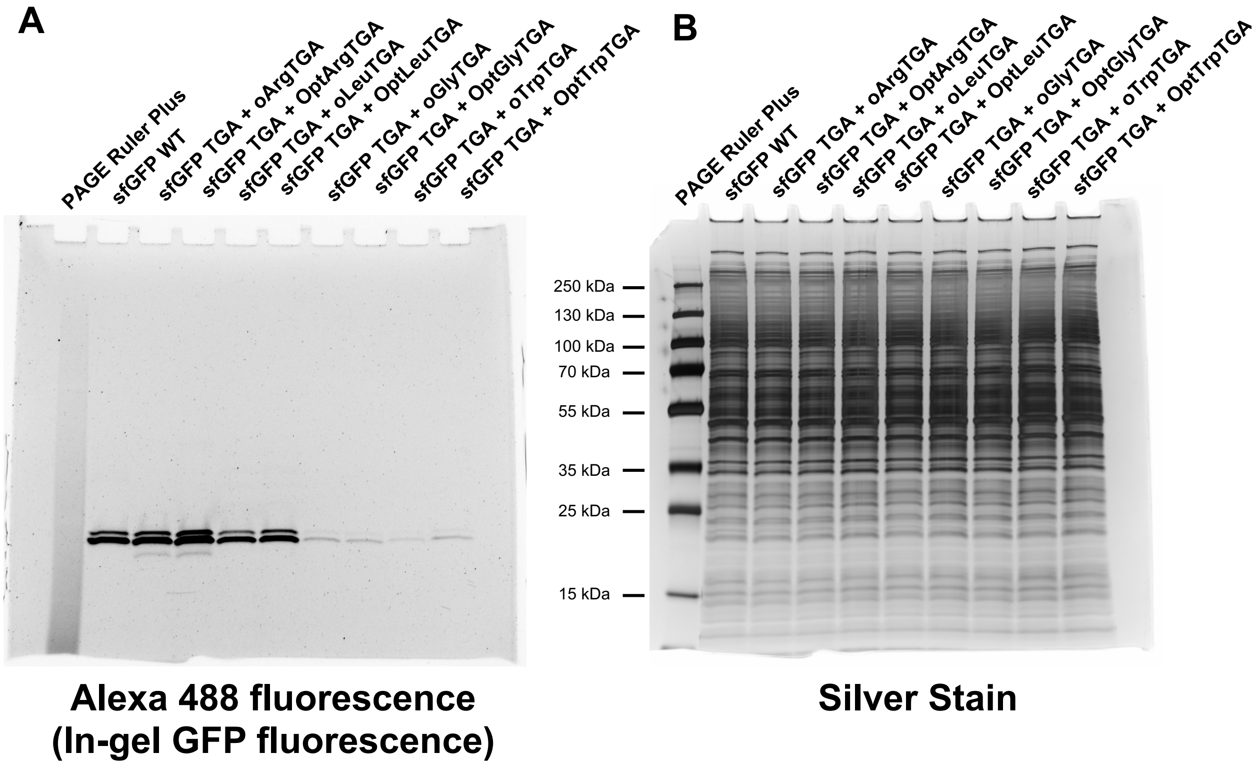


**Supplemental Figure 9. Uncropped gel images of gels shown in Figure 6B** (A) Alexa 488 fluorescence image of the crude 10-20% gradient SDS-PAGE gel (Invitrogen) of HEK293 cell lysate containing WT- or sfGFP-TGA-150 (B) subsequent silver staining of the crude SDS-PAGE gel.

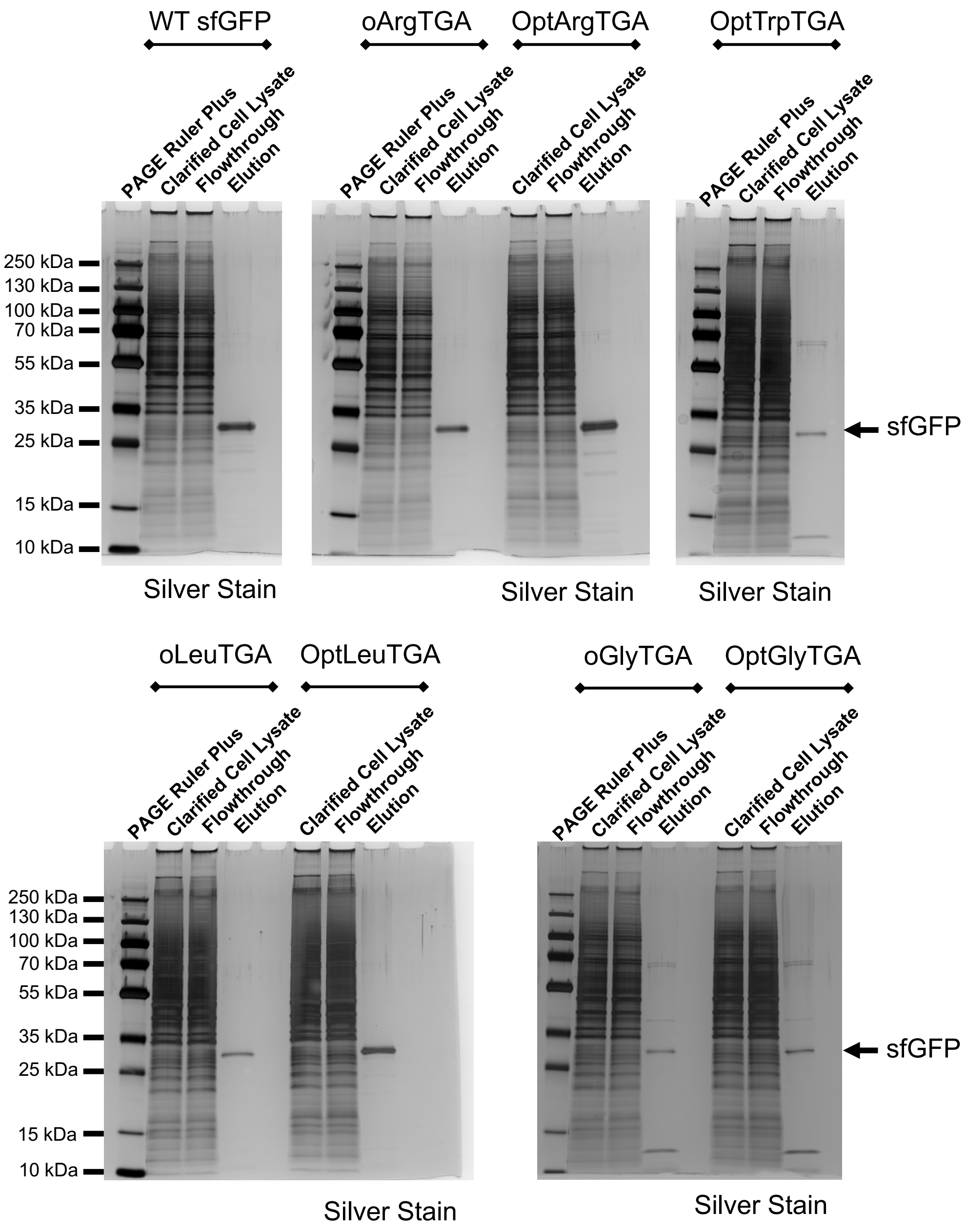


**Supplemental Figure 10. Purification of sfGFP from HEK293 cells** A construct expressing sfGFP-UGA-150-Strep-8xHis-Strep with a construct expressing 4 copies of the ACE-tRNA as noted above were co-transfected into HEK293 cells. 48 hours after transfection the cells were harvested, the soluble protein extracted via dounce homogenization, the lysate clarified via centrifugation (30 min at 25k rcf) followed by filtration (0.45 µm filter). The clarified cell lystate was allowed to flow through a Strep-Tactin XT Superflow column and the flowthrough was collected for analysis. The bound protein was washed extensively with wash buffer (100 mM Tris-HCl pH 8.0, 150 mM NaCl, 1 mM EDTA) and eluted in wash buffer containing 50 mM D-biotin. The protein samples were resolved on a 10-20% gradient SDS-PAGE gel (Invitrogen) and stained with the Pierce Silver Stain for MS kit (Invitrogen).

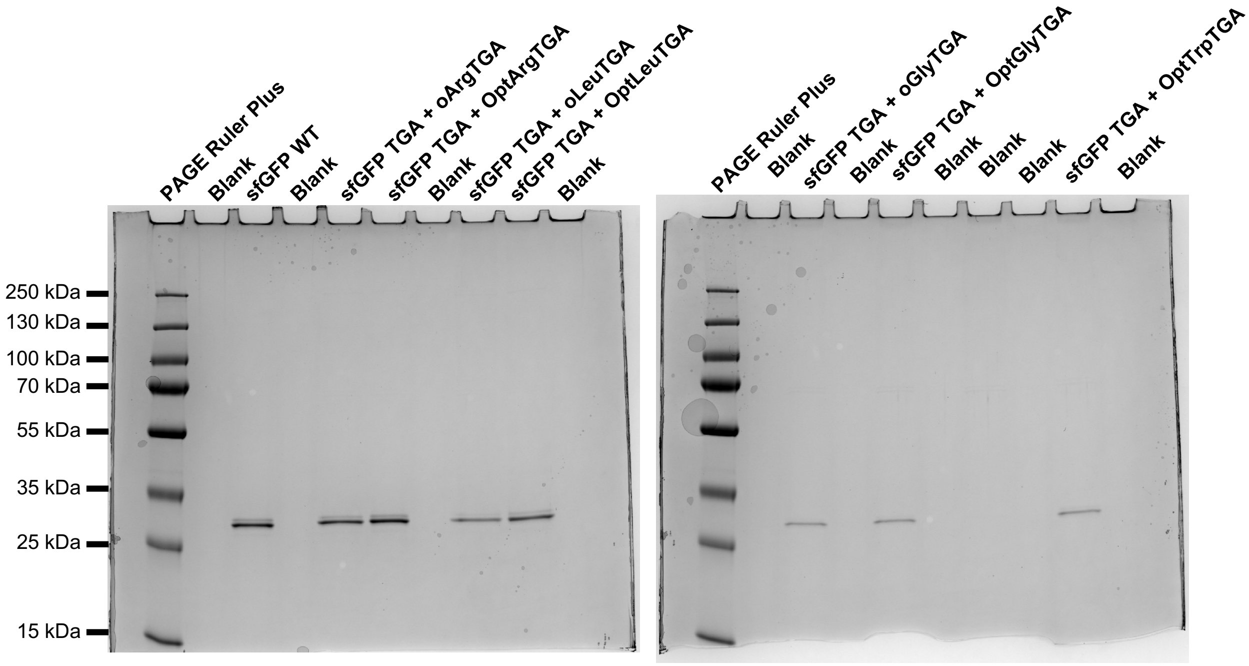


**Supplemental Figure 11. Uncropped gel images of gels shown in Figure 6C**

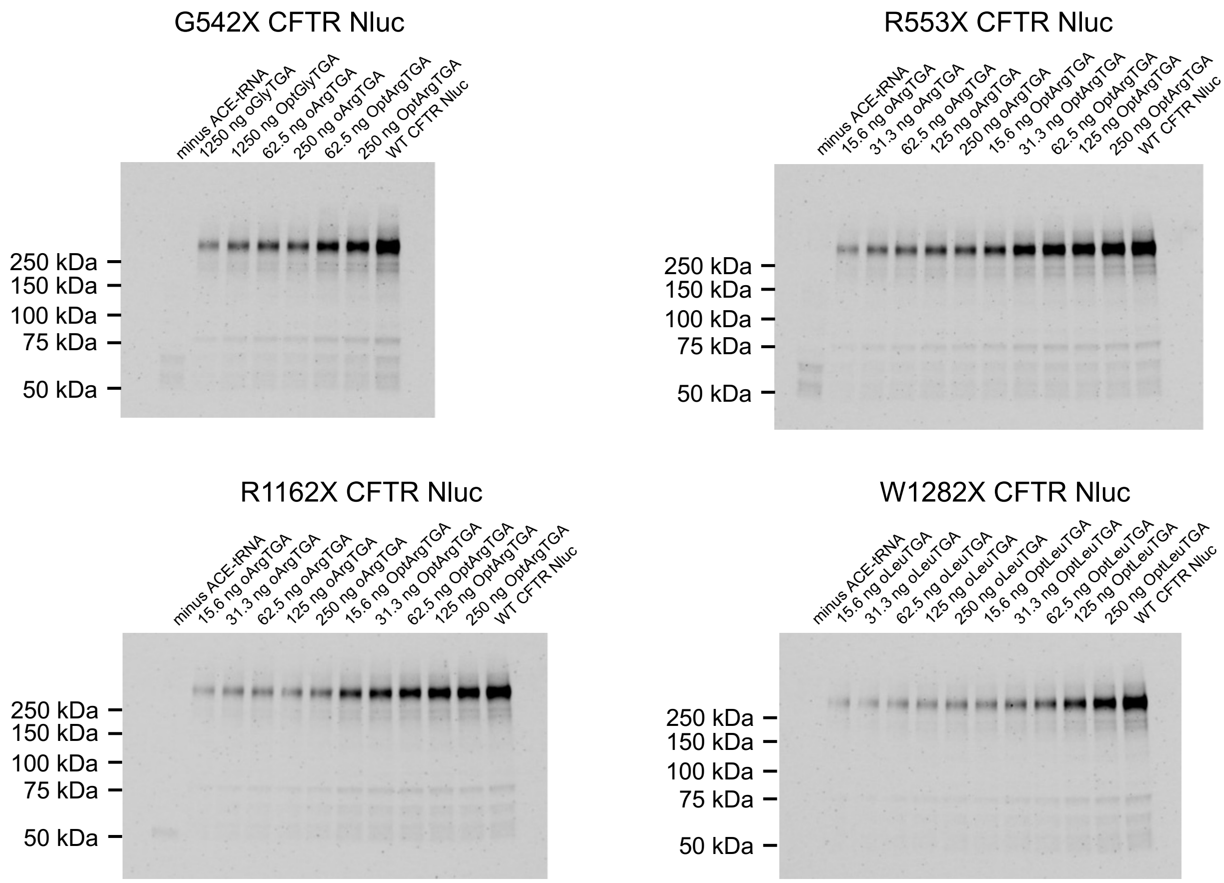


**Supplemental Figure 12. Uncropped Nluc in-gel luminescence images of gels shown in Figure 7B**

| ACE-tRNA expression cassette | $Sup_{max}$ (arbitrary units) | $DD_{50}$ (ng/μL) |
|---|---|---|
| Original ArgTGA | 4000 ± 150 | 11.3 ± 1.2 |
| Optimized ArgTGA | 5300 ± 150 | 5.0 ± 0.6 |
| Original LeuTGA | 1300 ± 30 | 5.6 ± 0.4 |
| Optimized LeuTGA | 1330 ± 35 | 1.6 ± 0.2 |
| Original GlyTGA | 560 ± 30 | 23 ± 3 |
| Optimized GlyTGA | 590 ± 35 | 16 ± 2 |
| Original TrpTGA | 66 ± 4 | 23 ± 3 |
| Optimized TrpTGA | 240 ± 20 | 24 ± 5 |

Supplemental Table 1. **Fit data for plots displayed in Figure 5.** Curves were fit as outlined in the legend for Supplemental Figure 7.

| | Arg (%) | Asn (%) | Leu (%) | Gly (%) |
|---|---|---|---|---|
| Original ArgTGA | 99.46 | 0.54 | 0.00 | 0.00 |
| Optimized ArgTGA | 99.85 | 0.15 | 0.00 | 0.00 |
| Original LeuTGA | 0.00 | 2.25 | 97.75 | 0.00 |
| Optimized LeuTGA | 0.19 | 0.45 | 99.29 | 0.07 |
| Optimized GlyTGA | 0.90 | 1.92 | 0.00 | 97.18 |
| Optimized TrpTGA | 99.99 | 0.01 | 0.00 | 0.00 |

Supplemental Table 2. **Percent of each amino acid incorporated at position 150 in sfGFP-UGA-150 as determined by mass spectrometry.** Percent values of incorporation for each amino acid for data shown in Figure 6.

**Genbank sequence for 5' HTCS vector**

```
LOCUS       5prime_pNanoRePo       7167 bp DNA     circular SYN 29-MAR-2023
DEFINITION  synthetic circular DNA
ACCESSION   .
VERSION     .
KEYWORDS    .
SOURCE      synthetic DNA construct
  ORGANISM  synthetic DNA construct
REFERENCE   1  (bases 1 to 7167)
  AUTHORS   .
  TITLE     Direct Submission
  JOURNAL   Exported Mar 29, 2023 from SnapGene 6.2.1
            https://www.snapgene.com
COMMENT     LOCUS       pcDNA3.1/Zeo(+)          5015 bp          DNA    SYN
            DEFINITION  pcDNA3.1/Zeo(+)
            ACCESSION
            KEYWORDS
            SOURCE
              ORGANISM  other sequences; artificial sequences; vectors.
FEATURES             Location/Qualifiers
     source          1..7167
                     /mol_type="other DNA"
                     /organism="synthetic DNA construct"
     source          5253..5325
                     /lab_host="E.coli"
                     /db_xref="taxon:66198"
                     /organism="Cloning vector pUC57"
     promoter        36..435
                     /label=UbC promoter
                     /note="human ubiquitin C promoter"
     promoter        445..463
                     /label=T7 promoter
                     /note="promoter for bacteriophage T7 RNA polymerase"
     regulatory      491..500
                     /label=Kozak sequence
                     /note="vertebrate consensus sequence for strong initiation
                     of translation (Kozak, 1987)"
                     /regulatory_class="other"
     CDS             497..2146
                     /codon_start=1
                     /product="firefly luciferase reporter gene luc2 (Photinus
                     pyralis) and is designed for high expression and reduced
                     anomalous transcription. This sequence was engineered with
                     fewer consensus regulatory sequences and has been codon
                     optimized for mammalian expression."
                     /label=Fluc2 (From Promega pGL4.10[luc2] vector)
                     /note="synthetic luc2 version of the luciferase gene"
                     /translation="MEDAKNIKKGPAPFYPLEDGTAGEQLHKAMKRYALVPGTIAFTDA
```

HIEVDITYAEYFEMSVRLAEAMKRYGLNTNHRIVVCSENSLQFFMPVLGALFIGVAVAP

ANDIYNERELLNSMGISQPTVVFVSKKGLQKILNVQKKLPIIQKIIIMDSKTDYQGFQS

MYTFVTSHLPPGFNEYDFVPESFDRDKTIALIMNSSGSTGLPKGVALPHRTACVRFSHA

RDPIFGNQIIPDTAILSVVPFHHGFGMFTTLGYLICGFRVVLMYRFEEELFLRSLQDYK

IQSALLVPTLFSFFAKSTLIDKYDLSNLHEIASGGAPLSKEVGEAVAKRFHLPGIRQGY

GLTETTSAILITPEGDDKPGAVGKVVPFFEAKVVDLDTGKTLGVNQRGELCVRGPMIMS

GYVNNPEATNALIDKDGWLHSGDIAYWDEDEHFFIVDRLKSLIKYKGYQVAPAELESIL

LQHPNIFDAGVAGLPDDDAGELPAAVVVLEHGKTMTEKEIVDYVASQVTTAKKLRGGVV
FVDEVPKGLTGKLDARKIREILIKAKKGGKIAV"

misc_feature 2162..2281
/label=SV40 poly(A) signal
enhancer 2351..2730
/label=CMV immediate early enhancer
/note="human cytomegalovirus immediate early enhancer"
promoter 2731..2934
/label=CMV promoter
/note="human cytomegalovirus (CMV) immediate early
promoter"
promoter 2979..2997
/label=T7 promoter
/note="promoter for bacteriophage T7 RNA polymerase"
regulatory 3033..3042
/label=Kozak sequence
/note="vertebrate consensus sequence for strong initiation
of translation (Kozak, 1987)"
/regulatory_class="other"
CDS 3039..3556
/codon_start=1
/product="NanoLuc(R) luciferase"
/label=Nluc
/note="human codon-optimized"
/translation="MVFTLEDFVGDWRQTAGYNLDQVLEQGGVSSLFQNLGVSVTPIQR

IVLSGENGLKIDIHVIIPYEGLSGDQMGQIEKIFKVVYPVDDHHFKVILHYGTLVIDGV

TPNMIDYFGRPYEGIAVFDGKKITVTGTLWNGNKIIDERLINPDGSLLFRVTING*VTG
WRLCERILA"

primer_bind 3447..3465
/label=NanoLueckFseq (121)
misc_feature 3516..3518
/label=PTC
misc_feature 3616..3747
/label=SV40 poly(A) signal

misc_feature    3785..3791
/label=SapI
promoter    3816..3846
/label=lac UV5 promoter
/note="E. coli lac promoter with an ""up"" mutation"
CDS    3900..4559
/codon_start=1
/gene="cat"
/product="chloramphenicol acetyltransferase"
/label=CmR
/note="confers resistance to chloramphenicol"
/translation="MEKKITGYTTVDISQWHRKEHFEAFQSVAQCTYNQTVQLDITAFL

KTVKKNKHKFYPAFIHILARLMNAHPEFRMAMKDGELVIWDSVHPCYTVFHEQTETFSS

LWSEYHDDFRQFLHIYSQDVACYGENLAYFPKGFIENMFFVSANPWVSFTSFDLNVANM

DNFFAPVFTMGKYYTQGDKVLMPLAIQVHHAVCDGFHVGRMLNELQQYCDEWQGGA"
CDS    4901..5206
/codon_start=1
/gene="ccdB"
/product="CcdB, a bacterial toxin that poisons DNA gyrase"
/label=ccdB
/note="Plasmids containing the ccdB gene cannot be
propagated in standard E. coli strains."
/translation="MQFKVYTYKRESRYRLFVDVQSDIIDTPGRRMVIPLASARLLSDK

VSRELYPVVHIGDESWRMMTTDMASVPVSVIGEEVADLSHRENDIKNAINLMFWGI"
misc_feature    5245..5251
/label=SapI
gene    complement(5253..5325)
/gene="lacZ"
/label=lacZ
misc_feature    5253..5325
/label=tRNA Arg
variation    5326..5337
/label=3' term signal
/note="3' term signal"
rep_origin    complement(5420..6008)
/direction=LEFT
/label=pBR322 ori
/note="high-copy-number ColE1/pMB1/pBR322/pUC origin of
replication"
CDS    complement(6179..7039)
/codon_start=1
/gene="bla"
/product="beta-lactamase"
/label=AmpR
/note="confers resistance to ampicillin, carbenicillin, and
related antibiotics"
/translation="MSIQHFRVALIPFFAAFCLPVFAHPETLVKVKDAEDQLGARVGYI

ELDLNSGKILESFRPEERFPMMSTFKVLLCGAVLSRIDAGQEQLGRRIHYSQNDLVEYS

PVTEKHLTDGMTVRELCSAAITMSDNTAANLLLTTIGGPKELTAFLHNMGDHVTRLDRW

EPELNEAIPNDERDTTMPVAMATTLRKLLTGELLTLASRQQLIDWMEADKVAGPLLRSA

LPAGWFIADKSGAGERGSRGIIAALGPDGKPSRIVVIYTTGSQATMDERNRQIAEIGAS
LIKHW"

```
     promoter        complement(7040..7144)
                     /gene="bla"
                     /label=AmpR promoter
ORIGIN
        1 ctgacgtcga cggatcggga gggttaatta cgcgtggcct ccgcgccggg ttttggcgcc
       61 tcccgcgggc gcccccctcc tcacggcgag cgctgccacg tcagacgaag ggcgcagcga
      121 gcgtcctgat ccttccgccc ggacgctcag gacagcggcc cgctgctcat aagactcggc
      181 cttagaaccc cagtatcagc agaaggacat tttaggacgg gacttgggtg actctagggc
      241 actggttttc tttccagaga gcggaacagg cgaggaaaag tagtcccttc tcggcgattc
      301 tgcggaggga tctccgtggg gcggtgaacg ccgatgatta tataaggacg cgccgggtgt
      361 ggcacagcta gttccgtcgc agccgggatt tgggtcgcgg ttcttgtttg tggatcgctg
      421 tgatcgtcac ttggttatcg aaattaatac gactcactat agggagaccc aagctgctag
      481 ccacgtggcc gccaccatgg aagatgccaa aaacattaag aagggcccag cgccattcta
      541 cccactcgag gacgggaccg ccggcgagca gctgcacaaa gccatgaagc gctacgccct
      601 ggtgcccggc accatcgcct ttaccgacgc acatatcgag gtggacatta cctacgccga
      661 gtacttcgag atgagcgttc ggctggcaga agctatgaag cgctatgggc tgaatacaaa
      721 ccatcggatc gtggtgtgca gcgagaatag cttgcagttc ttcatgcccg tgttgggtgc
      781 cctgttcatc ggtgtggctg tggccccagc taacgacatc tacaacgagc gcgagctgct
      841 gaacagcatg ggcatcagcc agcccaccgt cgtattcgtg agcaagaaag ggctgcaaaa
      901 gatcctcaac gtgcaaaaga agctaccgat catacaaaag atcatcatca tggatagcaa
      961 gaccgactac cagggcttcc aaagcatgta caccttcgtg acttcccatt tgccacccgg
     1021 cttcaacgag tacgacttcg tgcccgagag cttcgaccgg gacaaaacca tcgccctgat
     1081 catgaacagt agtggcagta ccggattgcc caagggcgta gccctaccgc accgcaccgc
     1141 ttgtgtccga ttcagtcatg cccgcgaccc catcttcggc aaccagatca tccccgacac
     1201 cgctatcctc agcgtggtgc catttcacca cggcttcggc atgttcacca cgctgggcta
     1261 cttgatctgc ggctttcggg tcgtgctcat gtaccgcttc gaggaggagc tattcttgcg
     1321 cagcttgcaa gactataaga ttcaatctgc cctgctggtg cccacactat ttagcttctt
     1381 cgctaagagc actctcatcg acaagtacga cctaagcaac ttgcacgaga tcgccagcgg
     1441 cggggcgccg ctcagcaagg aggtaggtga ggccgtggcc aaacgcttcc acctaccagg
     1501 catccgccag ggctacggcc tgacagaaac aaccagcgcc attctgatca cccccgaagg
     1561 ggacgacaag cctggcgcag taggcaaggt ggtgcccttc ttcgaggcta aggtggtgga
     1621 cttggacacc ggtaagacac tgggtgtgaa ccagcgcggc gagctgtgcg tccgtggccc
     1681 catgatcatg agcggctacg ttaacaaccc cgaggctaca aacgctctca tcgacaagga
     1741 cggctggctg cacagcggcg acatcgccta ctgggacgag gacgagcact tcttcatcgt
     1801 ggaccggctg aaaagcctga tcaaatacaa gggctaccag gtagccccag ccgaactgga
     1861 gagcatcctg ctgcaacacc ccaacatctt cgacgccggg gtcgccggcc tgcccgacga
     1921 cgatgccggc gagctgcccg ccgcagtcgt cgtgctggaa cacggtaaaa ccatgaccga
     1981 gaaggagatc gtggactatg tggccagcca ggttacaacc gccaagaagc tgcgcggtgg
     2041 tgttgtgttc gtggacgagg tgcctaaagg actgaccggc aagttggacg cccgcaagat
     2101 ccgcgagatt ctcattaagg ccaagaaggg cggcaagatc gccgtgtaat gatagggtac
     2161 cttgtttatt gcagcttata atggttacaa ataaagcaat agcatcacaa atttcacaaa
     2221 taaagcattt ttttcactgc attctagttg tggtttgtcc aaactcatca atgtatctta
```

2281 tgcatgaaga atctgcttag ggttaggcgt tttgcgctgc ttcgcgatgt acgggccaga
2341 tatacgcgtt gacattgatt attgactagt tattaatagt aatcaattac ggggtcatta
2401 gttcatagcc catatatgga gttccgcgtt acataactta cggtaaatgg cccgcctggc
2461 tgaccgccca acgacccccg cccattgacg tcaataatga cgtatgttcc catagtaacg
2521 ccaataggga ctttccattg acgtcaatgg gtggagtatt tacggtaaac tgcccacttg
2581 gcagtacatc aagtgtatca tatgccaagt acgcccccta ttgacgtcaa tgacggtaaa
2641 tggcccgcct ggcattatgc ccagtacatg accttatggg actttcctac ttggcagtac
2701 atctacgtat tagtcatcgc tattaccatg gtgatgcggt tttggcagta catcaatggg
2761 cgtggatagc ggtttgactc acggggattt ccaagtctcc accccattga cgtcaatggg
2821 agtttgtttt ggcaccaaaa tcaacgggac tttccaaaat gtcgtaacaa ctccgcccca
2881 ttgacgcaaa tgggcggtag gcgtgtacgg tgggaggtct atataagcag agctctctgg
2941 ctaactagag aacccactgc ttactggctt atcgaaatta atacgactca ctatagggag
3001 acccaagctg gctagcgttt aaacttaagc ttgccaccat ggtattcaca ctcgaagatt
3061 tcgttgggga ctggcgacag acagccggct acaacctgga ccaagtcctt gaacagggag
3121 gtgtgtccag tttgtttcag aatctcgggg tgtccgtaac tccgatccaa aggattgtcc
3181 tgagcggtga aaatgggctg aagatcgaca tccatgtcat catcccgtat gaaggtctga
3241 gcggcgacca aatgggccag atcgaaaaaa tttttaaggt ggtgtaccct gtggatgatc
3301 atcactttaa ggtgatcctg cactatggca cactggtaat cgacggggtt acgccgaaca
3361 tgatcgacta tttcggacgg ccgtatgaag gcatcgccgt gttcgacggc aaaaagatca
3421 ctgtaacagg gaccctgtgg aacggcaaca aaattatcga cgagcgcctg atcaaccccg
3481 acggctccct gctgttccga gtaaccatca acggatgagt gaccggctgg cggctgtgcg
3541 aacgcattct ggcgtaagaa ttctaagggc gaattctgca gatatccagc acagtggcgg
3601 ccgctcgagt ctagattgtt tattgcagct tataatggtt acaaataaag caatagcatc
3661 acaaatttca caaataaagc atttttttca ctgcattcta gttgtggttt gtccaaactc
3721 atcaatgtat cttatcatgt ctggatcgct agagggcctt cctaatacga ctcactatag
3781 tttcgaagag cgcggccgca ttaggcaccc caggctttac actttatgct tccggctcgt
3841 ataatgtgtg gattttgagt taggatccgt cgagattttc aggagctaag gaagctaaaa
3901 tggagaaaaa aatcactgga tataccaccg ttgatatatc ccaatggcat cgtaaagaac
3961 attttgaggc atttcagtca gttgctcaat gtacctataa ccagaccgtt cagctggata
4021 ttacggcctt tttaaagacc gtaaagaaaa ataagcacaa gttttatccg gcctttattc
4081 acattcttgc ccgcctgatg aatgctcatc cggaattccg tatggcaatg aaagacggtg
4141 agctggtgat atgggatagt gttcaccctt gttacaccgt tttccatgag caaactgaaa
4201 cgttttcatc gctctggagt gaataccacg acgatttccg gcagtttcta cacatatatt
4261 cgcaagatgt ggcgtgttac ggtgaaaacc tggcctattt ccctaaaggg tttattgaga
4321 atatgttttt cgtctcagcc aatccctggg tgagtttcac cagttttgat ttaaacgtgg
4381 ccaatatgga caacttcttc gcccccgttt tcaccatggg caaatattat acgcaaggcg
4441 acaaggtgct gatgccgctg gcgattcagg ttcatcatgc cgtttgtgat ggcttccatg
4501 tcggcagaat gcttaatgaa ttacaacagt actgcgatga gtggcagggc ggggcgtaaa
4561 gatctggatc cggcttacta aaagccagat aacagtatgc gtatttgcgc gctgattttt
4621 gcggtataag aatatatact gatatgtata cccgaagtat gtcaaaaaga ggtatgctat
4681 gaagcagcgt attacagtga cagttgacag cgacagctat cagttgctca aggcatatat
4741 gatgtcaata tctccggtct ggtaagcaca accatgcaga atgaagcccg tcgtctgcgt
4801 gccgaacgct ggaaagcgga aaatcaggaa gggatggctg aggtcgcccg gtttattgaa
4861 atgaacggct cttttgctga cgagaacagg ggctggtgaa atgcagttta aggtttacac
4921 ctataaaaga gagagccgtt atcgtctgtt tgtggatgta cagagtgata ttattgacac
4981 gcccgggcga cggatggtga tccccctggc cagtgcacgt ctgctgtcag ataaagtctc
5041 ccgtgaactt tacccggtgg tgcatatcgg ggatgaaagc tggcgcatga tgaccaccga
5101 tatggccagt gtgcccgtct ccgttatcgg ggaagaagtg gctgatctca gccaccgcga
5161 aaatgacatc aaaaacgcca ttaacctgat gttctgggga atataaatgt caggctccct
5221 tatacacagc cagtctgcag ggaagctctt cgggctctgt ggcgcaatgg atagcgcatt
5281 ggacttcaaa ttcaaaggtt gtgggttcga gtcccaccag agtcggtcct ttttttgctt

5341 tagtgagggt taattcaggc atgtgagcaa aaggccagca aaaggccagg aaccgtaaaa
5401 aggccgcgtt gctggcgttt ttccataggc tccgcccccc tgacgagcat cacaaaaatc
5461 gacgctcaag tcagaggtgg cgaaacccga caggactata aagataccag gcgtttcccc
5521 ctggaagctc cctcgtgcgc tctcctgttc cgaccctgcc gcttaccgga tacctgtccg
5581 cctttctccc ttcgggaagc gtggcgcttt ctcaatgctc acgctgtagg tatctcagtt
5641 cggtgtaggt cgttcgctcc aagctgggct gtgtgcacga accccccgtt cagcccgacc
5701 gctgcgcctt atccggtaac tatcgtcttg agtccaaccc ggtaagacac gacttatcgc
5761 cactggcagc agccactggt aacaggatta gcagagcgag gtatgtaggc ggtgctacag
5821 agttcttgaa gtggtggcct aactacggct acactagaag gacagtattt ggtatctgcg
5881 ctctgctgaa gccagttacc ttcggaaaaa gagttggtag ctcttgatcc ggcaaacaaa
5941 ccaccgctac cagcggtggt ttttttgttt gcaagcagca gattacgcgc agaaaaaaag
6001 gatctcaaga agatcctttg atcttttcta cggggtctga cgctcagtgg aacgaaaact
6061 cacgttaagg gattttggtc atgagattat caaaaaggat cttcacctag atccttttaa
6121 attaaaaatg aagttttaaa tcaatctaaa gtatatatga gtaaacttgg tctgacagtt
6181 accaatgctt aatcagtgag gcacctatct cagcgatctg tctatttcgt tcatccatag
6241 ttgcctgact ccccgtcgtg tagataacta cgatacggga gggcttacca tctggcccca
6301 gtgctgcaat gataccgcga gacccacgct caccggctcc agatttatca gcaataaacc
6361 agccagccgg aagggccgag cgcagaagtg gtcctgcaac tttatccgcc tccatccagt
6421 ctattaattg ttgccgggaa gctagagtaa gtagttcgcc agttaatagt ttgcgcaacg
6481 ttgttgccat tgctacaggc atcgtggtgt cacgctcgtc gtttggtatg gcttcattca
6541 gctccggttc ccaacgatca aggcgagtta catgatcccc catgttgtgc aaaaaagcgg
6601 ttagctcctt cggtcctccg atcgttgtca gaagtaagtt ggccgcagtg ttatcactca
6661 tggttatggc agcactgcat aattctctta ctgtcatgcc atccgtaaga tgcttttctg
6721 tgactggtga gtactcaacc aagtcattct gagaatagtg tatgcggcga ccgagttgct
6781 cttgcccggc gtcaatacgg gataataccg cgccacatag cagaacttta aaagtgctca
6841 tcattggaaa acgttcttcg gggcgaaaac tctcaaggat cttaccgctg ttgagatcca
6901 gttcgatgta acccactcgt gcacccaact gatcttcagc atcttttact ttcaccagcg
6961 tttctgggtg agcaaaaaca ggaaggcaaa atgccgcaaa aaagggaata agggcgacac
7021 ggaaatgttg aatactcata ctcttccttt ttcaatatta ttgaagcatt tatcagggtt
7081 attgtctcat gagcggatac atatttgaat gtatttagaa aaataaacaa ataggggttc
7141 cgcgcacatt tccccgaaaa gtgccac
//

**Genbank sequence for 3' HTCS vector**

```
LOCUS       3prime_pNanoRePo        7211 bp DNA     circular SYN 29-MAR-2023
DEFINITION  synthetic circular DNA
ACCESSION   .
VERSION     .
KEYWORDS    .
SOURCE      synthetic DNA construct
  ORGANISM  synthetic DNA construct
REFERENCE   1  (bases 1 to 7211)
  AUTHORS   .
  TITLE     Direct Submission
  JOURNAL   Exported Mar 29, 2023 from SnapGene 6.2.1
            https://www.snapgene.com
COMMENT     LOCUS       pcDNA3.1/Zeo(+)          5015 bp       DNA     SYN
            DEFINITION  pcDNA3.1/Zeo(+)
            ACCESSION
            KEYWORDS
            SOURCE
              ORGANISM  other sequences; artificial sequences; vectors.
FEATURES             Location/Qualifiers
     source          1..7211
                     /mol_type="other DNA"
                     /organism="synthetic DNA construct"
     source          3836..3908
                     /lab_host="E.coli"
                     /db_xref="taxon:66198"
                     /organism="Cloning vector pUC57"
     promoter        36..435
                     /label=UbC promoter
                     /note="human ubiquitin C promoter"
     promoter        445..463
                     /label=T7 promoter
                     /note="promoter for bacteriophage T7 RNA polymerase"
     regulatory      491..500
                     /label=Kozak sequence
                     /note="vertebrate consensus sequence for strong initiation
                     of translation (Kozak, 1987)"
                     /regulatory_class="other"
     CDS             497..2146
                     /codon_start=1
                     /product="firefly luciferase reporter gene luc2 (Photinus
                     pyralis) and is designed for high expression and reduced
                     anomalous transcription. This sequence was engineered with
                     fewer consensus regulatory sequences and has been codon
                     optimized for mammalian expression."
                     /label=Fluc2 (From Promega pGL4.10[luc2] vector)
                     /note="synthetic luc2 version of the luciferase gene"
                     /translation="MEDAKNIKKGPAPFYPLEDGTAGEQLHKAMKRYALVPGTIAFTDA

HIEVDITYAEYFEMSVRLAEAMKRYGLNTNHRIVVCSENSLQFFMPVLGALFIGVAVAP
```

ANDIYNERELLNSMGISQPTVVFVSKKGLQKILNVQKKLPIIQKIIIMDSKTDYQGFQS

MYTFVTSHLPPGFNEYDFVPESFDRDKTIALIMNSSGSTGLPKGVALPHRTACVRFSHA

RDPIFGNQIIPDTAILSVVPFHHGFGMFTTLGYLICGFRVVLMYRFEEELFLRSLQDYK

IQSALLVPTLFSFFAKSTLIDKYDLSNLHEIASGGAPLSKEVGEAVAKRFHLPGIRQGY

GLTETTSAILITPEGDDKPGAVGKVVPFFEAKVVDLDTGKTLGVNQRGELCVRGPMIMS

GYVNNPEATNALIDKDGWLHSGDIAYWDEDEHFFIVDRLKSLIKYKGYQVAPAELESIL

LQHPNIFDAGVAGLPDDDAGELPAAVVVLEHGKTMTEKEIVDYVASQVTTAKKLRGGVV
FVDEVPKGLTGKLDARKIREILIKAKKGGKIAV"
misc_feature 2162..2281
/label=SV40 poly(A) signal
enhancer 2351..2730
/label=CMV immediate early enhancer
/note="human cytomegalovirus immediate early enhancer"
promoter 2731..2934
/label=CMV promoter
/note="human cytomegalovirus (CMV) immediate early
promoter"
promoter 2979..2997
/label=T7 promoter
/note="promoter for bacteriophage T7 RNA polymerase"
regulatory 3033..3042
/label=Kozak sequence
/note="vertebrate consensus sequence for strong initiation
of translation (Kozak, 1987)"
/regulatory_class="other"
CDS 3039..3556
/codon_start=1
/product="NanoLuc(R) luciferase"
/label=Nluc
/note="human codon-optimized"
/translation="MVFTLEDFVGDWRQTAGYNLDQVLEQGGVSSLFQNLGVSVTPIQR

IVLSGENGLKIDIHVIIPYEGLSGDQMGQIEKIFKVVYPVDDHHFKVILHYGTLVIDGV

TPNMIDYFGRPYEGIAVFDGKKITVTGTLWNGNKIIDERLINPDGSLLFRVTING*VTG
WRLCERILA"
primer_bind 3447..3465
/label=NanoLueckFseq (121)
misc_feature 3516..3518
/label=PTC
misc_feature 3616..3747
/label=SV40 poly(A) signal
5'UTR 3763..3835
/label=5' human Tyr-tRNA leader

/note="5' human Tyr-tRNA leader"
gene complement(3836..3908)
/gene="lacZ"
/label=lacZ
misc_feature 3836..3908
/label=tRNA Arg
misc_feature 3911..3916
/label=BbsI
promoter 3941..3971
/label=lac UV5 promoter
/note="E. coli lac promoter with an ""up"" mutation"
CDS 4025..4684
/codon_start=1
/gene="cat"
/product="chloramphenicol acetyltransferase"
/label=CmR
/note="confers resistance to chloramphenicol"
/translation="MEKKITGYTTVDISQWHRKEHFEAFQSVAQCTYNQTVQLDITAFL

KTVKKNKHKFYPAFIHILARLMNAHPEFRMAMKDGELVIWDSVHPCYTVFHEQTETFSS

LWSEYHDDFRQFLHIYSQDVACYGENLAYFPKGFIENMFFVSANPWVSFTSFDLNVANM

DNFFAPVFTMGKYYTQGDKVLMPLAIQVHHAVCDGFHVGRMLNELQQYCDEWQGGA"
CDS 5026..5331
/codon_start=1
/gene="ccdB"
/product="CcdB, a bacterial toxin that poisons DNA gyrase"
/label=ccdB
/note="Plasmids containing the ccdB gene cannot be
propagated in standard E. coli strains."
/translation="MQFKVYTYKRESRYRLFVDVQSDIIDTPGRRMVIPLASARLLSDK

VSRELYPVVHIGDESWRMMTTDMASVPVSVIGEEVADLSHRENDIKNAINLMFWGI"
misc_feature 5370..5375
/label=BbsI
rep_origin complement(5464..6052)
/direction=LEFT
/label=pBR322 ori
/note="high-copy-number ColE1/pMB1/pBR322/pUC origin of
replication"
CDS complement(6223..7083)
/codon_start=1
/gene="bla"
/product="beta-lactamase"
/label=AmpR
/note="confers resistance to ampicillin, carbenicillin, and
related antibiotics"
/translation="MSIQHFRVALIPFFAAFCLPVFAHPETLVKVKDAEDQLGARVGYI

ELDLNSGKILESFRPEERFPMMSTFKVLLCGAVLSRIDAGQEQLGRRIHYSQNDLVEYS

PVTEKHLTDGMTVRELCSAAITMSDNTAANLLLTTIGGPKELTAFLHNMGDHVTRLDRW

EPELNEAIPNDERDTTMPVAMATTLRKLLTGELLTLASRQQLIDWMEADKVAGPLLRSA

LPAGWFIADKSGAGERGSRGIIAALGPDGKPSRIVVIYTTGSQATMDERNRQIAEIGAS
LIKHW"
promoter complement(7084..7188)
/gene="bla"
/label=AmpR promoter
ORIGIN
1 ctgacgtcga cggatcggga gggttaatta cgcgtggcct ccgcgccggg ttttggcgcc
61 tcccgcgggc gcccccctcc tcacggcgag cgctgccacg tcagacgaag ggcgcagcga
121 gcgtcctgat ccttccgccc ggacgctcag gacagcggcc cgctgctcat aagactcggc
181 cttagaaccc cagtatcagc agaaggacat tttaggacgg gacttgggtg actctagggc
241 actggttttc tttccagaga gcggaacagg cgaggaaaag tagtcccttc tcggcgattc
301 tgcggaggga tctccgtggg gcggtgaacg ccgatgatta tataaggacg cgccgggtgt
361 ggcacagcta gttccgtcgc agccgggatt tgggtcgcgg ttcttgtttg tggatcgctg
421 tgatcgtcac ttggttatcg aaattaatac gactcactat agggagaccc aagctgctag
481 ccacgtggcc gccaccatgg aagatgccaa aaacattaag aagggcccag cgccattcta
541 cccactcgag gacgggaccg ccggcgagca gctgcacaaa gccatgaagc gctacgccct
601 ggtgcccggc accatcgcct ttaccgacgc acatatcgag gtggacatta cctacgccga
661 gtacttcgag atgagcgttc ggctggcaga agctatgaag cgctatgggc tgaatacaaa
721 ccatcggatc gtggtgtgca gcgagaatag cttgcagttc ttcatgcccg tgttgggtgc
781 cctgttcatc ggtgtggctg tggccccagc taacgacatc tacaacgagc gcgagctgct
841 gaacagcatg ggcatcagcc agcccaccgt cgtattcgtg agcaagaaag ggctgcaaaa
901 gatcctcaac gtgcaaaaga agctaccgat catacaaaag atcatcatca tggatagcaa
961 gaccgactac cagggcttcc aaagcatgta caccttcgtg acttcccatt tgccacccgg
1021 cttcaacgag tacgacttcg tgcccgagag cttcgaccgg gacaaaacca tcgccctgat
1081 catgaacagt agtggcagta ccggattgcc caagggcgta gccctaccgc accgcaccgc
1141 ttgtgtccga ttcagtcatg cccgcgaccc catcttcggc aaccagatca tccccgacac
1201 cgctatcctc agcgtggtgc catttcacca cggcttcggc atgttcacca cgctgggcta
1261 cttgatctgc ggctttcggg tcgtgctcat gtaccgcttc gaggaggagc tattcttgcg
1321 cagcttgcaa gactataaga ttcaatctgc cctgctggtg cccacactat ttagcttctt
1381 cgctaagagc actctcatcg acaagtacga cctaagcaac ttgcacgaga tcgccagcgg
1441 cggggcgccg ctcagcaagg aggtaggtga ggccgtggcc aaacgcttcc acctaccagg
1501 catccgccag ggctacggcc tgacagaaac aaccagcgcc attctgatca cccccgaagg
1561 ggacgacaag cctggcgcag taggcaaggt ggtgcccttc ttcgaggcta aggtggtgga
1621 cttggacacc ggtaagacac tgggtgtgaa ccagcgcggc gagctgtgcg tccgtggccc
1681 catgatcatg agcggctacg ttaacaaccc cgaggctaca aacgctctca tcgacaagga
1741 cggctggctg cacagcggcg acatcgccta ctgggacgag gacgagcact tcttcatcgt
1801 ggaccggctg aagagcctga tcaaatacaa gggctaccag gtagccccag ccgaactgga
1861 gagcatcctg ctgcaacacc ccaacatctt cgacgccggg gtcgccggcc tgcccgacga
1921 cgatgccggc gagctgcccg ccgcagtcgt cgtgctggaa cacggtaaaa ccatgaccga
1981 gaaggagatc gtggactatg tggccagcca ggttacaacc gccaagaagc tgcgcggtgg
2041 tgttgtgttc gtggacgagg tgcctaaagg actgaccggc aagttggacg cccgcaagat
2101 ccgcgagatt ctcattaagg ccaagaaggg cggcaagatc gccgtgtaat gatagggtac
2161 cttgtttatt gcagcttata atggttacaa ataaagcaat agcatcacaa atttcacaaa
2221 taaagcattt ttttcactgc attctagttg tggtttgtcc aaactcatca atgtatctta
2281 tgcatgaaga atctgcttag ggttaggcgt tttgcgctgc ttcgcgatgt acgggccaga
2341 tatacgcgtt gacattgatt attgactagt tattaatagt aatcaattac ggggtcatta

2401 gttcatagcc catatatgga gttccgcgtt acataactta cggtaaatgg cccgcctggc
2461 tgaccgccca acgacccccg cccattgacg tcaataatga cgtatgttcc catagtaacg
2521 ccaataggga ctttccattg acgtcaatgg gtggagtatt tacggtaaac tgcccacttg
2581 gcagtacatc aagtgtatca tatgccaagt acgcccccta ttgacgtcaa tgacggtaaa
2641 tggcccgcct ggcattatgc ccagtacatg accttatggg actttcctac ttggcagtac
2701 atctacgtat tagtcatcgc tattaccatg gtgatgcggt tttggcagta catcaatggg
2761 cgtggatagc ggtttgactc acggggattt ccaagtctcc accccattga cgtcaatggg
2821 agtttgtttt ggcaccaaaa tcaacgggac tttccaaaat gtcgtaacaa ctccgcccca
2881 ttgacgcaaa tgggcggtag gcgtgtacgg tgggaggtct atataagcag agctctctgg
2941 ctaactagag aacccactgc ttactggctt atcgaaatta atacgactca ctatagggag
3001 acccaagctg gctagcgttt aaacttaagc ttgccaccat ggtattcaca ctcgaagatt
3061 tcgttgggga ctggcgacag acagccggct acaacctgga ccaagtcctt gaacagggag
3121 gtgtgtccag tttgtttcag aatctcgggg tgtccgtaac tccgatccaa aggattgtcc
3181 tgagcggtga aaatgggctg aagatcgaca tccatgtcat catcccgtat gaaggtctga
3241 gcggcgacca aatgggccag atcgaaaaaa tttttaaggt ggtgtaccct gtggatgatc
3301 atcactttaa ggtgatcctg cactatggca cactggtaat cgacggggtt acgccgaaca
3361 tgatcgacta tttcggacgg ccgtatgaag gcatcgccgt gttcgacggc aaaaagatca
3421 ctgtaacagg gaccctgtgg aacggcaaca aaattatcga cgagcgcctg atcaaccccg
3481 acggctccct gctgttccga gtaaccatca acggatgagt gaccggctgg cggctgtgcg
3541 aacgcattct ggcgtaagaa ttctaagggc gaattctgca gatatccagc acagtggcgg
3601 ccgctcgagt ctagattgtt tattgcagct tataatggtt acaaataaag caatagcatc
3661 acaaatttca caaataaagc atttttttca ctgcattcta gttgtggttt gtccaaactc
3721 atcaatgtat cttatcatgt ctggatcgct agagggcctt cctaatacga ctcactatag
3781 agcgctccgg tttttctgtg ctgaacctca ggggacgccg acacacgtac acgtcggctc
3841 tgtggcgcaa tggatagcgc attggacttc aaattcaaag gttgtgggtt cgagtcccac
3901 cagagtcgta gtcttcgcgg ccgcattagg caccccaggc tttacacttt atgcttccgg
3961 ctcgtataat gtgtggattt tgagttagga tccgtcgaga ttttcaggag ctaaggaagc
4021 taaaatggag aaaaaaatca ctggatatac caccgttgat atatcccaat ggcatcgtaa
4081 agaacatttt gaggcatttc agtcagttgc tcaatgtacc tataaccaga ccgttcagct
4141 ggatattacg gcctttttaa agaccgtaaa gaaaaataag cacaagtttt atccggcctt
4201 tattcacatt cttgcccgcc tgatgaatgc tcatccggaa ttccgtatgg caatgaaaga
4261 cggtgagctg gtgatatggg atagtgttca cccttgttac accgttttcc atgagcaaac
4321 tgaaacgttt tcatcgctct ggagtgaata ccacgacgat ttccggcagt ttctacacat
4381 atattcgcaa gatgtggcgt gttacggtga aaacctggcc tatttcccta aagggtttat
4441 tgagaatatg tttttcgtct cagccaatcc ctgggtgagt ttcaccagtt ttgatttaaa
4501 cgtggccaat atggacaact tcttcgcccc cgttttcacc atgggcaaat attatacgca
4561 aggcgacaag gtgctgatgc cgctggcgat tcaggttcat catgccgttt gtgatggctt
4621 ccatgtcggc agaatgctta atgaattaca acagtactgc gatgagtggc agggcggggc
4681 gtaaagatct ggatccggct tactaaaagc cagataacag tatgcgtatt tgcgcgctga
4741 tttttgcggt ataagaatat atactgatat gtatacccga agtatgtcaa aaagaggtat
4801 gctatgaagc agcgtattac agtgacagtt gacagcgaca gctatcagtt gctcaaggca
4861 tatatgatgt caatatctcc ggtctggtaa gcacaaccat gcagaatgaa gcccgtcgtc
4921 tgcgtgccga acgctggaaa gcggaaaatc aggaagggat ggctgaggtc gcccggttta
4981 ttgaaatgaa cggctctttt gctgacgaga acaggggctg gtgaaatgca gtttaaggtt
5041 tacacctata aaagagagag ccgttatcgt ctgtttgtgg atgtacagag tgatattatt
5101 gacacgcccg ggcgacggat ggtgatcccc ctggccagtg cacgtctgct gtcagataaa
5161 gtctcccgtg aactttaccc ggtggtgcat atcggggatg aaagctggcg catgatgacc
5221 accgatatgg ccagtgtgcc cgtctccgtt atcggggaag aagtggctga tctcagccac
5281 cgcgaaaatg acatcaaaaa cgccattaac ctgatgttct ggggaatata aatgtcaggc
5341 tcccttatac acagccagtc tgcagggaag aagaccggtc cctttagtga gggttaattc
5401 aggcatgtga gcaaaaggcc agcaaaaggc caggaaccgt aaaaaggccg cgttgctggc

5461 gtttttccat aggctccgcc cccctgacga gcatcacaaa aatcgacgct caagtcagag
5521 gtggcgaaac ccgacaggac tataaagata ccaggcgttt ccccctggaa gctccctcgt
5581 gcgctctcct gttccgaccc tgccgcttac cggatacctg tccgcctttc tcccttcggg
5641 aagcgtggcg ctttctcaat gctcacgctg taggtatctc agttcggtgt aggtcgttcg
5701 ctccaagctg ggctgtgtgc acgaaccccc cgttcagccc gaccgctgcg ccttatccgg
5761 taactatcgt cttgagtcca acccggtaag acacgactta tcgccactgg cagcagccac
5821 tggtaacagg attagcagag cgaggtatgt aggcggtgct acagagttct tgaagtggtg
5881 gcctaactac ggctacacta gaaggacagt atttggtatc tgcgctctgc tgaagccagt
5941 taccttcgga aaaagagttg gtagctcttg atccggcaaa caaaccaccg ctaccagcgg
6001 tggttttttt gtttgcaagc agcagattac gcgcagaaaa aaaggatctc aagaagatcc
6061 tttgatcttt tctacggggt ctgacgctca gtggaacgaa aactcacgtt aagggatttt
6121 ggtcatgaga ttatcaaaaa ggatcttcac ctagatcctt ttaaattaaa aatgaagttt
6181 taaatcaatc taaagtatat atgagtaaac ttggtctgac agttaccaat gcttaatcag
6241 tgaggcacct atctcagcga tctgtctatt tcgttcatcc atagttgcct gactccccgt
6301 cgtgtagata actacgatac gggagggctt accatctggc cccagtgctg caatgatacc
6361 gcgagaccca cgctcaccgg ctccagattt atcagcaata aaccagccag ccggaagggc
6421 cgagcgcaga agtggtcctg caactttatc cgcctccatc cagtctatta attgttgccg
6481 ggaagctaga gtaagtagtt cgccagttaa tagtttgcgc aacgttgttg ccattgctac
6541 aggcatcgtg gtgtcacgct cgtcgtttgg tatggcttca ttcagctccg gttcccaacg
6601 atcaaggcga gttacatgat cccccatgtt gtgcaaaaaa gcggttagct ccttcggtcc
6661 tccgatcgtt gtcagaagta agttggccgc agtgttatca ctcatggtta tggcagcact
6721 gcataattct cttactgtca tgccatccgt aagatgcttt tctgtgactg gtgagtactc
6781 aaccaagtca ttctgagaat agtgtatgcg gcgaccgagt tgctcttgcc cggcgtcaat
6841 acgggataat accgcgccac atagcagaac tttaaaagtg ctcatcattg gaaaacgttc
6901 ttcggggcga aaactctcaa ggatcttacc gctgttgaga tccagttcga tgtaacccac
6961 tcgtgcaccc aactgatctt cagcatcttt tactttcacc agcgtttctg ggtgagcaaa
7021 aacaggaagg caaaatgccg caaaaaaggg aataagggcg acacggaaat gttgaatact
7081 catactcttc ctttttcaat attattgaag catttatcag ggttattgtc tcatgagcgg
7141 atacatattt gaatgtattt agaaaaataa acaaataggg gttccgcgca catttccccg
7201 aaaagtgcca c
//

**Genbank sequence for tRNA HTCS vector**

```
LOCUS       tRNA_screening_p        7146 bp DNA     circular SYN 29-MAR-2023
DEFINITION  synthetic circular DNA
ACCESSION   .
VERSION     .
KEYWORDS    .
SOURCE      synthetic DNA construct
  ORGANISM  synthetic DNA construct
REFERENCE   1  (bases 1 to 7146)
  AUTHORS   .
  TITLE     Direct Submission
  JOURNAL   Exported Mar 29, 2023 from SnapGene 6.2.1
            https://www.snapgene.com
COMMENT     LOCUS       pcDNA3.1/Zeo(+)          5015 bp          DNA    SYN
            DEFINITION  pcDNA3.1/Zeo(+)
            ACCESSION
            KEYWORDS
            SOURCE
              ORGANISM  other sequences; artificial sequences; vectors.
FEATURES             Location/Qualifiers
     source          1..7146
                     /mol_type="other DNA"
                     /organism="synthetic DNA construct"
     promoter        36..435
                     /label=UbC promoter
                     /note="human ubiquitin C promoter"
     promoter        445..463
                     /label=T7 promoter
                     /note="promoter for bacteriophage T7 RNA polymerase"
     regulatory      491..500
                     /label=Kozak sequence
                     /note="vertebrate consensus sequence for strong initiation
                     of translation (Kozak, 1987)"
                     /regulatory_class="other"
     CDS             497..2146
                     /codon_start=1
                     /product="firefly luciferase reporter gene luc2 (Photinus
                     pyralis) and is designed for high expression and reduced
                     anomalous transcription. This sequence was engineered with
                     fewer consensus regulatory sequences and has been codon
                     optimized for mammalian expression."
                     /label=Fluc2 (From Promega pGL4.10[luc2] vector)
                     /note="synthetic luc2 version of the luciferase gene"
                     /translation="MEDAKNIKKGPAPFYPLEDGTAGEQLHKAMKRYALVPGTIAFTDA

HIEVDITYAEYFEMSVRLAEAMKRYGLNTNHRIVVCSENSLQFFMPVLGALFIGVAVAP

ANDIYNERELLNSMGISQPTVVFVSKKGLQKILNVQKKLPIIQKIIIMDSKTDYQGFQS

MYTFVTSHLPPGFNEYDFVPESFDRDKTIALIMNSSGSTGLPKGVALPHRTACVRFSHA
```

RDPIFGNQIIPDTAILSVVPFHHGFGMFTTLGYLICGFRVVLMYRFEEELFLRSLQDYK

IQSALLVPTLFSFFAKSTLIDKYDLSNLHEIASGGAPLSKEVGEAVAKRFHLPGIRQGY

GLTETTSAILITPEGDDKPGAVGKVVPFFEAKVVDLDTGKTLGVNQRGELCVRGPMIMS

GYVNNPEATNALIDKDGWLHSGDIAYWDEDEHFFIVDRLKSLIKYKGYQVAPAELESIL

LQHPNIFDAGVAGLPDDDAGELPAAVVVLEHGKTMTEKEIVDYVASQVTTAKKLRGGVV
FVDEVPKGLTGKLDARKIREILIKAKKGGKIAV"

```
     misc_feature    2162..2281
                     /label=SV40 poly(A) signal
     enhancer        2351..2730
                     /label=CMV immediate early enhancer
                     /note="human cytomegalovirus immediate early enhancer"
     promoter        2731..2934
                     /label=CMV promoter
                     /note="human cytomegalovirus (CMV) immediate early
                     promoter"
     promoter        2979..2997
                     /label=T7 promoter
                     /note="promoter for bacteriophage T7 RNA polymerase"
     regulatory      3033..3042
                     /label=Kozak sequence
                     /note="vertebrate consensus sequence for strong initiation
                     of translation (Kozak, 1987)"
                     /regulatory_class="other"
     CDS             3039..3556
                     /codon_start=1
                     /product="NanoLuc(R) luciferase"
                     /label=Nluc
                     /note="human codon-optimized"
                     /translation="MVFTLEDFVGDWRQTAGYNLDQVLEQGGVSSLFQNLGVSVTPIQR
```

IVLSGENGLKIDIHVIIPYEGLSGDQMGQIEKIFKVVYPVDDHHFKVILHYGTLVIDGV

TPNMIDYFGRPYEGIAVFDGKKITVTGTLWNGNKIIDERLINPDGSLLFRVTING*VTG
WRLCERILA"

```
     primer_bind     3447..3465
                     /label=NanoLueckFseq (121)
     misc_feature    3516..3518
                     /label=PTC
     misc_feature    3616..3747
                     /label=SV40 poly(A) signal
     5'UTR           3763..3835
                     /label=5' human Tyr-tRNA leader
                     /note="5' human Tyr-tRNA leader"
     misc_feature    3838..3843
                     /label=BbsI
     promoter        3868..3898
```

/label=lac UV5 promoter
/note="E. coli lac promoter with an ""up"" mutation"
CDS 3952..4611
/codon_start=1
/gene="cat"
/product="chloramphenicol acetyltransferase"
/label=CmR
/note="confers resistance to chloramphenicol"
/translation="MEKKITGYTTVDISQWHRKEHFEAFQSVAQCTYNQTVQLDITAFL

KTVKKNKHKFYPAFIHILARLMNAHPEFRMAMKDGELVIWDSVHPCYTVFHEQTETFSS

LWSEYHDDFRQFLHIYSQDVACYGENLAYFPKGFIENMFFVSANPWVSFTSFDLNVANM

DNFFAPVFTMGKYYTQGDKVLMPLAIQVHHAVCDGFHVGRMLNELQQYCDEWQGGA"
CDS 4953..5258
/codon_start=1
/gene="ccdB"
/product="CcdB, a bacterial toxin that poisons DNA gyrase"
/label=ccdB
/note="Plasmids containing the ccdB gene cannot be
propagated in standard E. coli strains."
/translation="MQFKVYTYKRESRYRLFVDVQSDIIDTPGRRMVIPLASARLLSDK

VSRELYPVVHIGDESWRMMTTDMASVPVSVIGEEVADLSHRENDIKNAINLMFWGI"
misc_feature 5297..5302
/label=BbsI
variation 5305..5316
/label=3' term signal
/note="3' term signal"
rep_origin complement(5399..5987)
/direction=LEFT
/label=pBR322 ori
/note="high-copy-number ColE1/pMB1/pBR322/pUC origin of
replication"
CDS complement(6158..7018)
/codon_start=1
/gene="bla"
/product="beta-lactamase"
/label=AmpR
/note="confers resistance to ampicillin, carbenicillin, and
related antibiotics"
/translation="MSIQHFRVALIPFFAAFCLPVFAHPETLVKVKDAEDQLGARVGYI

ELDLNSGKILESFRPEERFPMMSTFKVLLCGAVLSRIDAGQEQLGRRIHYSQNDLVEYS

PVTEKHLTDGMTVRELCSAAITMSDNTAANLLLTTIGGPKELTAFLHNMGDHVTRLDRW

EPELNEAIPNDERDTTMPVAMATTLRKLLTGELLTLASRQQLIDWMEADKVAGPLLRSA

LPAGWFIADKSGAGERGSRGIIALGPDGKPSRIVVIYTTGSQATMDERNRQIAEIGAS

```
                     LIKHW"
     promoter        complement(7019..7123)
                     /gene="bla"
                     /label=AmpR promoter
ORIGIN
        1 ctgacgtcga cggatcggga gggttaatta cgcgtggcct ccgcgccggg ttttggcgcc
       61 tcccgcgggc gcccccctcc tcacggcgag cgctgccacg tcagacgaag ggcgcagcga
      121 gcgtcctgat ccttccgccc ggacgctcag gacagcggcc cgctgctcat aagactcggc
      181 cttagaaccc cagtatcagc agaaggacat tttaggacgg gacttgggtg actctagggc
      241 actggttttc tttccagaga gcggaacagg cgaggaaaag tagtcccttc tcggcgattc
      301 tgcggaggga tctccgtggg gcggtgaacg ccgatgatta tataaggacg cgccgggtgt
      361 ggcacagcta gttccgtcgc agccgggatt tgggtcgcgg ttcttgtttg tggatcgctg
      421 tgatcgtcac ttggttatcg aaattaatac gactcactat agggagaccc aagctgctag
      481 ccacgtggcc gccaccatgg aagatgccaa aaacattaag aagggcccag cgccattcta
      541 cccactcgag gacgggaccg ccggcgagca gctgcacaaa gccatgaagc gctacgccct
      601 ggtgcccggc accatcgcct ttaccgacgc acatatcgag gtggacatta cctacgccga
      661 gtacttcgag atgagcgttc ggctggcaga agctatgaag cgctatgggc tgaatacaaa
      721 ccatcggatc gtggtgtgca gcgagaatag cttgcagttc ttcatgcccg tgttgggtgc
      781 cctgttcatc ggtgtggctg tggccccagc taacgacatc tacaacgagc gcgagctgct
      841 gaacagcatg ggcatcagcc agcccaccgt cgtattcgtg agcaagaaag ggctgcaaaa
      901 gatcctcaac gtgcaaaaga agctaccgat catacaaaag atcatcatca tggatagcaa
      961 gaccgactac cagggcttcc aaagcatgta caccttcgtg acttcccatt tgccacccgg
     1021 cttcaacgag tacgacttcg tgcccgagag cttcgaccgg gacaaaacca tcgccctgat
     1081 catgaacagt agtggcagta ccggattgcc caagggcgta gccctaccgc accgcaccgc
     1141 ttgtgtccga ttcagtcatg cccgcgaccc catcttcggc aaccagatca tccccgacac
     1201 cgctatcctc agcgtggtgc catttcacca cggcttcggc atgttcacca cgctgggcta
     1261 cttgatctgc ggctttcggg tcgtgctcat gtaccgcttc gaggaggagc tattcttgcg
     1321 cagcttgcaa gactataaga ttcaatctgc cctgctggtg cccacactat ttagcttctt
     1381 cgctaagagc actctcatcg acaagtacga cctaagcaac ttgcacgaga tcgccagcgg
     1441 cggggcgccg ctcagcaagg aggtaggtga ggccgtggcc aaacgcttcc acctaccagg
     1501 catccgccag ggctacggcc tgacagaaac aaccagcgcc attctgatca cccccgaagg
     1561 ggacgacaag cctggcgcag taggcaaggt ggtgcccttc ttcgaggcta aggtggtgga
     1621 cttggacacc ggtaagacac tgggtgtgaa ccagcgcggc gagctgtgcg tccgtggccc
     1681 catgatcatg agcggctacg ttaacaaccc cgaggctaca aacgctctca tcgacaagga
     1741 cggctggctg cacagcggcg acatcgccta ctgggacgag gacgagcact tcttcatcgt
     1801 ggaccggctg aagagcctga tcaaatacaa gggctaccag gtagccccag ccgaactgga
     1861 gagcatcctg ctgcaacacc ccaacatctt cgacgccggg gtcgccggcc tgcccgacga
     1921 cgatgccggc gagctgcccg ccgcagtcgt cgtgctggaa cacggtaaaa ccatgaccga
     1981 gaaggagatc gtggactatg tggccagcca ggttacaacc gccaagaagc tgcgcggtgg
     2041 tgttgtgttc gtggacgagg tgcctaaagg actgaccggc aagttggacg cccgcaagat
     2101 ccgcgagatt ctcattaagg ccaagaaggg cggcaagatc gccgtgtaat gatagggtac
     2161 cttgtttatt gcagcttata atggttacaa ataaagcaat agcatcacaa atttcacaaa
     2221 taaagcattt ttttcactgc attctagttg tggtttgtcc aaactcatca atgtatctta
     2281 tgcatgaaga atctgcttag ggttaggcgt tttgcgctgc ttcgcgatgt acgggccaga
     2341 tatacgcgtt gacattgatt attgactagt tattaatagt aatcaattac ggggtcatta
     2401 gttcatagcc catatatgga gttccgcgtt acataactta cggtaaatgg cccgcctggc
     2461 tgaccgccca acgacccccg cccattgacg tcaataatga cgtatgttcc catagtaacg
     2521 ccaataggga ctttccattg acgtcaatgg gtggagtatt tacggtaaac tgcccacttg
     2581 gcagtacatc aagtgtatca tatgccaagt acgcccccta ttgacgtcaa tgacggtaaa
     2641 tggcccgcct ggcattatgc ccagtacatg accttatggg actttcctac ttggcagtac
     2701 atctacgtat tagtcatcgc tattaccatg gtgatgcggt tttggcagta catcaatggg
```

2761 cgtggatagc ggtttgactc acggggattt ccaagtctcc accccattga cgtcaatggg
2821 agtttgtttt ggcaccaaaa tcaacgggac tttccaaaat gtcgtaacaa ctccgcccca
2881 ttgacgcaaa tgggcggtag gcgtgtacgg tgggaggtct atataagcag agctctctgg
2941 ctaactagag aacccactgc ttactggctt atcgaaatta atacgactca ctatagggag
3001 acccaagctg gctagcgttt aaacttaagc ttgccaccat ggtattcaca ctcgaagatt
3061 tcgttgggga ctggcgacag acagccggct acaacctgga ccaagtcctt gaacagggag
3121 gtgtgtccag tttgtttcag aatctcgggg tgtccgtaac tccgatccaa aggattgtcc
3181 tgagcggtga aaatgggctg aagatcgaca tccatgtcat catcccgtat gaaggtctga
3241 gcggcgacca aatgggccag atcgaaaaaa tttttaaggt ggtgtaccct gtggatgatc
3301 atcactttaa ggtgatcctg cactatggca cactggtaat cgacggggtt acgccgaaca
3361 tgatcgacta tttcggacgg ccgtatgaag gcatcgccgt gttcgacggc aaaaagatca
3421 ctgtaacagg gaccctgtgg aacggcaaca aaattatcga cgagcgcctg atcaaccccg
3481 acggctccct gctgttccga gtaaccatca acggatgagt gaccggctgg cggctgtgcg
3541 aacgcattct ggcgtaagaa ttctaagggc gaattctgca gatatccagc acagtggcgg
3601 ccgctcgagt ctagattgtt tattgcagct tataatggtt acaaataaag caatagcatc
3661 acaaatttca caaataaagc atttttttca ctgcattcta gttgtggttt gtccaaactc
3721 atcaatgtat cttatcatgt ctggatcgct agagggcctt cctaatacga ctcactatag
3781 agcgctccgg tttttctgtg ctgaacctca ggggacgccg acacacgtac acgtctagtc
3841 ttcgcggccg cattaggcac cccaggcttt acactttatg cttccggctc gtataatgtg
3901 tggattttga gttaggatcc gtcgagattt tcaggagcta aggaagctaa aatggagaaa
3961 aaaatcactg gatataccac cgttgatata tcccaatggc atcgtaaaga acattttgag
4021 gcatttcagt cagttgctca atgtacctat aaccagaccg ttcagctgga tattacggcc
4081 tttttaaaga ccgtaaagaa aaataagcac aagttttatc cggcctttat tcacattctt
4141 gcccgcctga tgaatgctca tccggaattc cgtatggcaa tgaaagacgg tgagctggtg
4201 atatgggata gtgttcaccc ttgttacacc gttttccatg agcaaactga aacgttttca
4261 tcgctctgga gtgaatacca cgacgatttc cggcagtttc tacacatata ttcgcaagat
4321 gtggcgtgtt acggtgaaaa cctggcctat ttccctaaag ggtttattga gaatatgttt
4381 ttcgtctcag ccaatccctg ggtgagtttc accagttttg atttaaacgt ggccaatatg
4441 gacaacttct tcgcccccgt tttcaccatg ggcaaatatt atacgcaagg cgacaaggtg
4501 ctgatgccgc tggcgattca ggttcatcat gccgtttgtg atggcttcca tgtcggcaga
4561 atgcttaatg aattacaaca gtactgcgat gagtggcagg gcggggcgta aagatctgga
4621 tccggcttac taaaagccag ataacagtat gcgtatttgc gcgctgattt ttgcggtata
4681 agaatatata ctgatatgta tacccgaagt atgtcaaaaa gaggtatgct atgaagcagc
4741 gtattacagt gacagttgac agcgacagct atcagttgct caaggcatat atgatgtcaa
4801 tatctccggt ctggtaagca caaccatgca gaatgaagcc cgtcgtctgc gtgccgaacg
4861 ctggaaagcg gaaaatcagg aagggatggc tgaggtcgcc cggtttattg aaatgaacgg
4921 ctcttttgct gacgagaaca ggggctggtg aaatgcagtt taaggtttac acctataaaa
4981 gagagagccg ttatcgtctg tttgtggatg tacagagtga tattattgac acgcccgggc
5041 gacggatggt gatccccctg gccagtgcac gtctgctgtc agataaagtc tcccgtgaac
5101 tttacccggt ggtgcatatc ggggatgaaa gctggcgcat gatgaccacc gatatggcca
5161 gtgtgcccgt ctccgttatc ggggaagaag tggctgatct cagccaccgc gaaaatgaca
5221 tcaaaaacgc cattaacctg atgttctggg gaatataaat gtcaggctcc cttatacaca
5281 gccagtctgc agggaagaag accggtcctt tttttgcttt agtgagggtt aattcaggca
5341 tgtgagcaaa aggccagcaa aaggccagga accgtaaaaa ggccgcgttg ctggcgtttt
5401 tccataggct ccgcccccct gacgagcatc acaaaaatcg acgctcaagt cagaggtggc
5461 gaaacccgac aggactataa agataccagg cgtttccccc tggaagctcc ctcgtgcgct
5521 ctcctgttcc gaccctgccg cttaccggat acctgtccgc ctttctccct tcgggaagcg
5581 tggcgctttc tcaatgctca cgctgtaggt atctcagttc ggtgtaggtc gttcgctcca
5641 agctgggctg tgtgcacgaa ccccccgttc agcccgaccg ctgcgcctta tccggtaact
5701 atcgtcttga gtccaacccg gtaagacacg acttatcgcc actggcagca gccactggta
5761 acaggattag cagagcgagg tatgtaggcg gtgctacaga gttcttgaag tggtggccta

5821 actacggcta cactagaagg acagtatttg gtatctgcgc tctgctgaag ccagttacct
5881 tcggaaaaag agttggtagc tcttgatccg gcaaacaaac caccgctacc agcggtggtt
5941 tttttgtttg caagcagcag attacgcgca gaaaaaaagg atctcaagaa gatcctttga
6001 tcttttctac ggggtctgac gctcagtgga acgaaaactc acgttaaggg attttggtca
6061 tgagattatc aaaaaggatc ttcacctaga tccttttaaa ttaaaaatga agttttaaat
6121 caatctaaag tatatatgag taaacttggt ctgacagtta ccaatgctta atcagtgagg
6181 cacctatctc agcgatctgt ctatttcgtt catccatagt tgcctgactc cccgtcgtgt
6241 agataactac gatacgggag ggcttaccat ctggccccag tgctgcaatg ataccgcgag
6301 acccacgctc accggctcca gatttatcag caataaacca gccagccgga agggccgagc
6361 gcagaagtgg tcctgcaact ttatccgcct ccatccagtc tattaattgt tgccgggaag
6421 ctagagtaag tagttcgcca gttaatagtt tgcgcaacgt tgttgccatt gctacaggca
6481 tcgtggtgtc acgctcgtcg tttggtatgg cttcattcag ctccggttcc caacgatcaa
6541 ggcgagttac atgatccccc atgttgtgca aaaaagcggt tagctccttc ggtcctccga
6601 tcgttgtcag aagtaagttg gccgcagtgt tatcactcat ggttatggca gcactgcata
6661 attctcttac tgtcatgcca tccgtaagat gcttttctgt gactggtgag tactcaacca
6721 agtcattctg agaatagtgt atgcggcgac cgagttgctc ttgcccggcg tcaatacggg
6781 ataataccgc gccacatagc agaactttaa aagtgctcat cattggaaaa cgttcttcgg
6841 ggcgaaaact ctcaaggatc ttaccgctgt tgagatccag ttcgatgtaa cccactcgtg
6901 cacccaactg atcttcagca tcttttactt tcaccagcgt ttctgggtga gcaaaaacag
6961 gaaggcaaaa tgccgcaaaa aagggaataa gggcgacacg gaaatgttga atactcatac
7021 tcttcctttt tcaatattat tgaagcattt atcagggtta ttgtctcatg agcggataca
7081 tatttgaatg tatttagaaa aataaacaaa taggggttcc gcgcacattt ccccgaaaag
7141 tgccac
//

**Genbank sequence for sfGFP-TGA-150 pcDNA3.1 Hygro(+)**

```
LOCUS       sfGFP-TGA-150_pc        6322 bp DNA     circular SYN 29-MAR-2023
DEFINITION  pcDNA3.1/Hygro(+).
ACCESSION   .
VERSION     .
KEYWORDS    .
SOURCE      synthetic DNA construct
  ORGANISM  synthetic DNA construct
REFERENCE   1  (bases 1 to 6322)
  AUTHORS   Lueck
  TITLE     Direct Submission
  JOURNAL   Exported Mar 29, 2023 from SnapGene 6.2.1
            https://www.snapgene.com
FEATURES             Location/Qualifiers
     source          1..6322
                     /mol_type="other DNA"
                     /organism="pcDNA3.1/Hygro(+)"
     enhancer        235..614
                     /label=CMV enhancer
                     /note="human cytomegalovirus immediate early enhancer"
     promoter        615..818
                     /label=CMV promoter
                     /note="human cytomegalovirus (CMV) immediate early
                     promoter"
     promoter        863..881
                     /label=T7 promoter
                     /note="promoter for bacteriophage T7 RNA polymerase"
     regulatory      903..912
                     /label=Kozak sequence
                     /note="vertebrate consensus sequence for strong initiation
                     of translation (Kozak, 1987)"
                     /regulatory_class="other"
     CDS             909..1625
                     /codon_start=1
                     /product="GFP variant that folds robustly even when fused
                     to poorly folded proteins (Pedelacq et al., 2006)"
                     /label=superfolder GFP
                     /note="mammalian codon-optimized"
                     /translation="MVSKGEELFTGVVPILVELDGDVNGHKFSVRGEGEGDATNGKLTL

KFICTTGKLPVPWPTLVTTLTYGVQCFSRYPDHMKRHDFFKSAMPEGYVQERTISFKDD

GTYKTRAEVKFEGDTLVNRIELKGIDFKEDGNILGHKLEYNFNSH*VYITADKQKNGIK

ANFKIRHNVEDGSVQLADHYQQNTPIGDGPVLLPDNHYLSTQSVLSKDPNEKRDHMVLL
                     EFVTAAGITHGMDELYK"
     misc_feature    1356..1358
                     /label=TGA150
     CDS             1635..1658
                     /codon_start=1
```

```
                     /product="peptide that binds Strep-Tactin(R), an engineered
                     form of streptavidin"
                     /label=Strep-Tag II
                     /translation="WSHPQFEK"
     CDS             1662..1685
                     /codon_start=1
                     /product="8xHis affinity tag"
                     /label=8xHis
                     /translation="HHHHHHHH"
     CDS             1701..1724
                     /codon_start=1
                     /product="peptide that binds Strep-Tactin(R), an engineered
                     form of streptavidin"
                     /label=Strep-Tag II
                     /translation="WSHPQFEK"
     polyA_signal    1753..1977
                     /label=bGH poly(A) signal
                     /note="bovine growth hormone polyadenylation signal"
     rep_origin      2023..2451
                     /direction=RIGHT
                     /label=f1 ori
                     /note="f1 bacteriophage origin of replication; arrow
                     indicates direction of (+) strand synthesis"
     promoter        2465..2794
                     /label=SV40 promoter
                     /note="SV40 enhancer and early promoter"
     rep_origin      2645..2780
                     /label=SV40 ori
                     /note="SV40 origin of replication"
     CDS             2843..3868
                     /codon_start=1
                     /gene="aph(4)-Ia"
                     /product="aminoglycoside phosphotransferase from E. coli"
                     /label=HygR
                     /note="confers resistance to hygromycin"
                     /translation="MKKPELTATSVEKFLIEKFDSVSDLMQLSEGEESRAFSFDVGGRG

YVLRVNSCADGFYKDRYVYRHFASAALPIPEVLDIGEFSESLTYCISRRAQGVTLQDLP

ETELPAVLQPVAEAMDAIAAADLSQTSGFGPFGPQGIGQYTTWRDFICAIADPHVYHWQ

TVMDDTVSASVAQALDELMLWAEDCPEVRHLVHADFGSNNVLTDNGRITAVIDWSEAMF

GDSQYEVANIFFWRPWLACMEQQTRYFERRHPELAGSPRLRAYMLRIGLDQLYQSLVDG

NFDDAAWAQGRCDAIVRSGAGTVGRTQIARRSAAVWTDGCVEVLADSGNRRPSTRPRAK
                     E"
     polyA_signal    3998..4119
                     /label=SV40 poly(A) signal
                     /note="SV40 polyadenylation signal"
     primer_bind     complement(4168..4184)
```

/label=M13 rev
/note="common sequencing primer, one of multiple similar variants"
protein_bind 4192..4208
/label=lac operator
/bound_moiety="lac repressor encoded by lacI"
/note="The lac repressor binds to the lac operator to inhibit transcription in E. coli. This inhibition can be relieved by adding lactose or isopropyl-beta-D-thiogalactopyranoside (IPTG)."
promoter complement(4216..4246)
/label=lac promoter
/note="promoter for the E. coli lac operon"
protein_bind 4261..4282
/label=CAP binding site
/bound_moiety="E. coli catabolite activator protein"
/note="CAP binding activates transcription in the presence of cAMP."
rep_origin complement(4570..5155)
/direction=LEFT
/label=ori
/note="high-copy-number ColE1/pMB1/pBR322/pUC origin of replication"
CDS complement(5326..6186)
/codon_start=1
/gene="bla"
/product="beta-lactamase"
/label=AmpR
/note="confers resistance to ampicillin, carbenicillin, and related antibiotics"
/translation="MSIQHFRVALIPFFAAFCLPVFAHPETLVKVKDAEDQLGARVGYI

ELDLNSGKILESFRPEERFPMMSTFKVLLCGAVLSRIDAGQEQLGRRIHYSQNDLVEYS

PVTEKHLTDGMTVRELCSAAITMSDNTAANLLLTTIGGPKELTAFLHNMGDHVTRLDRW

EPELNEAIPNDERDTTMPVAMATTLRKLLTGELLTLASRQQLIDWMEADKVAGPLLRSA

LPAGWFIADKSGAGERGSRGIIAALGPDGKPSRIVVIYTTGSQATMDERNRQIAEIGAS
LIKHW"
promoter complement(6187..6291)
/gene="bla"
/label=AmpR promoter
ORIGIN
1 gacggatcgg gagatctccc gatcccctat ggtgcactct cagtacaatc tgctctgatg
61 ccgcatagtt aagccagtat ctgctccctg cttgtgtgtt ggaggtcgct gagtagtgcg
121 cgagcaaaat ttaagctaca acaaggcaag gcttgaccga caattgcatg aagaatctgc
181 ttagggttag gcgttttgcg ctgcttcgcg atgtacgggc cagatatacg cgttgacatt
241 gattattgac tagttattaa tagtaatcaa ttacggggtc attagttcat agcccatata
301 tggagttccg cgttacataa cttacggtaa atggcccgcc tggctgaccg cccaacgacc
361 cccgcccatt gacgtcaata atgacgtatg ttcccatagt aacgccaata gggactttcc

421 attgacgtca atgggtggag tatttacggt aaactgccca cttggcagta catcaagtgt
481 atcatatgcc aagtacgccc cctattgacg tcaatgacgg taaatggccc gcctggcatt
541 atgcccagta catgaccttа tgggactttc ctacttggca gtacatctac gtattagtca
601 tcgctattac catggtgatg cggttttggc agtacatcaa tgggcgtgga tagcggtttg
661 actcacgggg atttccaagt ctccacccca ttgacgtcaa tgggagtttg ttttggcacc
721 aaaatcaacg ggactttcca aaatgtcgta acaactccgc cccattgacg caaatgggcg
781 gtaggcgtgt acggtgggag gtctatataa gcagagctct ctggctaact agagaaccca
841 ctgcttactg gcttatcgaa attaatacga ctcactatag ggacacccaa gctggctagg
901 ccgccaccat ggtttccaaa ggagaagagt tgttcacagg tgtcgttcct atccttgtcg
961 agctggacgg tgatgtgaat ggacacaagt ttagtgtcag gggggaagga gagggcgatg
1021 ctaccaacgg gaaactgacc ctgaaattca tttgtacaac gggtaagctg cctgttcctt
1081 ggcccacgct ggtcacgacg cttacctatg gcgtgcaatg cttcagtcgc tatcccgatc
1141 atatgaaaag acatgacttt ttcaagtccg caatgccaga aggctacgtg caggagagaa
1201 caatcagctt taaggacgat ggcacgtaca aaactcgggc cgaggtcaag tttgagggag
1261 acacattggt aaacagaatt gagctgaagg ggatcgactt taaggaagac ggtaatattc
1321 tcggacacaa gttggagtat aacttcaatt cacactgagt gtatataact gccgataagc
1381 aaaaaaatgg tatcaaggct aattttaaaa tccggcacaa tgtagaagac ggctccgtgc
1441 aactggcgga tcactaccag cagaacaccc ccatcggcga tggtccagtt ttgctgcccg
1501 ataatcatta tctcagcacc cagagcgtgc tttctaaaga tccaaatgaa aagagggatc
1561 acatggtcct tttggagttt gttacggctg ccggaatcac ccacgggatg gacgagctct
1621 acaagggcgg aagttggagc catccgcagt ttgaaaaagc gcaccaccat caccatcatc
1681 atcactccgg gggcagtgca tggtcacacc ctcagtttga gaagtaatag tgaacccgct
1741 gatcagcctc gactgtgcct tctagttgcc agccatctgt tgtttgcccc tcccccgtgc
1801 cttccttgac cctggaaggt gccactccca ctgtcctttc ctaataaaat gaggaaattg
1861 catcgcattg tctgagtagg tgtcattcta ttctgggggg tggggtgggg caggacagca
1921 agggggagga ttgggaagac aatagcaggc atgctgggga tgcggtgggc tctatggctt
1981 ctgaggcgga aagaaccagc tggggctcta gggggtatcc ccacgcgccc tgtagcggcg
2041 cattaagcgc ggcgggtgtg gtggttacgc gcagcgtgac cgctacactt gccagcgccc
2101 tagcgcccgc tcctttcgct ttcttccctt cctttctcgc cacgttcgcc ggctttcccc
2161 gtcaagctct aaatcggggg ctccctttag ggttccgatt tagtgcttta cggcacctcg
2221 accccaaaaa acttgattag ggtgatggtt cacgtagtgg gccatcgccc tgatagacgg
2281 tttttcgccc tttgacgttg gagtccacgt tctttaatag tggactcttg ttccaaactg
2341 gaacaacact caaccctatc tcggtctatt cttttgattt ataagggatt ttgccgattt
2401 cggcctattg gttaaaaaat gagctgattt aacaaaaatt taacgcgaat taattctgtg
2461 gaatgtgtgt cagttagggt gtggaaagtc cccaggctcc ccagcaggca gaagtatgca
2521 aagcatgcat ctcaattagt cagcaaccag gtgtggaaag tccccaggct ccccagcagg
2581 cagaagtatg caaagcatgc atctcaatta gtcagcaacc atagtcccgc ccctaactcc
2641 gcccatcccg cccctaactc cgcccagttc cgcccattct ccgccccatg gctgactaat
2701 ttttttattt tatgcagagg ccgaggccgc ctctgcctct gagctattcc agaagtagtg
2761 aggaggcttt tttggaggcc taggcttttg caaaaagctc ccgggagctt gtatatccat
2821 tttcggatct gatcagcacg tgatgaaaaa gcctgaactc accgcgacgt ctgtcgagaa
2881 gtttctgatc gaaaagttcg acagcgtctc cgacctgatg cagctctcgg agggcgaaga
2941 atctcgtgct ttcagcttcg atgtaggagg gcgtggatat gtcctgcggg taaatagctg
3001 cgccgatggt ttctacaaag atcgttatgt ttatcggcac tttgcatcgg ccgcgctccc
3061 gattccggaa gtgcttgaca ttggggaatt cagcgagagc ctgacctatt gcatctcccg
3121 ccgtgcacag ggtgtcacgt tgcaagacct gcctgaaacc gaactgcccg ctgttctgca
3181 gccggtcgcg gaggccatgg atgcgatcgc tgcggccgat cttagccaga cgagcgggtt
3241 cggcccattc ggaccgcaag gaatcggtca atacactaca tggcgtgatt tcatatgcgc
3301 gattgctgat ccccatgtgt atcactggca aactgtgatg gacgacaccg tcagtgcgtc
3361 cgtcgcgcag gctctcgatg agctgatgct ttgggccgag gactgccccg aagtccggca
3421 cctcgtgcac gcggatttcg gctccaacaa tgtcctgacg gacaatggcc gcataacagc

3481 ggtcattgac tggagcgagg cgatgttcgg ggattcccaa tacgaggtcg ccaacatctt
3541 cttctggagg ccgtggttgg cttgtatgga gcagcagacg cgctacttcg agcggaggca
3601 tccggagctt gcaggatcgc cgcggctccg ggcgtatatg ctccgcattg gtcttgacca
3661 actctatcag agcttggttg acggcaattt cgatgatgca gcttgggcgc agggtcgatg
3721 cgacgcaatc gtccgatccg gagccgggac tgtcgggcgt acacaaatcg cccgcagaag
3781 cgcggccgtc tggaccgatg gctgtgtaga agtactcgcc gatagtggaa accgacgccc
3841 cagcactcgt ccgagggcaa aggaatagca cgtgctacga gatttcgatt ccaccgccgc
3901 cttctatgaa aggttgggct tcggaatcgt tttccgggac gccggctgga tgatcctcca
3961 gcgcggggat ctcatgctgg agttcttcgc ccaccccaac ttgtttattg cagcttataa
4021 tggttacaaa taaagcaata gcatcacaaa tttcacaaat aaagcatttt tttcactgca
4081 ttctagttgt ggtttgtcca aactcatcaa tgtatcttat catgtctgta taccgtcgac
4141 ctctagctag agcttggcgt aatcatggtc atagctgttt cctgtgtgaa attgttatcc
4201 gctcacaatt ccacacaaca tacgagccgg aagcataaag tgtaaagcct ggggtgccta
4261 atgagtgagc taactcacat taattgcgtt gcgctcactg cccgctttcc agtcgggaaa
4321 cctgtcgtgc cagctgcatt aatgaatcgg ccaacgcgcg gggagaggcg gtttgcgtat
4381 tgggcgctct tccgcttcct cgctcactga ctcgctgcgc tcggtcgttc ggctgcggcg
4441 agcggtatca gctcactcaa aggcggtaat acggttatcc acagaatcag gggataacgc
4501 aggaaagaac atgtgagcaa aaggccagca aaaggccagg aaccgtaaaa aggccgcgtt
4561 gctggcgttt ttccataggc tccgcccccc tgacgagcat cacaaaaatc gacgctcaag
4621 tcagaggtgg cgaaacccga caggactata aagataccag gcgtttcccc ctggaagctc
4681 cctcgtgcgc tctcctgttc cgaccctgcc gcttaccgga tacctgtccg cctttctccc
4741 ttcgggaagc gtggcgcttt ctcatagctc acgctgtagg tatctcagtt cggtgtaggt
4801 cgttcgctcc aagctgggct gtgtgcacga accccccgtt cagcccgacc gctgcgcctt
4861 atccggtaac tatcgtcttg agtccaaccc ggtaagacac gacttatcgc cactggcagc
4921 agccactggt aacaggatta gcagagcgag gtatgtaggc ggtgctacag agttcttgaa
4981 gtggtggcct aactacggct acactagaag aacagtattt ggtatctgcg ctctgctgaa
5041 gccagttacc ttcggaaaaa gagttggtag ctcttgatcc ggcaaacaaa ccaccgctgg
5101 tagcggtttt tttgtttgca agcagcagat tacgcgcaga aaaaaaggat ctcaagaaga
5161 tcctttgatc ttttctacgg ggtctgacgc tcagtggaac gaaaactcac gttaagggat
5221 tttggtcatg agattatcaa aaaggatctt cacctagatc cttttaaatt aaaaatgaag
5281 ttttaaatca atctaaagta tatatgagta aacttggtct gacagttacc aatgcttaat
5341 cagtgaggca cctatctcag cgatctgtct atttcgttca tccatagttg cctgactccc
5401 cgtcgtgtag ataactacga tacgggaggg cttaccatct ggccccagtg ctgcaatgat
5461 accgcgagac ccacgctcac cggctccaga tttatcagca ataaaccagc cagccggaag
5521 ggccgagcgc agaagtggtc ctgcaacttt atccgcctcc atccagtcta ttaattgttg
5581 ccgggaagct agagtaagta gttcgccagt taatagtttg cgcaacgttg ttgccattgc
5641 tacaggcatc gtggtgtcac gctcgtcgtt tggtatggct tcattcagct ccggttccca
5701 acgatcaagg cgagttacat gatcccccat gttgtgcaaa aaagcggtta gctccttcgg
5761 tcctccgatc gttgtcagaa gtaagttggc cgcagtgtta tcactcatgg ttatggcagc
5821 actgcataat tctcttactg tcatgccatc cgtaagatgc ttttctgtga ctggtgagta
5881 ctcaaccaag tcattctgag aatagtgtat gcggcgaccg agttgctctt gcccggcgtc
5941 aatacgggat aataccgcgc cacatagcag aactttaaaa gtgctcatca ttggaaaacg
6001 ttcttcgggg cgaaaactct caaggatctt accgctgttg agatccagtt cgatgtaacc
6061 cactcgtgca cccaactgat cttcagcatc ttttactttc accagcgttt ctgggtgagc
6121 aaaaacagga aggcaaaatg ccgcaaaaaa gggaataagg gcgacacgga aatgttgaat
6181 actcatactc ttcctttttc aatattattg aagcatttat cagggttatt gtctcatgag
6241 cggatacata tttgaatgta tttagaaaaa taaacaaata ggggttccgc gcacatttcc
6301 ccgaaaagtg ccacctgacg tc

//
